# Cell-based high-content approach for SARS-CoV-2 neutralization identifies unique monoclonal antibodies and PI3K pathway inhibitors

**DOI:** 10.1101/2024.10.04.616743

**Authors:** Carly R. Cabel, Briana A. Guzman, Elaheh Alizadeh, Shuaizhi Li, Cameron Holberg, Chonlarat Wichaidit, Darren A. Cusanovich, Andrew L. Paek, Gregory R. J. Thatcher, Koenraad Van Doorslaer, Rachel S. Nargi, Rachel E. Sutton, Naveenchandra Suryadevara, James E. Crowe, Robert H. Carnahan, Samuel K. Campos, Curtis A. Thorne

## Abstract

The sudden rise of the SARS-CoV-2 virus and the delay in the development of effective therapeutics to mitigate it made evident a need for ways to screen for compounds that can block infection and prevent further pathogenesis and spread. Yet, identifying effective drugs efficacious against viral infection and replication with minimal toxicity for the patient can be difficult. Monoclonal antibodies were shown to be effective, yet as the SARS-CoV-2 mutated, these antibodies became ineffective. Small molecule antivirals were identified using pseudovirus constructs to recapitulate infection in non-human cells, such as Vero E6 cells. However, the impact was limited due to poor translation of these compounds in the clinical setting. This is partly due to the lack of similarity of screening platforms to the in vivo physiology of the patient and partly because drugs effective in vitro showed dose-limiting toxicities. In this study, we performed two high-throughput screens in human lung adenocarcinoma cells with authentic SARS-CoV-2 virus to identify both monoclonal antibodies that neutralize the virus and clinically useful kinase inhibitors to block the virus and prioritize minimal host toxicity. Using high-content imaging combined with single-cell and multidimensional analysis, we identified antibodies and kinase inhibitors that reduce virus infection without affecting the host. Our screening technique uncovered novel antibodies and overlooked kinase inhibitors (i.e. PIK3i, mTORi, multiple RTKi) that could be effective against SARS-CoV-2 virus. Further characterization of these molecules will streamline the repurposing of compounds for the treatment of future pandemics and uncover novel mechanisms viruses use to hijack and infect host cells.

## INTRODUCTION

The novel coronavirus SARS-CoV-2 initiated the COVID-19 pandemic, spread rapidly, and caused over 6 million deaths and long-term side effects in many who recovered from infection (1). SARS-CoV-2 is the third of the SARS-related viruses that have caused a global pandemic. The first viral pandemic was in 2003 with SARS-CoV, which originated in the Guangdong Province in China in late 2002 (2). The second is Middle East Respiratory Syndrome (MERS), reported in Saudi Arabia in 2012 (3). As SARS-CoV-2 continues to evolve into new strains and cause disease, it will be a public health challenge for years to come (4–6). There is a need to develop a variety of therapeutics that can target and clear infection safely. Previous work has used SARS-CoV-2-positive patient samples to develop antibodies to neutralize the virus (7). While this approach is certainly effective, new variants of the virus arise, the epitopes targeted by the monoclonal antibodies evolve, and neutralizing activity attenuates (8, 9). Anti-viral small molecules target the function of essential viral proteins, but are not safe from evolving viruses (10). Finding specific, efficacious anti-viral small molecules has been a challenge and takes years of drug optimization (11–13). Repurposing clinically approved drugs, however, has some clear advantages to developing novel drugs. Repurposed drugs are already approved for use or in clinical trials, and come with human toxicity profiles (14, 15). Thus, trials to test efficacy against SARS-CoV-2 infection severity can be streamlined. Much work has been done around repurposing known FDA approved drugs to treat the virus (16), however, many of these drugs, while effective at clearing the virus, also have significant, dose-limiting toxicities in patients (17). Identifying clinical drugs with anti-viral activity without significant host toxicities is a major challenge.

Effective repurposed drugs have been hard to identify, partly because of the simplicity of the drug screening assays for which they are identified. Viral infection and replication assays that capture natural SARS-CoV-2 strain infection and the nuances of host cell viability and health are needed. Many reported high-throughput screens have used pseudovirus systems in non-physiological cells such as Vero cells (13–15) or performed as virtual screens (18). Ideally, high-throughput screening techniques would use live virus and human airway epithelium to measure a compound’s effect on live virus levels and account for epithelial cell health.

High-content image-based screening is ideal for capturing many cell-level measurements of infection and host cell health. Here, we performed two high-throughput in vitro screens with human lung adenocarcinoma cells and authentic SARS-CoV-2 virus to identify 1) neutralizing antibodies that effectively target and neutralize the virus and 2) kinase small molecule inhibitors that can be repurposed for use in the clinic against the virus. We identify a new way to screen for viral neutralization and discover monoclonal antibodies and kinase inhibitors (e.g., PI3K) that can block viral entry and replication. The approach we describe proposes a method of evaluating drug effectiveness against SARS-CoV-2 infection while also selecting for minimal host cell perturbation.

## RESULTS

### Development of High-Content SARS-CoV-2 infection assay

A number of screens for small molecule inhibitors of SARS-CoV-2 infection and replication have been performed, yet most have been simple, non-physiological in nature, limiting their impact (19–21). We choose to conduct an image-based screen of patient-derived monoclonal antibodies and small molecule kinase inhibitors in the epithelial lung adenocarcinoma, Calu-3, combined with automated high-content imaging to uncover novel viral replication mechanisms and identify potent and more specific therapeutic strategies (Figure 1). Our approach would allow us to work with true SARS-CoV-2 in human cells and capture viral replications information and small to large host toxicity effects. Caco-2 and Calu-3 cells have been used previously in SARS-CoV-2 research due to their expression of ACE2 receptor as well as the protease TMPRSS2, which make them permissible to infection (22–24). We found that Calu-3 cells had higher expression of ACE2 receptor and were chosen to move forward with the screens (Sup Fig 1). We used 384-well optically clear imaging plates and authentic SARS-CoV-2 (strain 2019 n-CoV/USA_WA1/2020) virus to perform this screen. We plated cells in a BSL-2 facility and immediately treated them with small molecules/antibodies before transferring to the BSL-3 facility and infecting them. Mock-infected wells served as our positive control for the desired outcome of infection prevention. Infected with vehicle treatment wells served as negative controls and displayed robust viral amplification. The infection and replication were allowed to proceed for 48 hours, then cells were fixed by full submersion in 4% PFA for 30 minutes. Plates were moved out of the BSL-3 facility and stained for host cell markers phalloidin (to mark epithelial cell cortex and actin cytoskeleton) and DAPI (to mark nuclei and morphology), as well as dsRNA (to mark infected cells with active, replicating virus). dsRNA is an intermediate product during SARS-CoV-2 viral replication and is a robust marker for infected cells in Calu-3 cells (25–27). Combining these three cell stains allowed us to observe how compounds affect viral infection while monitoring effects on host cell number and morphology. To validate our assay, we performed two large-scale screens of neutralizing antibodies and kinase inhibitors to observe effects on virus replication and the host cell number and morphology.

**Figure 1:**
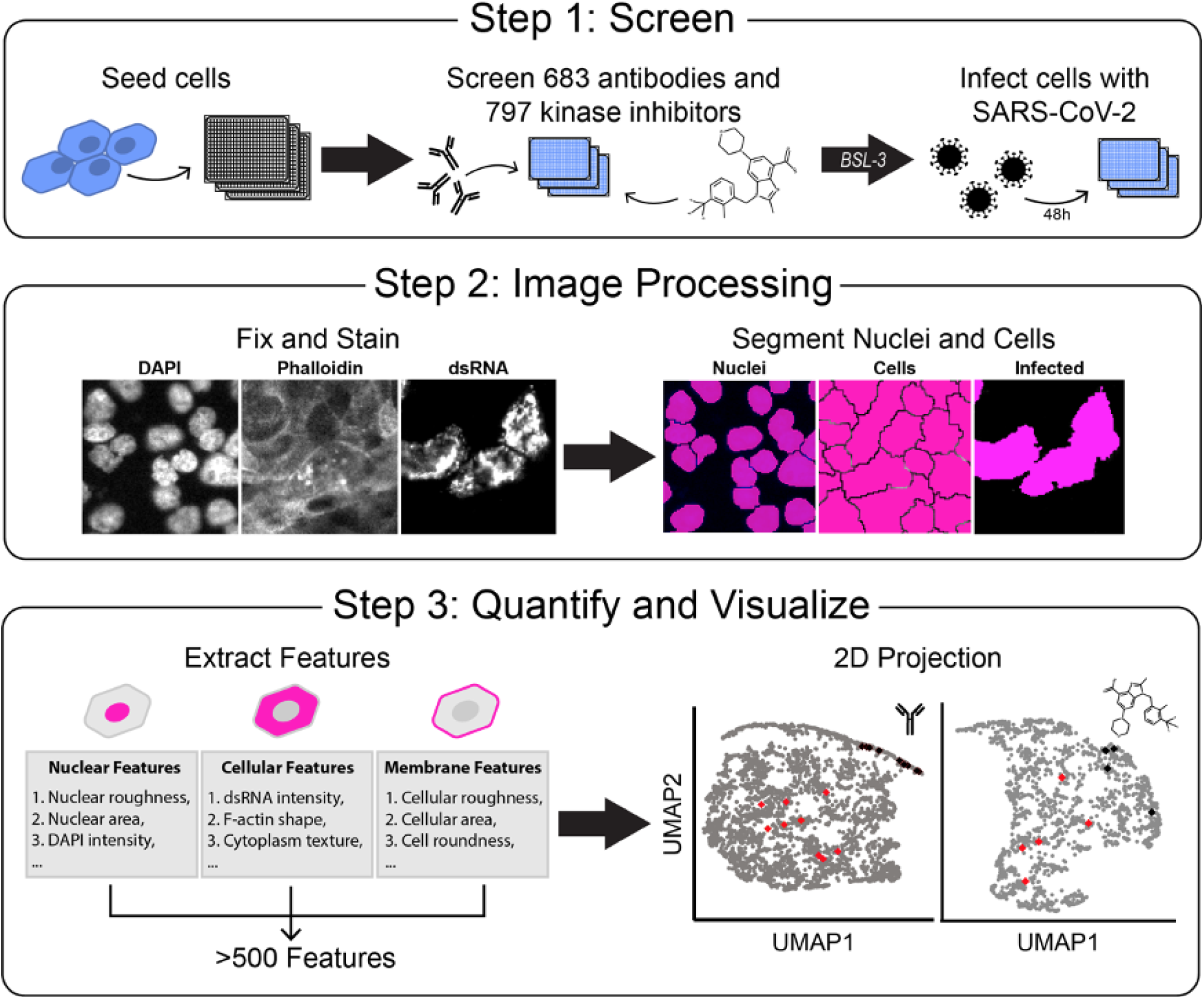
Development of High-throughput SARS-CoV-2 detection screen to identify inhibitors of viral infection. A. Overview of experimental design. Step 1: Human lung carcinoma cells are seeded in a 384-well microplate and incubated with treatment, and moved to a BSL-3 facility. Cells are then treated with authentic virus for 48 hours. Step 2: Plates are then fixed, stained for viral dsRNA, actin (phalloidin), and DNA (DAPI), and imaged by fluorescent microscopy and processed with segmentation software to identify single-cells objects. Step 3: Hundreds of cell features are computationally extracted, and data visualized in 2D UMAP projections.

The screening produced large image sets of cells stained for viral replication (dsRNA), actin cytoskeleton (phalloidin), and nuclear morphology (DAPI). We processed images with CellProfiler (28–30) to segment the images into single-cell objects and measure hundreds of cell-level features in three major classes: 1) Morphology, 2) Intensity, 3) Texture. These features were averaged per well and uniform manifold approximation and projection (UMAP) dimensionality reduction techniques were used to project cells into 2-dimensional space for evaluation (31, 32). Overall, this approach made for scalable, 384-well, automated imaging and analysis format for testing hundreds of compounds.

### Neutralizing antibodies reduce SARS-CoV-2 infection in Calu-3 cells

One particularly effective therapeutic strategy for SARS-CoV-2 infection is the delivery of neutralizing monoclonal antibodies (mAbs) developed against the SARS-CoV-2 spike protein that block viral entry. We tested the effects of 683 patient-derived antibodies to block infection of SARS-CoV-2 in human lung adenocarcinoma cells. These antibodies have been developed through B cell enrichment of four COVID-19 positive patients (7, 33, 34) and tested for binding affinity against the virus spike protein. They were previously classified into five classes representing their binding to the spike protein and whether they cross-reacted with SARS-CoV-1 (Fig. 2A): Class I antibodies show binding ability to the ectodomain of trimeric S2 (S2P_ecto_) and the RBD domain of SARS-CoV-2 alone; Class II antibodies bind to the S2P_ecto_ and the RBD domain of SARS-CoV-2 but cross-reacted with the S2P_ecto_ of SARS-CoV-1; Class III bind S2P_ecto_ and the NTD subdomain of SARS-CoV-2 only; Class IV bind S2P_ecto_ of SARS-CoV-2 only with no S1 domain activity; and Class V bind S2P_ecto_ of SARS-CoV-2 and cross-reacted with S2P_ecto_ of SARS-CoV-1. We also tested a Class VI of antibodies that were not shown to interact with spike protein in SARS viruses from binding assays (7), yet some from this class could diminish dsRNA signal Calu-3 cells. Their capacity to block authentic virus replication in cells was unknown except for two lead antibodies validated previously (35). We tested these antibodies in a dose response of four ten-fold dilutions, infected with an MOI of 1, for 48 hours to determine if antibodies were efficacious against authentic virus challenge. After fixing we stained for dsRNA as a measure of viral replication, phalloidin to measure f-actin of the host cells, DAPI to visualize the nuclei, and imaged using automated microscopy (Fig 2B). We first measured total dsRNA from each of the antibody conditions and plotted them as area under the curve graphs to observe which antibodies reduced viral burden. Of the antibodies that were tested, the ones in Class I had the most potent response against the virus (Fig 2C). The majority of the Class I antibodies reduced the dsRNA intensity to mock infection levels, while the other classes had fewer antibodies with such potency. Class VI had the largest number of positive hits, but also was the largest class of antibodies. Importantly, class VI antibodies were not identified in initial spike protein binding assays, yet we readily observed many apparent potent antibodies in this group. Class VI hits in our screen likely represent false negatives from the previous spike binding characterization or some other unknown mechanism of action. This demonstrates that our high-content and high-throughput assay effectively identifies potent SARS-CoV-2 neutralizing antibodies and could be more sensitive and/or specific than spike protein binding assays.

**Figure 2:**
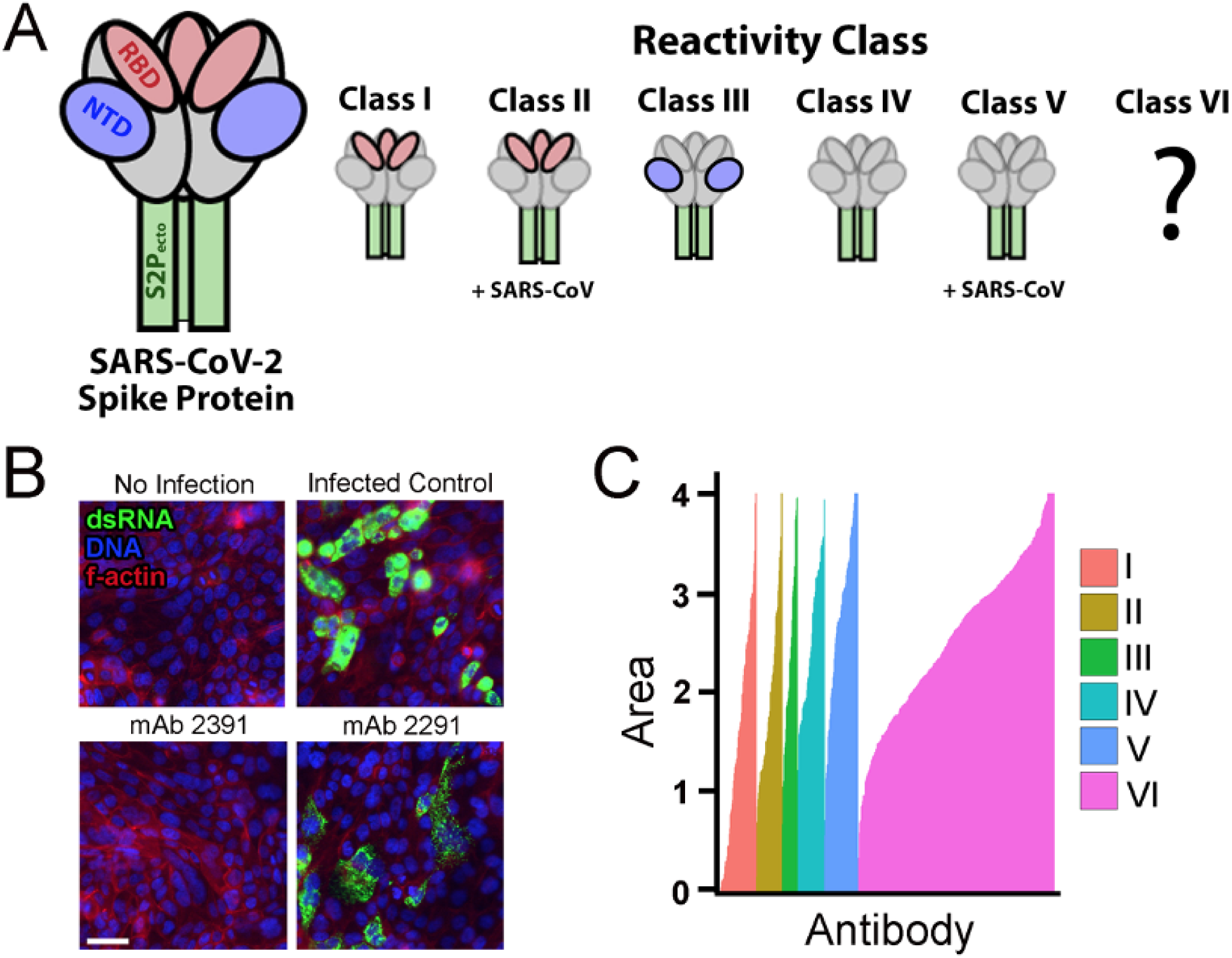
Neutralizing antibodies block virus in Calu-2 cells. A. Description of classes of putative binding properties of patient-derived antibodies. Classes I-V were as previously described. Class VI is an uncharacterized class of antibodies that were isolated from the same SARS-CoV-2 patients as Classes I-V but lacked activity in the secondary spike protein ELISA binding assay. B. Representative images from the antibody screen. No infection represents the Calu-3 cells with mock media, and no virus added. Monoclonal antibody (antibody) 2391 was a positive result, showing neutralization of the virus in cells. Antibody 2291 represents a less successful antibody, with clear detection of dsRNA from SARS-CoV-2. Nuclei are stained blue with DAPI. F-actin is stained with phalloidin in red. dsRNA is stained in green. Scale bar = 50µM. C. Cumulative dsRNA measurements for all antibody treatments. Colors represent the different reactivity classes of antibody targets against the virus. Area under the curve (AUC) is plotted for four increasing concentrations of antibodies. The control concentration has been normalized to a value of 1, making the max value of each dose also 1. AUC of the curve is the sum of all four antibody concentrations.

### Multidimensional Profiling Demonstrates that neutralizing antibodies reduce SARS-CoV-2 infection with minimal effect on phenotype of Calu-3 cells

Next, we wanted to understand how the phenotype of host cells is affected by antibody treatment and if capturing host morphology measurements enriches “hit” antibody selection. The ideal antibody neutralizes the virus leaving the host cells indistinguishable from uninfected control conditions. As described in Fig. 1, we measured over 500 features per cell and performed feature reduction to discard highly correlated measurements. This reduced the features measured down to 150 measurable features used in analysis. Because SARS-CoV-2 highly infected cells only make up about 1% of the Calu-3 cells under our culture conditions (Sup Fig. 2), we discarded all cells except those with the top 1% of dsRNA signal. Among this top 1%, we averaged the host cells features at well-level and created a multidimensional profile for each antibody treatment. We performed Uniform Manifold Approximation and Projection (UMAP) to reduce dimensionality and visualize in two dimensions (Fig. 3A). The 683 antibodies, in general, produced subtle effects on host cell morphology as might be expected from the fact that antibodies target very specific epitopes. Importantly, the mock controls clustered together (black diamonds, Fig. 3A), and the infected controls clustered together (red diamonds). The treatment conditions (gray dots) grouped towards either the infected controls, suggesting those antibodies had poor or no neutralizing activity and the virus distorted host cell morphology, or they grouped with the mock controls, suggesting robust viral neutralization and maintenance of host cell morphology. To understand if this was a concentration-dependent response, we plotted each of the conditions from low to high separately to observe how the antibodies influenced host cell morphology. Visualization of each of the four antibody concentrations separately shows a subtle movement in phenotype from high infection phenotype towards the uninfected phenotype (Sup Fig 3). This confirms that the antibodies do behave in a dose-dependent manner and improve both neutralizing capabilities while preserving host morphology as we increase concentration.

**Figure 3:**
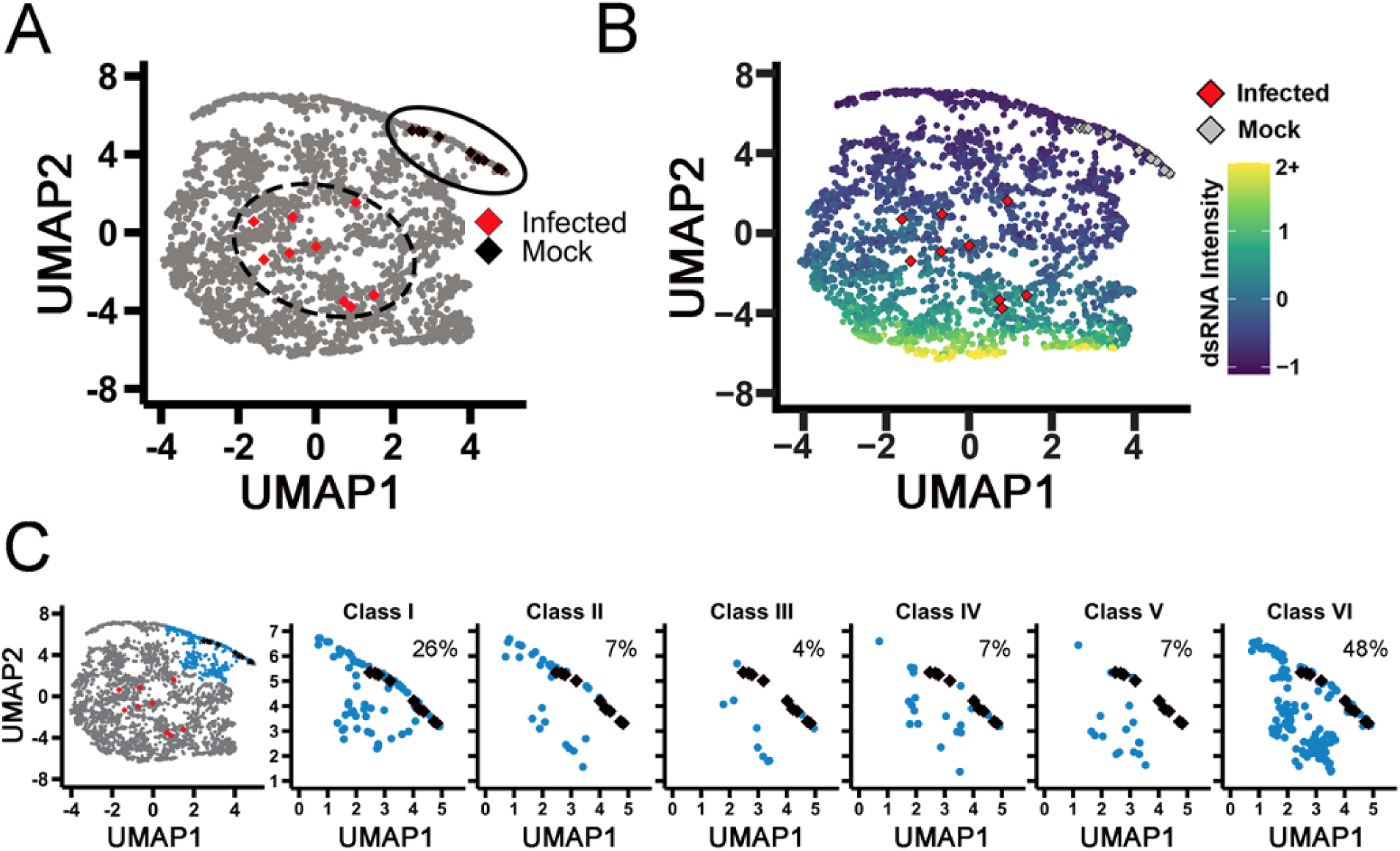
High-dimensional profiling show classes of mAbs that have varying neutralizing activity. A. UMAP of all antibodies. Black diamonds represent mock controls; infected controls are represented by red diamonds. Each point on the map represents a single antibody at a single dose. B. The UMAP in panel A recolored as increasing intensity of dsRNA. Infected controls are represented by red diamonds and mock controls are represented by gray diamonds. C. Low-virus cluster (blue) broken down by the identified classes. Each graph is labeled with % of class antibodies that had potent neutralizing. Classes I and VI had the highest percent of antibodies in the blue cluster compared to other classes.

To compare dsRNA levels, we colored the UMAP by median log_10_-transformed intensity of dsRNA signal (Fig 3B). We can clearly see a shift from the bottom to the top of the plot as the dsRNA signal decreases; the morphology shifts closer to the mock control (gray diamonds) and away from the infected control (red diamonds). To identify the top antibodies in our screen, we performed cluster analysis and identified a subgroup of antibodies that co-cluster with mock conditions, consistent with strong viral neutralization activity. When we broke this group down based on spike protein binding class, we see that class I has the highest hit rate of previously defined antibodies classes at 26% (Fig 3C), consistent with many studies showing the RBD domain is the most druggable part of the spike protein (7, 33, 36).

### Kinase inhibitors act to suppress SARS-CoV-2 infection in Calu-3 cells

Kinase inhibitors have been repurposed in the past to act as viral inhibitors for many types of viruses (14). Tyrosine kinase inhibitors have been identified to block SARS-CoV viruses in Vero E6 cells with minor toxicity to the host cells (20, 37). In order to test the efficacy of kinase inhibitors against naturally occurring SARS-CoV-2 virus, we tested 796 kinase inhibitors that span the human kinome in our high-content assay with the authentic virus in the same format as the antibody screen described above. We tested two doses, 100nM and 10µM, and infected for 48 hours with the same immunofluorescent markers (Fig 4A). When we measure mean dsRNA intensity per well as a measure, nearly half of the kinase inhibitor library showed some level of reduction in dsRNA staining (Fig. 4B). This is likely due to the fundamental role kinases play in cell signaling and cell physiology.

**Figure 4:**
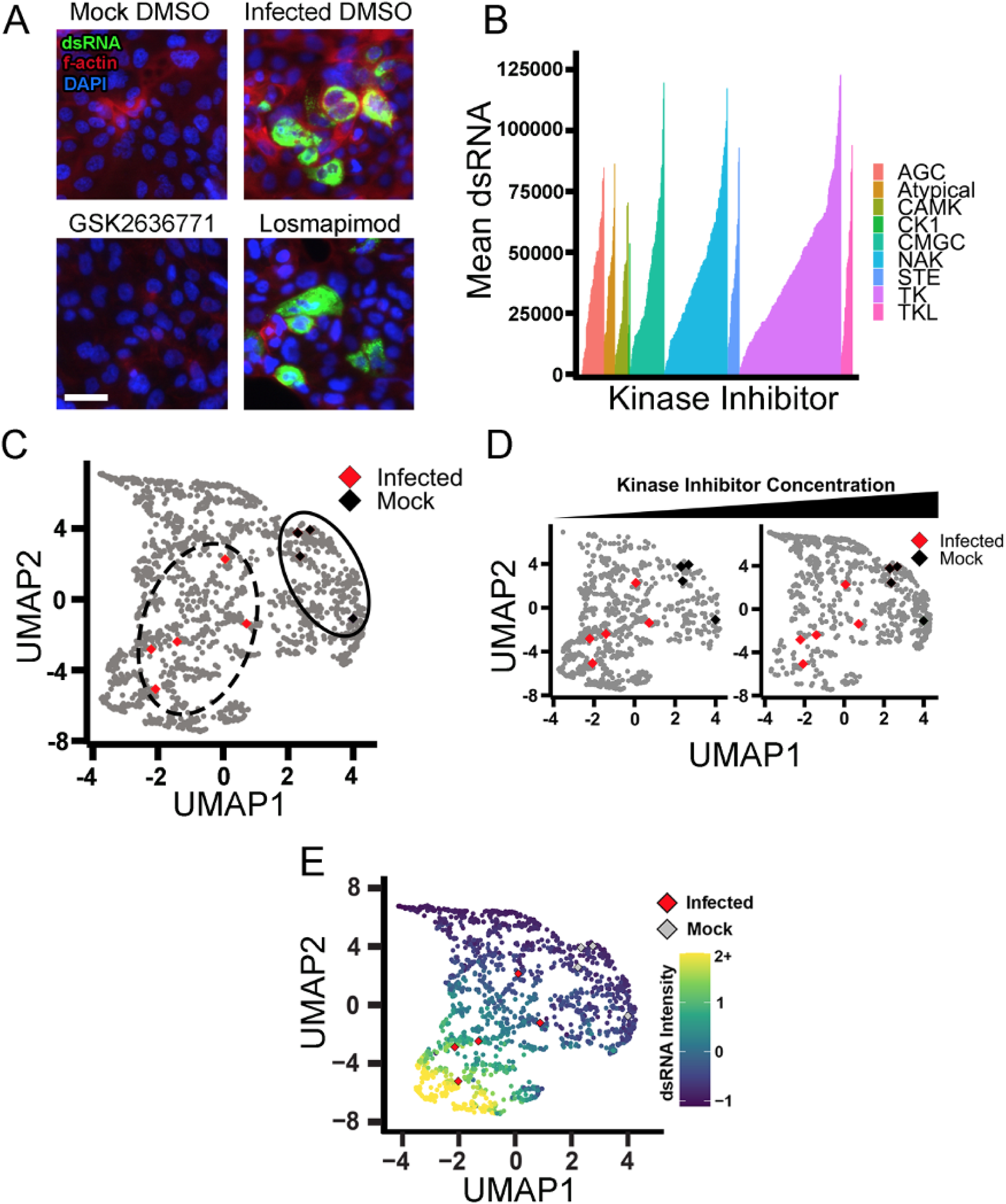
Kinase inhibitors from Tyrosine Kinase family actively inhibit viral infection. A. Representative images of mock and infected Calu-3 cells with DMSO control. Nuclei are stained in blue DAPI. F-actin is stained with phalloidin in red. dsRNA was stained in green showing active SARS-CoV-2 infection. Scale = 50µM. GSK26367871 represents a kinase inhibitor that reduced infection greatly. Losmapimod represents a kinase inhibitor with poor activity. B. Kinase inhibitors plotted as mean dsRNA, ranked from left to right by ability to reduce dsRNA. Colors are representative of the branch of the kinome tree they belong to. NAK = “not a kinase (inhibitor)” and represents compounds whose activity is poorly understood (e.g. flavanoids). C. UMAP of kinase inhibitors as compared to infected and mock controls. Each point represented a single kinase inhibitor at a single dose. Infected controls are represented by red diamonds and black diamonds represent mock controls. D. UMAP of kinase inhibitors separated by concentration. Left plot is 100 nM; right plot is 10µM. The shift of the density of points indicates that with an increasing concentration of kinase inhibitor, the cell phenotype moves closer to the mock control (black diamonds) and away from the infected control (red diamonds). E. The UMAP from (D) was recolored to show dsRNA intensity. Red diamonds represent infected controls, and gray diamonds represent mock controls.

Despite the promise of many kinase inhibitors reducing apparent viral replication as observed by reduced dsRNA, negative effects on host cells could be responsible for much of this, as kinases are critical for host cell function. To triage compounds that have a deleterious effect on the host cell, we again performed our high-content analysis described in Fig. 1. Multidimensional analysis showed that the infected and mock controls were distinct and that kinase inhibitors induced a spread of morphological changes (Fig. 4C). Additionally, we saw that movement from infected to mock with increased compound concentration, similar to what we saw with increasing antibody concentrations (Fig 4D). We then evaluated the UMAP with dsRNA median intensity. We confirmed that the cells with lower dsRNA clustered with the mock controls, while the higher dsRNA clustered with the infected control (Fig 4E).

### Targeting the PI3K pathway is a putative strategy for SARS-CoV-2 suppression

To evaluate the effect of the kinase inhibitors on the host cell health and morphology, we performed cluster analysis to identify groups that blocked viral inhibition while having minimal effects on cell morphology (Fig 5A). Seven groups of phenotypes were identified, with red cluster #1 representing drug treatment conditions where cells appear healthy and uninfected. When the top hits from single parameter analysis of dsRNA (Fig 4B) are plotted in UMAP space (Fig 5B), we found that though they reduced the dsRNA signal in cells, only a few showed a healthy host cell phenotype (as denoted by being in the red cluster). Next, we wanted to evaluate how the original two-concentration screen compared to a 12-concentration confirmation study. We hypothesized that an ideal inhibitor that blocks viral replication with minimal host toxicity would phenotypically move into the red group as the treatment concentration increased. Further, the kinase inhibitors blocking viral replication but causing host cell toxicity would not move into the red cluster but away to some other phenotypic cluster. In Figure 5 C-D and Sup Figure 4, we indicated the two doses of each inhibitor with a gray X. Then we showed low to high doses with an arrow to visualize the directionality of increasing compound concentration in UMAP space. Among the top 33 compounds, we observed that our image analysis could identify a wide variety of host cell changes, many of which would be undesirable. An example of this is the drug Ibrutinib, a BTK inhibitor (38–40), which showed a shift away from the red cluster with increasing concentration (Fig 5C). This suggests that although the inhibitor blocked viral infection, leading it to be a top hit, it likely affected the host cell in a toxic way. Indeed, when we challenged the cells with a dose response to block viral infection, it did block the virus in high concentrations but at the cost of significant cell toxicity. Low concentrations had little effect on the virus; at high concentrations, the decrease in dsRNA signal is likely due to a decrease in overall cell host cell viability.

**Figure 5:**
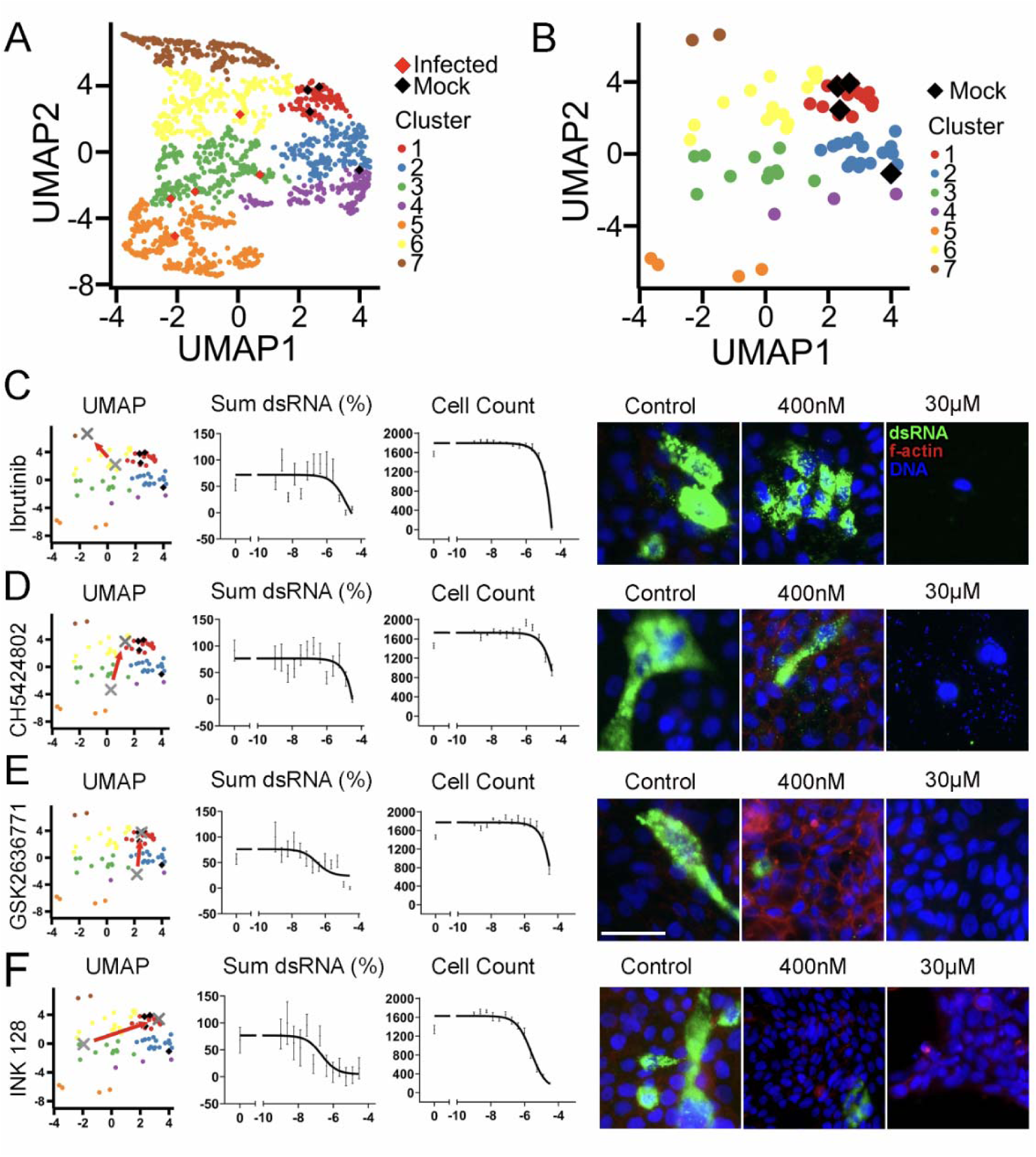
PI3K pathway identified as a strategy for SARS-CoV-2 suppression. A. Kinase inhibitor screen hits grouped by phenotypic clusters. The red cluster 1 groups closely with the mock controls (black diamonds) while the infected controls (red diamonds) expand upon several other clusters. B. The top hits from simple mean dsRNA measurement (Fig. 4B) were plotted alone in UMAP. Compounds in the red 1 cluster still closely resemble the mock controls, yet a wide spread of phenotypes is observed. C., D., E., F., The top compounds were retested in a 12-concentration dose-response curve. The gray X indicates the low and high doses from the initial screen. The red arrow points from the low dose to the high dose to emphasize the shift in location UMAP space. Sum dsRNA was normalized to percent of response to control. Cell count from dose response curve was measured by counting DAPI stained nuclei. Right side, representative images of the specified kinase inhibitor with low and high concentrations. Nuclei are stained with DAPI in blue, F-actin is stained with phalloidin in red, and dsRNA is stained in green. Scale bar = 50µM. Error bars represent SEM.

Another example of the reliability of our UMAP analysis is CH5424802, an ALK inhibitor (41, 42). While it reduced dsRNA very well and seemed to pass the cell viability test, looking at morphology, we see that it impacted the nuclei of the host cells by causing blebbing and general toxicity (Fig 5D). This was apparent in the shift in our UMAP data. The high concentration dose ended up in the adjacent yellow cluster, being outside of the target red cluster, showing that the inhibitor affected the host cell negatively.

In contrast to Ibrutinib and CH5424802 was GSK2636771, a PI3K inhibitor (43). GSK2636771 showed a shift closer to cluster 1 controls in the two-dose test as well as positive performance in the dose response (Fig 5E). It showed a reduction of dsRNA while having minimal effect on host cell viability and morphology. Unsurprisingly, INK128, an mTOR inhibitor, also performed well in our assay. Moving in greatly to cluster 1 and having little effect on host cell viability (Fig 5F.). We observed that for all 33 dsRNA hits, the ones that moved closer to cluster 1 generally exhibited lower toxicity and should be considered strong candidates for repurposing (Table 1, Sup Fig 4).

**Table 1:**
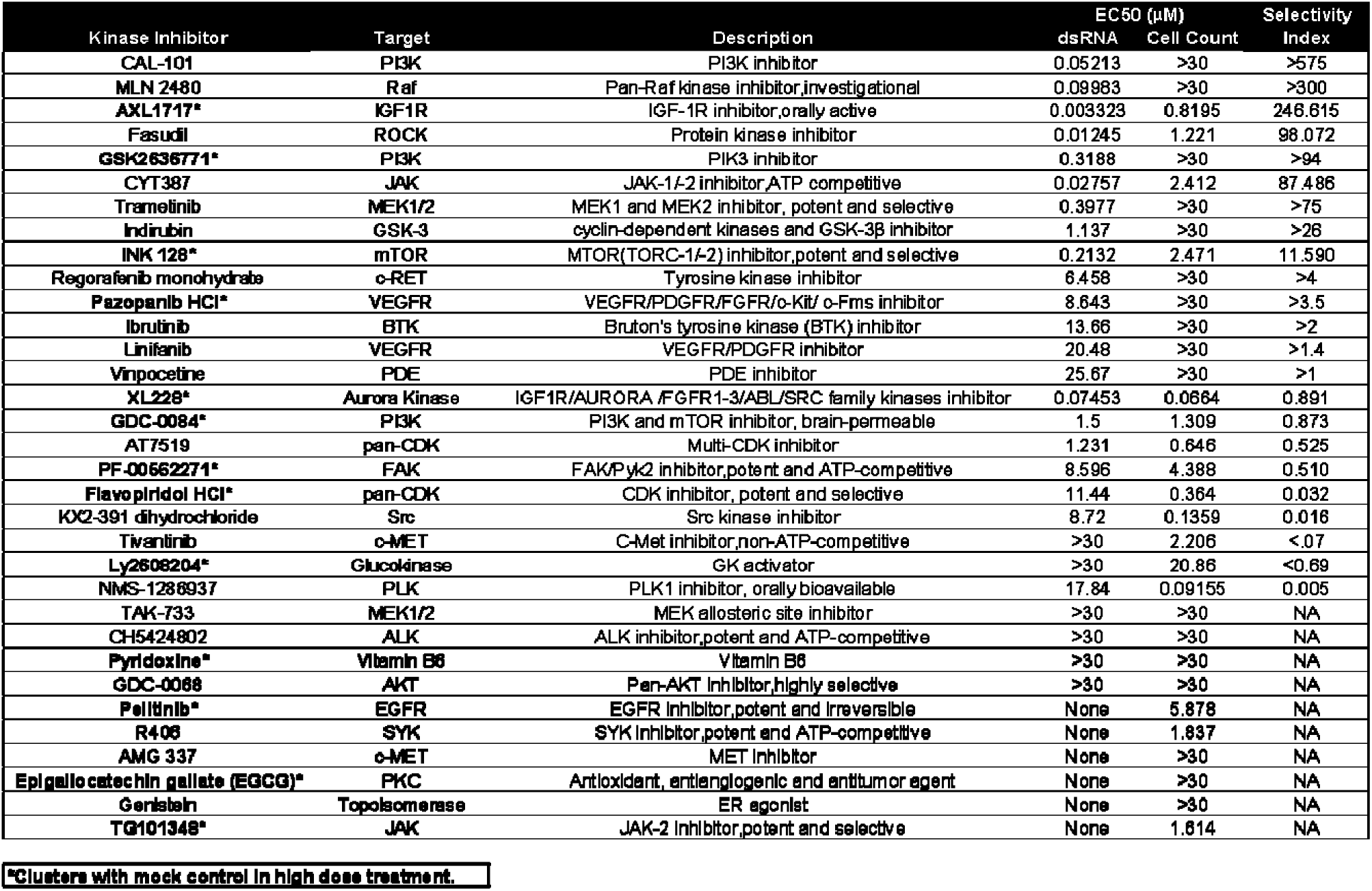
Top kinase inhibitors from Figure 5 were tested in a dose-response curve. EC_50_s for dsRNA and cell count are reported. Fold column shows cell count over dsRNA to support a therapeutic window of treatment may exist. Any compound in which the EC_50_ was above 30 μM is reported as >30. *Represent compounds that group with the red cluster from Figure 5 and represent the strongest candidates for in vivo validation.

We also wanted to test these top compounds in variants of the SARS-CoV-2 virus. SARS-CoV-2 has shown mutation capability and evading drug response, and we hypothesized that using small molecules, some identified in our screens, would still be effective across cell lines as well as viral strains. We tested three compounds in both A549 and Calu-3 lung cells: INK 128 and GSK2636771 which were identified in our screen, and Nirmatrelvir (a protease inhibitor) (44) . We performed a dose-response curve against two additional strains, Omicron BA.1 (B.1.1.529) and Delta (B.1.617.2)and saw similar responses of these small molecule inhibitors against the different strains (Sup Fig 5).

### Combination treatment of monoclonal antibodies with small molecule inhibitors increases the efficacy of neutralization

With the success of monoclonal antibodies in the clinic, and the promise of our small molecule inhibitor screen in identifying potential blockers of SARS-CoV-2, we hypothesized that combining both treatments together would lead to increased efficacy in neutralizing the virus. To test this, we performed combination treatments using a select few monoclonal antibodies, mAb 2355, mAb 2489, and mAb 2819, and Nirmatrelvir to test their synergistic effects against the SARS-Cov-2 WA strain (Fig 6A). Nirmatrelvir is a protease inhibitor compound that targets viral proteases that contribute to viral replication in host cells. This compound has been used in treatment for SARS-CoV-2 and its variants with success (45). Using A549 lung cells, we infected these cells with SARS-CoV-2 and saw under DMSO only conditions robust infection with virus as compared to mock control (Fig 6B).

**Fig 6:**
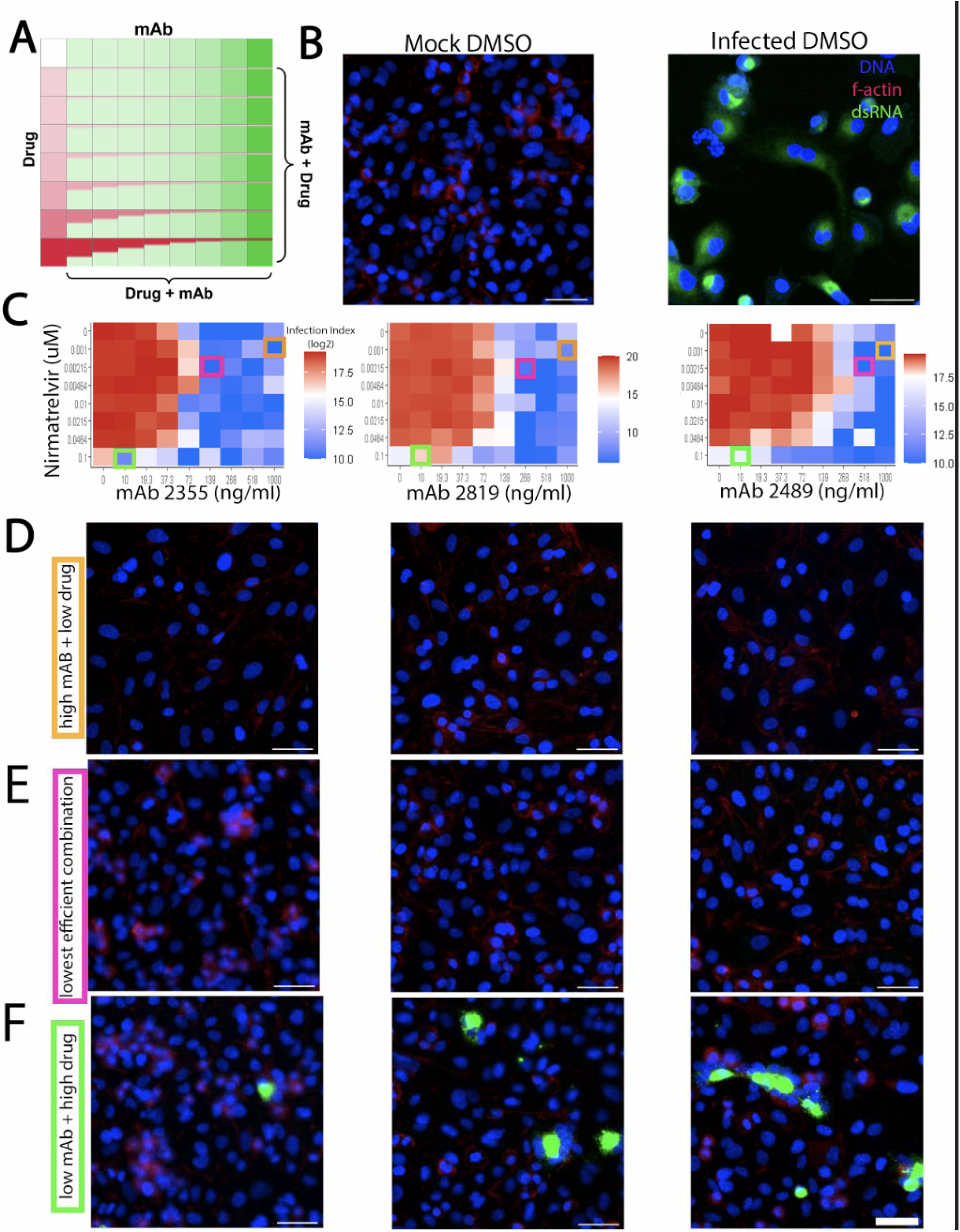
Effect of combination treatment of monoclonal antibodies with small molecule inhibitors on the efficacy of neutralization. A. Schematic of small molecule and antibody combinatorial treatment. Monoclonal antibodies were added in increasing concentrations across the plate, while small molecules were added in reverse concentration at the same time. B. A549 cells were infected with SARS-CoV-2. Mock “infected” cells were not infected, while DMSO treated cells with SARS-CoV-2 were highly infected. C. Heatmap of infection index of synergy screen. Infection index = Average sum intensity of dsRNA/Cell count. Red indicates higher infection index, while blue represents lower infection index. Orange, pink, and green boxes around individual data points indicate the respective representative images shown in (D-F). D., E., F. Representative images of the synergy screen for their respective monoclonal antibodies shown right above. D. All images correspond to the high monoclonal antibody (10ug/ml) and low drug concentration (1 nM) performed in this screen. E. All images correspond to the lowest efficient combination of monoclonal antibody and drug concentration. F. All images correspond to the low monoclonal antibody (0.1 ug/ml) and high drug concentration (100 nM) combination. DAPI is shown in blue, f-actin is shown in red, and dsRNA as viral infection is shown in green. Scale bar = 50µM.

To assess whether this compound could synergize with a monoclonal antibody, we tested various concentrations of Nirmatrelvir with each antibody in a dose-response curve (Fig 6A) and calculated a synergy score (ZIP synergy model [64]). As shown in Fig 6C, combination treatment between Nirmatrelvir and mAb 2355 and 2489 showed effects with increasing concentrations. Combining Nirmtrelvir with mAb 2355, the effective concentrations of each compound were reduced (Fig 6C, pink boxes) as compared to either mAb or compound alone (Fig 6C, orange and green boxes, respectively). However, none of our drug/antibody combinations reach synergy by the ZIP synergy model. In fact, the combinations showed antagonism. This could be due to the nature of their binding. Both mAbs 2355 and 2819 are Class I antibodies that target the RBD domain, while mAb 2489 is a Class III that binds to the NTD domain, as previously described (Fig 2A). Antagonism was also seen with INK128, a PI3K kinase inhibitor previously tested by our group (Sup Fig 6).

From our screen and high-dimensional analysis, we have identified many pathways that may be involved in the hijacking of host cell machinery by the virus. In particular, we observed many hits from inhibitors upstream and downstream of the PI3K pathway. These include inhibitors of EGFR, PI3K, AKT, GSK3, and mTOR. These signals are critical in cell proliferation and survival and may be critical for viral replication. Our data show that designing a high-throughput screening strategy that accounts for not only reducing a single variate (in our case, dsRNA) but also evaluating the dose-dependent effect, cell viability, and morphological response can benefit drug discovery efforts.

## DISCUSSION

Here, we developed a high-throughput and high-dimensional assay for characterizing SARS-CoV-2 infection of human lung cancer cells that captures authentic virus replication and host cell-level features. We tested our assay with monoclonal antibodies and small molecule kinase inhibitors and identified effective perturbations with minimal host cell toxicity. Our high-dimensional analytical approach allowed us to identify antibodies and kinase inhibitors missed or not reported by other groups. This work highlights a new set of compounds ready to be tested for efficacy in animal models and human clinical trials.

A challenge with the COVID-19 outbreak–and any pandemic–is the need to identify effective therapeutics rapidly. This led researchers to screen bioactive and FDA-approved compounds in simplistic assays that had limited physiological relevance. Vero E6 cell drug screens were common (19–21). In hindsight, these kidney epithelial cells derived from African green monkey were not ideal mimics of human lung physiology. Additionally, the cells fail to express the necessary proteins to ensure faithful infection and propagation by SARS-CoV-2 as a human cell line would. Additionally, SARS-CoV-2 pseudovirus systems have emerged as valuable tools for researchers studying coronavirus. These genetically modified viruses retain the key structural features of SARS-CoV-2, but lack the ability to cause a full-blown infection and cause mutations (46). By incorporating the spike protein of the original virus into a harmless viral backbone, the pseudovirus systems mimic the entry process of SARS-CoV-2 into cells. This enables scientists to study the virus’s behavior, assess the efficacy of potential treatments or vaccines, and investigate the neutralizing capacity of antibodies. The pseudovirus systems offer a safer alternative to working with the live virus and do not require BSL3 facilities, allowing for adoption among researchers worldwide. The inability of pseudovirus to replicate and spread like the actual virus limits the study of viral dynamics and replication within the host. Due to their simplified viral structure and the potential for variability between different pseudovirus constructs, findings obtained using pseudovirus systems may not fully capture the complexities of the original virus and may not accurately reflect its behavior (46–48).

Our approach using authentic SARS-CoV-2 had challenges as well. Previously, we developed a high-throughput organoid monolayer system for drug screening (49). Initially, we tested SARS-CoV-2 in our colonic organoid model as a screening platform yet observed exceedingly low infection rates. This could be due to several reasons, such as poor viral receptor expression, host cell interferon response, or structural impairments, such as well-polarized cells and mucus secretion. Using human lung cancer cells and full SARS-CoV-2 virus also resulted in low infectivity but was sufficient for our study. A caveat of the cell lines we used may be that they do not fully capture the toxicity profile of normal healthy lung epithelium. Also, single-cell image-based analysis required custom image-processing algorithms that were developed for this study, thus more time-consuming than simple cell viability or plaque assays. An additional challenge was that using authentic virus required a BSL-3 facility, which not all researchers have access to, and added additional layers to the protocol not needed in a BSL-2 facility. Our study best approximates treating COVID-19 prophylactically, which involves administering preventive measures or medications to reduce the risk of infection before exposure or early in the course of the virus. This approach can help bolster immunity and inhibit viral replication but may not model treatment after a robust infection has occurred.

Our screening identified hits from both an antibody collection and a kinase inhibitor library. The antibody hits are exciting as they could rapidly move into a clinical setting and likely have minimal toxicities. This collection of antibodies already produced two potent neutralizing antibodies that lead to Evusheld, an AstraZeneca cocktail globally distributed and accounting for thousands of people treated (50). However, as new strains of SARS-CoV-2 have evolved, Evusheld effectiveness has waned (51, 52). Our hits can be used as likely candidates for second-line antibody development. Additionally, our high-content assay could be worked into antibody discovery pipelines to rapidly identify efficacious antibodies. We observed hits from all the binding classes of antibodies. From previously defined classes, Class I are RBD-domain-targeted and had the highest hit rate in our screen. This is consistent with the RBD-domain’s functional role in ACE2 receptor binding (53, 54). Class II and V antibodies were previously shown to have binding to first SARS-

CoV-1 and SARS-CoV-2; thus, these hits might have broad efficacy against coronaviruses that infect humans with similar spike protein structures. Class III are NTD-domain-targeted and had the lowest hit rate in our screen. This is consistent with other reports (55, 56) that targeting NTD aids viral neutralization, yet the genetic variability of this domain would provide challenges. Class VI hits in our screen likely represent false negatives from the previous spike binding characterization (7). Further structural and biochemical work will need to be done to identify their precise mechanism of action. Finally, little work has been done testing antibody combinations that target the different binding classes. Could an antibody cocktail be identified that is challenging for the virus to evolve away from and lead to longer-lasting clinical efficacy?

Our kinase hits include compounds that target PI3K, EGFR, VEGFR, GSK3, mTOR and others, some of which have been identified in other studies (57–62). Our study identified GSK2636771, a PI3K inhibitor, a clinically useful therapeutic that could move swiftly into clinical trials. Hits from kinase inhibitors, in general, are significant in that 1) their well-characterized mode of action can help tease apart the host signaling pathways involved in viral entry and replication, and 2) they can serve as leads for rapid testing in human clinical trials. A caveat to our study is that it does not account for host immune response, which is a significant aspect of how patients respond to therapeutic intervention of SARS-CoV-2 infection.

Monoclonal antibodies and small molecule inhibitors are two distinct approaches for targeting SARS-CoV-2, each with their own strengths and weaknesses. mAbs can specifically bind to viral proteins, such as the spike protein of SARS-CoV-2, to neutralize the virus. They have the advantage of high specificity, potency, and the ability to directly block viral entry into host cells with minimal host toxicity. However, mAbs typically require intravenous administration, which limits their accessibility and practicality for widespread use. Additionally, the emergence of viral variants with mutations in the antibody-binding regions reduces the effectiveness of mAbs and leads to a short life as a clinical option (52, 63).

In contrast to mAbs, small molecule inhibitors are compounds that interfere with specific viral enzymes or cellular factors necessary for viral replication. They can be orally administered, allowing for easier distribution and patient compliance. Small molecule inhibitors have a broad spectrum of antiviral activity and can target conserved regions of viral proteins, potentially reducing the impact of viral variants. However, they may lack the high specificity and potency of mAbs, and the development of resistance to small molecule inhibitors can occur over time.

By combining the strengths of both approaches, there is potential for synergistic effects and improved therapeutic outcomes. Combinations of mAbs and small molecule inhibitors can provide a dual mechanism of action, targeting different stages of the viral life cycle. In our study, this combination approach did not show statistically significant synergy. In fact, we observed antagonism. This could be due to the distinct, likely independent MOA between the drugs and antibodies. This is potentially important in a clinical setting where a patient may, in fact, be receiving PI3K inhibitors for cancer and may respond poorly to COVID-neutralizing antibody therapy. Future work testing mAbs combinations that target multiple regions of the spike protein or small molecules in synergistic pathways could potentially enhance antiviral efficacy, mitigate the emergence of drug resistance, and offer broader coverage against viral variants. Understanding the complementary strengths and weaknesses of monoclonal antibodies and small molecule inhibitors is crucial in developing effective treatment strategies against SARS-CoV-2. Continued research and clinical trials are needed to optimize and evaluate the potential of these therapeutic modalities in combating COVID.

## METHODS

### CELL CULTURE

Calu-3 (ATCC, HTB-55) and Caco-2 (ATCC, HTB-37) cells were maintained at 37°C in 5% CO_2_ in DMEM with high glucose and 1x GlutaMAX (Invitrogen 35050-061), containing 10% heat inactivated fetal bovine serum (FBS), and100 U/mL of penicillin-streptomycin. Cells were seeded at 5,000 cells per well in 384 Screenstar microplates (Greiner #781866) and cultured overnight to allow for adherence. The next day cells were treated with either kinase inhibitors or antibodies before infection for 48 hours with SARS-CoV-2 virus. After 48 hours of infection, plates were submerged in 4% paraformaldehyde/sucrose for 30 mins and stained for immunofluorescence.

### IMMUNOFLUORESCENCE

Cells were permeabilized in 0.2% triton-x in PBS for 10 minutes. The plates were then blocked with 5% BSA in PBS for 1 hour at room temperature before staining. Cells were stained with primary antibody for dsRNA (SCIONS 10010500) at 1:1000 in BSA/PBS overnight at 4C in the dark. Cells were washed thrice in 0.1% Tween-20/PBS (PBST) for 15 minutes and incubated in secondary 546 Alexa Fluor anti-mouse 1:1000 (Fisher A11003) and phalloidin 647 (Biotium 00041) 1:80 for 2 hours in the dark at room temperature in BSA/PBS. Plates were washed thrice with PBST and then incubated with 1x DAPI solution in PBS for 30 minutes at room temperature in the dark. Plates were then washed and stored in PBS. Plates were imaged on a Nikon Eclipse Ti2 automated microscopy system on 20x objective. Intensity of dsRNA was measured using NIKON NIS Elements General Analysis software for the Kinase inhibitor assay Neutralizing antibody assay and cell segmentation was performed using Cell Profiler for the cross analysis and UMAP production. Graphs were produced in Jupyter Notebook using R software.

### VIRAL INFECTION

USA-WA1/2020 isolate of SARS-CoV-2 was obtained from WRCEVA and a stock was generated by infecting Vero CCL81 cells for 48h. Lysate was collected and pelleted to remove cell debris. Genome sequencing of the virus by Nanopore confirmed the genome sequence to be identical to the GenBank WA1/2020 sequence (MN985325.1), with no mutations in the spike furin cleavage site. Virus was added 1:1 to plates and allowed to incubate at 37°C for 48 hours. Plates were submerged in 4% paraformaldehyde/sucrose solution for 30 minutes to fix cells and decontaminate the plates for removal from BSL-3 facilities. All work with infectious SARS-CoV-2 was performed in Institutional Biosafety Committee-approved BSL3 using appropriate positive pressure air respirators and protective equipment.

#### Viral assay For A549 ace2/TMPRSS2 cells

SARS-Cov-2 variants Omicron B.1.1.529 and Delta B. 1.617.2 SARS-COV-2 were obtained from BEI. A plaque assay using VeroE6 cells was done to generate viral titers. A MOI of .01 was used for WA1 with the viral titer measured at 9e6 pfu/mL. A MOI of .03 was used for Omicron BA.1 with a viral titer of 2.4e6 pfu/mL. a MOI of .01 was used for delta with a viral titer of 4e6 pfu/mL. Cells were infected and left to incubate for 48 hrs at 37°C/5% CO2 for 48 hours. Note: the calculation is the same for any of the MOIs that were used.

After 48 hrs media was removed and 4% paraformaldehyde phosphate buffer solution (Fujifilm Wako Pure chemical Corporation) (PFA) was used to fix cells. PFA was added for 30 mins per BSL3 decontamination protocol. PFA was removed and cells were washed using PBS and then a 100uL of new PBS was added to each well and sent for imaging. Plates were sealed using parafilm.

### NEUTRALIZING ANTIBODY ASSAY

Cells were seeded at 5,000 cells per well in 384-well microplates (Greiner 789836) and allowed to adhere overnight. A panel of ∼500 neutralizing antibodies (7) were added in four ten-fold doses starting at 1:100 on stock antibodies. The cells were then treated with SARS-CoV-2 virus for 48 hours before fixing in 4% paraformaldehyde/sucrose solution for 30 minutes and stained in the methods described. Plates were imaged on a Nikon Eclipse Ti2 automated microscope at 20x in the same manner as described above. Six frames were captured and analyzed for dsRNA intensity on the whole field and DAPI was used to measure cell count. Normalization was performed the same as the kinase inhibitor assay. Raw data was extracted from NIKON NIS Elements software and plotted in R.

### KINASE INHIBITOR ASSAY

Cells were seeded at 5,000 cells per well in 384-well microplates (Greiner 789836) and allowed to adhere overnight. A panel of ∼800 kinase inhibitors (ApeXBio L1024) were added in two doses, 10µM and 100nM, on the cells. The cells were then treated with SARS-CoV-2 virus for 48 hours before fixing in 4% paraformaldehyde/sucrose solution for 30 minutes and stained in the methods described. Plates were imaged on a Nikon Eclipse Ti2 automated microscope at 20x. Six frames were captured and analyzed for dsRNA and DAPI was used to measure cell count. Overall intensity for each inhibitor was measured by segmenting images for dsRNA threshold and measuring the sum intensity of 546 and dividing by cell count to normalize by well-to-well variation. Wells were then normalized to control wells with mock infection and treatment with DMSO to account for plate-to-plate variability. Raw data was extracted from NIKON NIS Elements software and plotted in R. Follow-up confirmation tests were performed with 34 top kinase inhibitors that showed increased efficiency with higher dose and were drugs in clinical trials. Those kinase inhibitors were dosed in a 14-dose response curve. Cells were plated the same as described above and drugs diluted in DMSO were delivered in a dose curve using a Tecan d300e Digital Dispenser. Dose-response was plotted in GraphPad Prism as the percent of sum dsRNA to control. Cell count was the average of all wells and counting nuclei by DAPI using NIKON NIS Elements software and plotted in prism. EC_50_s were calculated in GraphPad. Error bars represent the standard error of the mean of all wells and duplicates.

### COMBINATION OF NEUTRALIZING ANTIBODY AND KINASE INHIBITOR ASSAY

Cells were seeded at 8,000 cells per well in a 96-well plate (Greiner 655090) and allowed to adhere overnight. One antibody was added per plate in a 8-dose response curve from 0.01 ug/ml to 10 ug/ml from left to right. At the same time, either compound (INK 128 or Nirmatrelvir) was added in a 7-dose response curve from 1 nM to 100 nM. The cells were then treated with SARS-COV-2 virus for 48 hours before fixing in 4% paraformaldehyde/sucrose solution for 30 minutes and stained in the methods described above. Plates were imaged on a Nikon Eclipse Ti2 automated microscope at 20x in the same manner as described above. Six frames were captured and analyzed for dsRNA intensity on the whole field and DAPI was used to measure cell count. Normalization was performed the same as the kinase inhibitor assay. Raw data was extracted from NIKON NIS Elements software and plotted in R. Cells were plated the same as described above and drugs diluted in DMSO were delivered in a dose curve using a Tecan d300e Digital Dispenser. Heat maps were generated in R.

### SEGMENTATION AND FEATURE EXTRACTION

We used CellProfiler (CP) software to segment images into individual cells. DAPI was used to segment individual nuclei into primary objects. Phalloidin was used as secondary objects to define individual cell borders and to be used as cell masks. Features were then extracted for individual cells using CP. To calculate the dsRNA features, we used the dsRNA channel and cell mask to quantify different measures based on intensity of dsRNA channel within the cell region.

### DATA PREPROCESSING

#### Finding infection rate of the experiment

To find the rate of infection we plotted cellular area vs mean intensity of dsRNA in both mock and infected cells treated with DMSO. The most infected cells make up 1% of cell population and were used to evaluate response of both monoclonal antibodies and kinase inhibitors in this study.

Moving forward, the top 1% of cells in all conditions were evaluated for response.

#### Normalizing the data

Normalization factor is calculated for each screening set (kinase inhibitor or neutralizing antibody) separately. This factor is the mean of mock DMSO cells in all the plates in each experiment for each feature. The cells are then standardized to make sure all the features are treated the same in the data analysis.

#### Calculating UMAP

After standardizing the data, we project it into UMAP space (arXiv:1802.03426) using the UMAP package in R. Individual points represent the average of a single treatment condition at a single dose. Plots were colored according to increasing dsRNA intensity.

### CLUSTERING ANALYSIS

Mock DMSO wells in each plate were used as normalization factors. All measures were averaged, and single cell features were normalized to the mock DMSO condition. We then calculated the mean per condition and projected the data into UMAP space. Using hierarchical clustering, we clustered the data points to find out how the data is scattered in UMAP space and which conditions end up next to our controls.

## DATA AVAILABILITY

Data is available upon request; contact. Image files and Cell Profiler pipelines are housed on Thorne Lab servers at the University of Arizona and available for download upon request.

## ACKNOWLEDGEMENTS

We thank the Crowe, Campos, and Thorne labs for their support and advice. We also thank C. Bradshaw, J. Urlaub, and J. Nikolich-Žugich for use of the BSL-3 facility, propagating SARS-CoV-2 virus, and providing insights and support.

## FUNDING AND ADDITIONAL INFORMATION

This work was supported by the National Institutes of Health (T32CA009213 to C.R.C., DK103126, GM147128 to C.A.T, GM136853 to S.K.C., GM130864 to A.L.P., AI157155 to J.E.C.), The ARCS Foundation Phoenix (Theresa F Jennings Award to C.R.C.), Wellcome Trust, UK (WT223952/Z/21/Z to C.A.T.), the State of Arizona (TRIF to C.A.T., K.V.D.), (TRIF, UA COVID-19 Seed Grant #002196 to S.K.C.), and the Defense Advanced Research Projects Agency (HR0011-18-3-0001 to J.E.C.).

