## Supplemental Info for "Cell-based high-content approach for SARS-CoV-2 neutralization identifies unique monoclonal antibodies and PI3K pathway inhibitors"

USA.

^2^Cancer Biology Graduate Interdisciplinary Program, The University of Arizona, Tucson, Arizona, 85724, USA.

^3^The Jackson Laboratory, Bar Harbor, ME, 04609, USA

^4^Asthma and Airway Disease Research Center, The University of Arizona, Tucson, 85724, USA.

^5^Department of Immunobiology, BIO5 Institute, University of Arizona, Tucson, AZ, 85724, USA.

^6^ Department of Pharmacology & Toxicology, R. Ken Coit College of Pharmacy, University of Arizona, Tucson 85721, Arizona, United States

^7^Department of Molecular and Cellular Biology, The University of Arizona, Tucson, 85724, USA.

^8^Vanderbilt Vaccine Center, Vanderbilt University Medical Center, Nashville, TN, 85724 USA.

^9^Department of Pathology, Microbiology, and Immunology, Vanderbilt University Medical Center, Nashville, TN, 37232, USA

^10^Department of Pediatrics, Vanderbilt University Medical Center, Nashville, TN, 37232, USA.

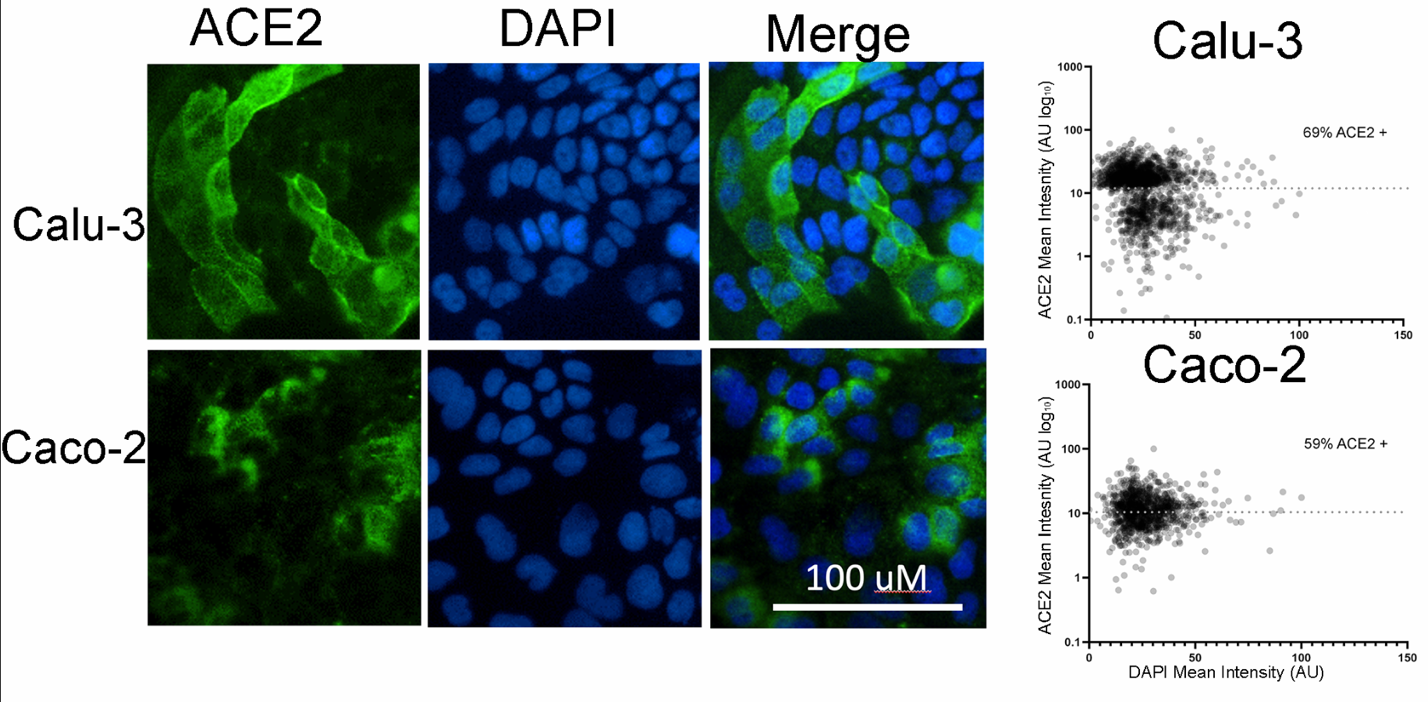

**Fig S1: Calu-3 Cells more Permissible to Infection than Caco-2 Cells**

Immunofluorescent staining for ACE2 (green), DNA (blue) shows ACE2 receptor expression in Calu-3 and Caco-2 cells. Expression of ACE2 receptor was normalized by ACE2 mean intensity and plotted for Calu-3 cells and Caco-2 cells shown in scatter plot. Percent ACE2 positive cells for each group were calculated based on amount of ACE2 positive cells among all cells in dataset.

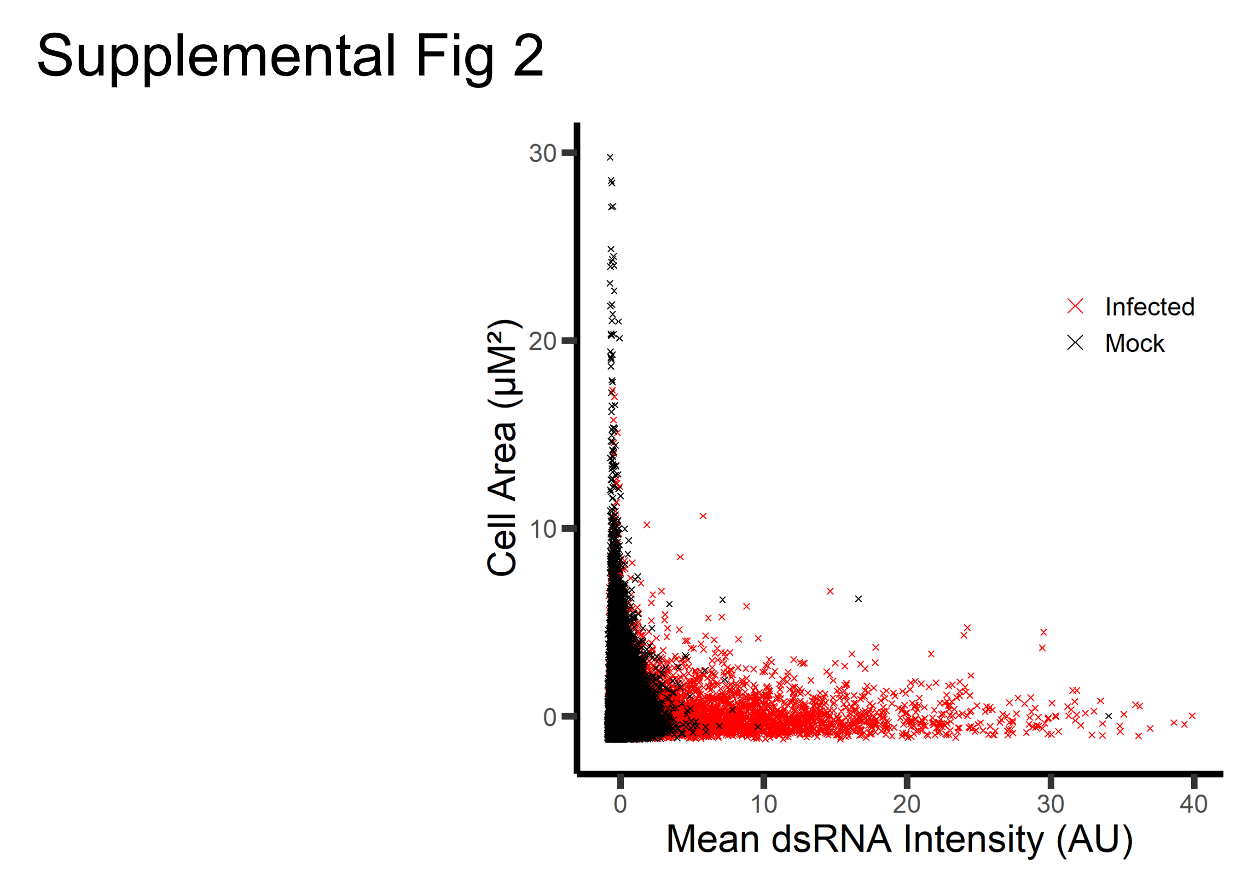

**Fig S2: Infection Rate of Calu-3 Cells with SARS-CoV-2**

Scatter plot of Calu-3 cells mock or SARS-CoV-2 infected. Cells were stained for dsRNA, imaged, segmented in single-cell objects, and dsRNA quantified.

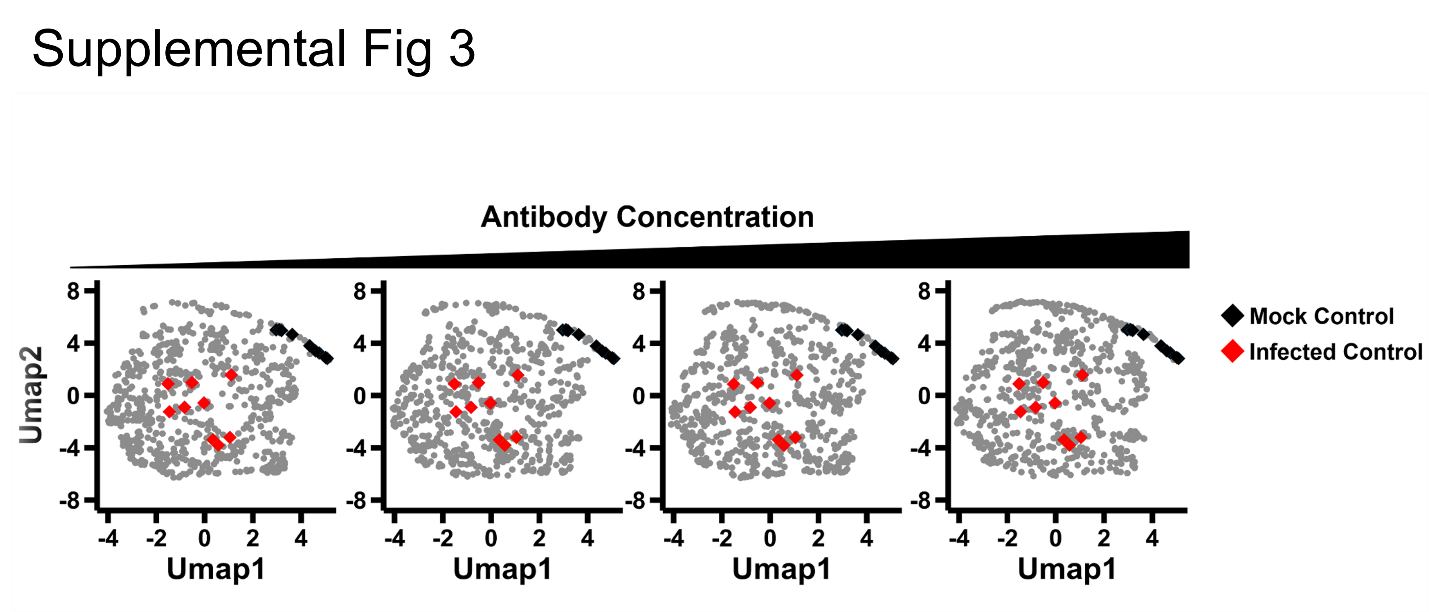

**Fig S3: Antibody Response by Concentration**

UMAP of antibodies separated by concentration. Right to left show serial dilutions. The shift of the density of the points indicates that with an increasing concentration of antibodies, the cell phenotype subtly moves closer to the mock control (black diamonds) and away from the infected control (red diamonds).

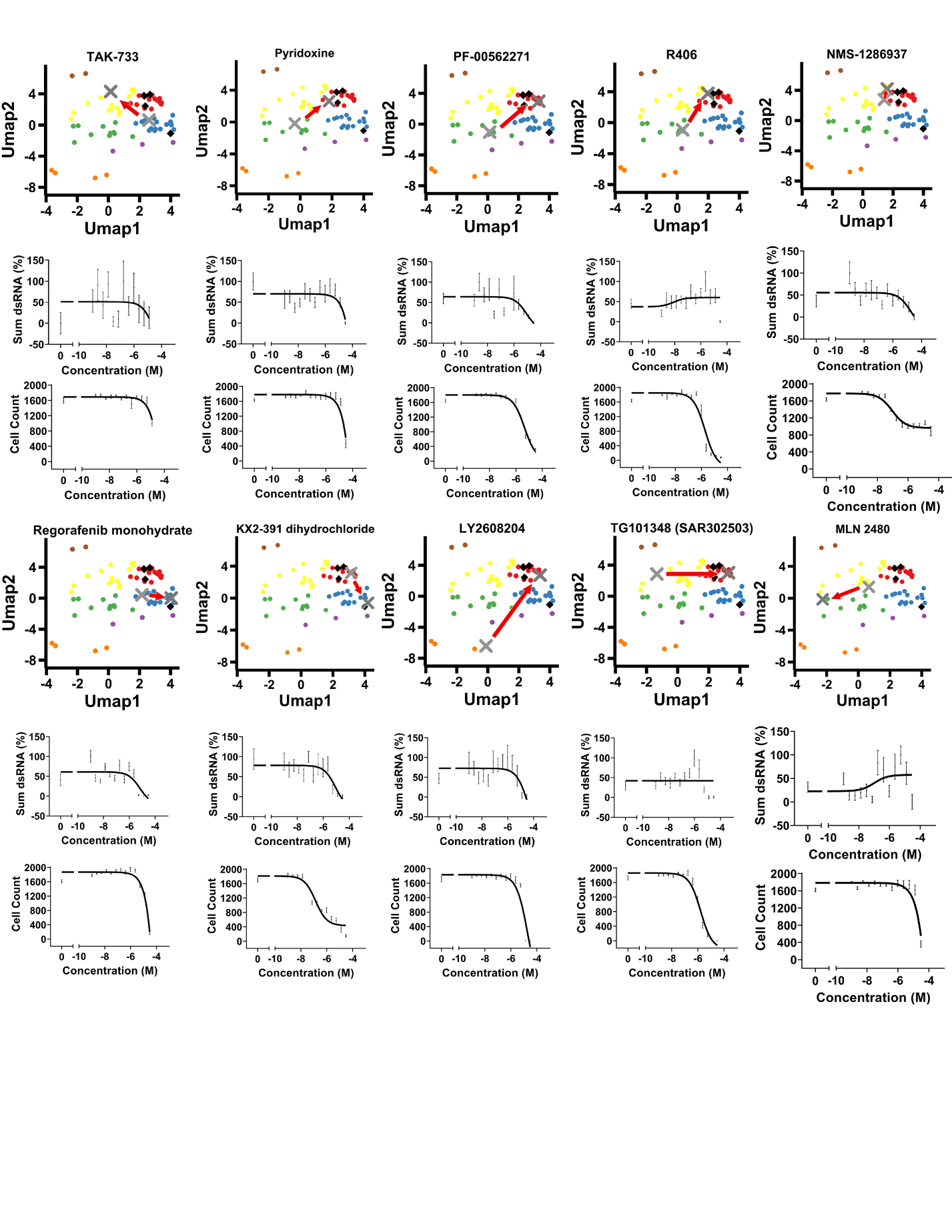

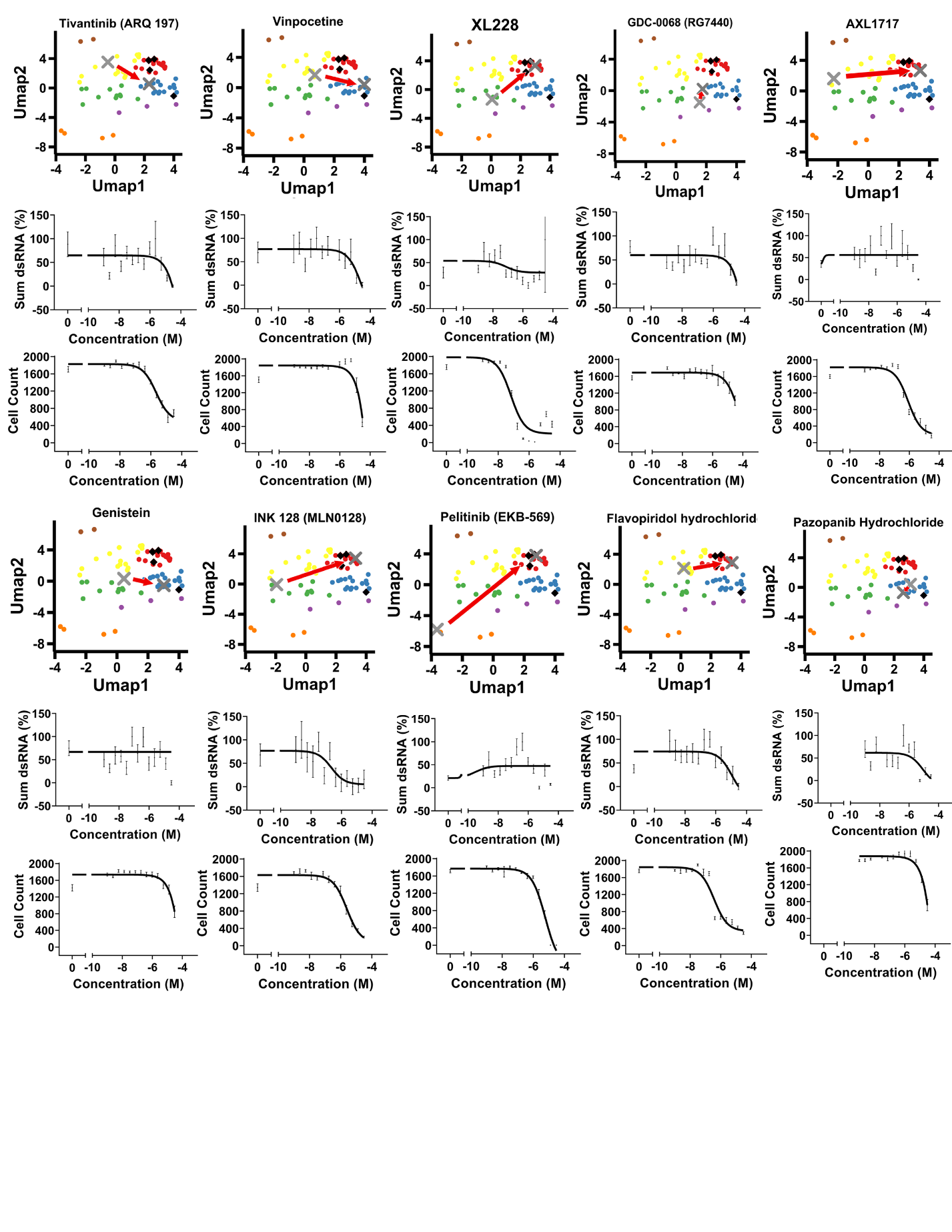

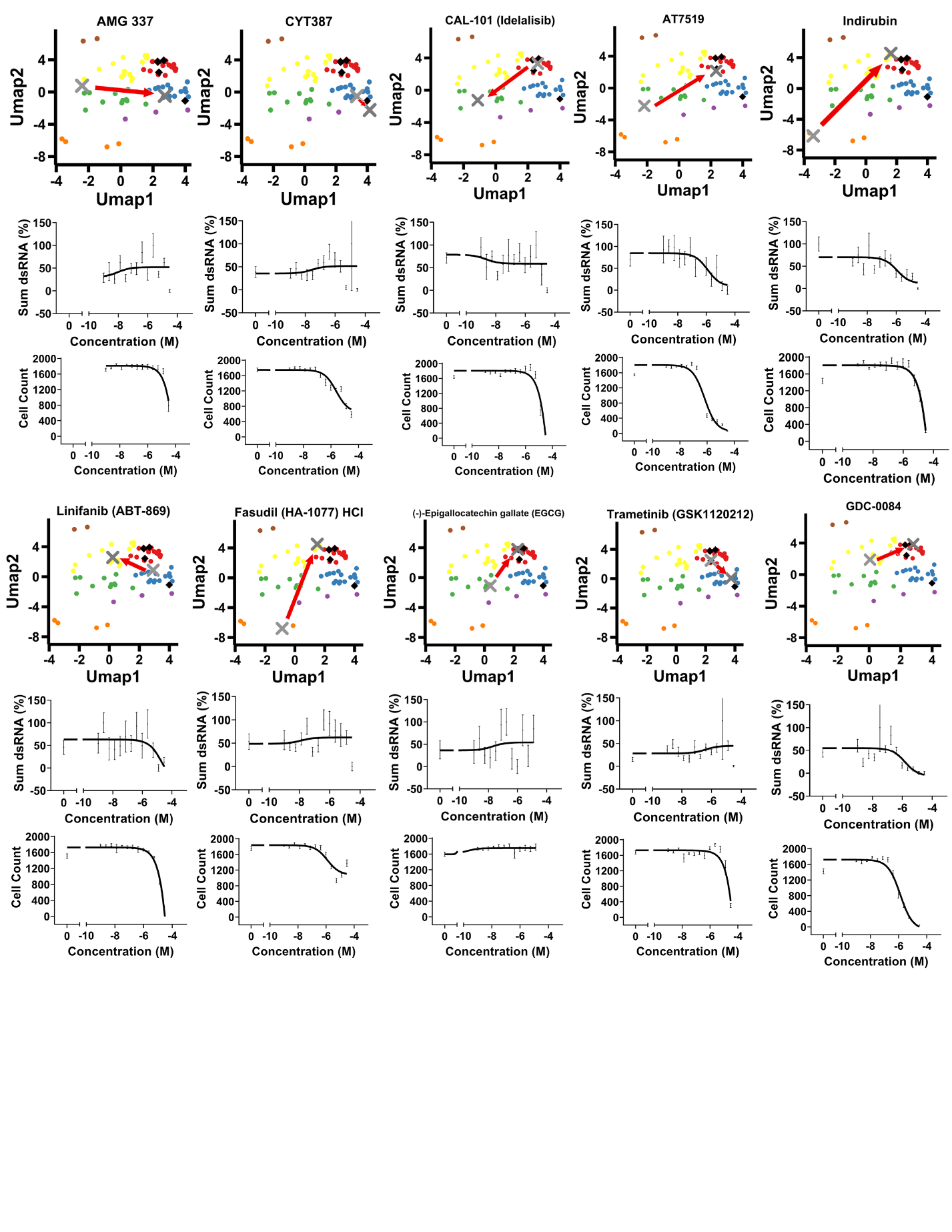

**Fig S4: Kinase Inhibitor Screen Results**

As in main Figure 5 C-E, a continuation of plots representing the retesting of strong hits from the initial kinase inhibitor screen. The gray X’s indicate the low and high doses from the initial screen. The red arrow points from the low dose to the high dose to emphasize the shift in location UMAP space. Sum dsRNA was normalized to percent of response to control. Cell count from dose-response curve was also measured by counting DAPI. Error bars represent SEM.

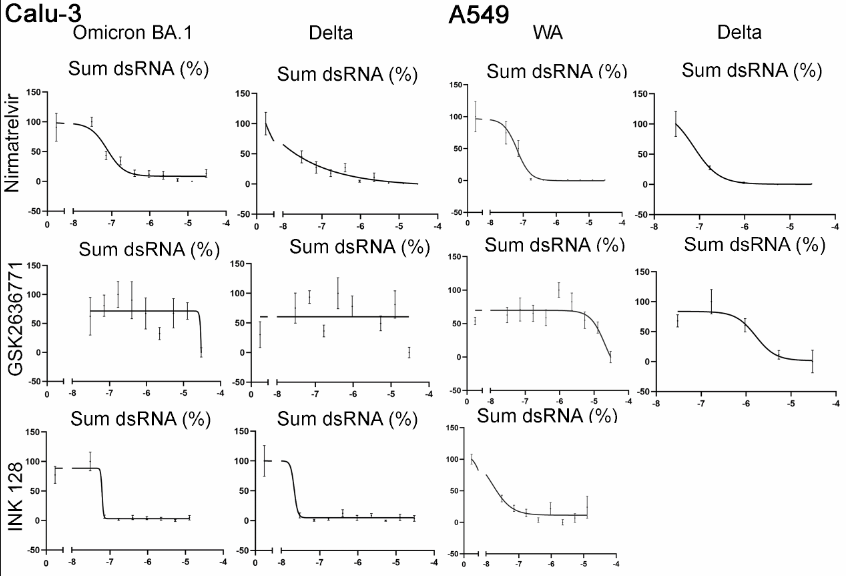

**Fig S5: Kinase Inhibitor Screen Results with Omicron Ba.1 and Delta variant in Calu3 and A549 cells**

To enhance the robustness and applicability of our findings, we validated the efficacy of our positive control and two compounds from our kinase inhibitor screen across an additional cell line and against two more virus variants, Omicron BA.1 and Delta. The three compounds were tested in an 8-concentration dose response. Sum dsRNA was normalized to percent of response to control. Error bars represent SEM.

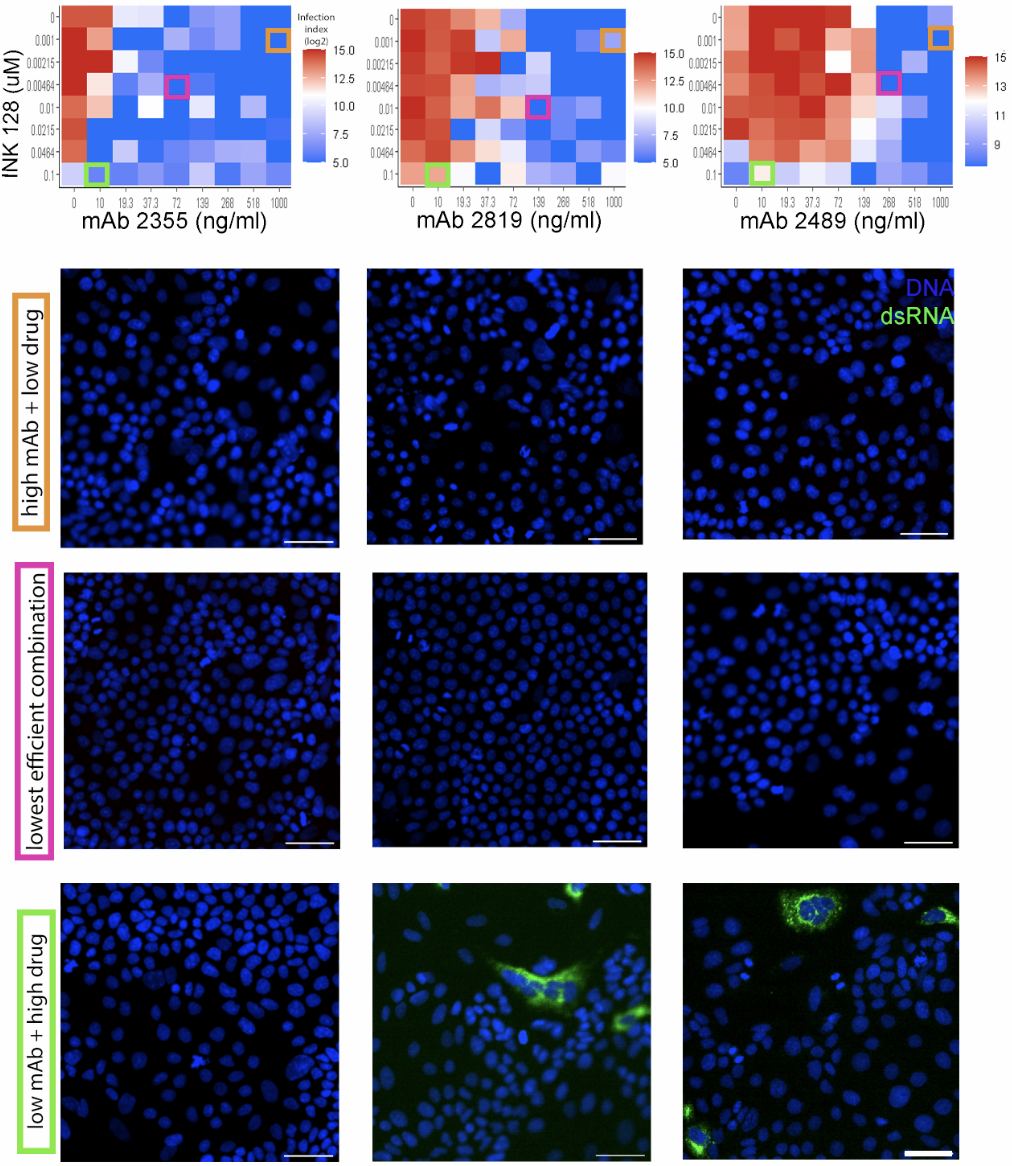

**Fig S6: Combination of Neutralizing Antibody and Kinase Inhibitor Results.** As in main Figure 6 C-F, a continuation of heatmaps and respective representative images are shown for the combination treatment of monoclonal antibodies 2355, 2819, and 2489, with mTOR inhibitor, INK 128. Representative images of a synergy experiment. The infection index was calculated by measuring the average sum intensity of dsRNA per cell count. DAPI is shown in blue, and dsRNA as viral infection is shown in green. Scale bar = 50µM.

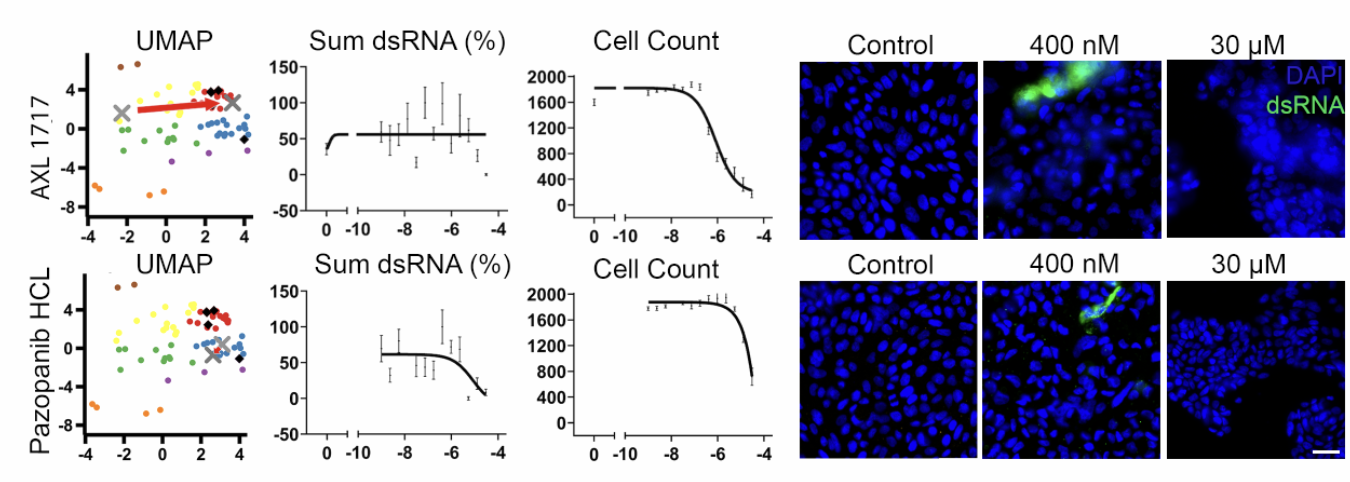

**Fig S7: Candidates among our Top Kinase Inhibitor**

As in main Figure 5 C-F, a continuation of our candidates from the kinase inhibitor screen are shown. AXL 1717 and Pazopanib HCL were retested in a 12-concentration dose-response curve. The gray X indicates the low and high doses from the initial screen. The red arrow points from the low dose to the high dose to emphasize the shift in location UMAP space. Sum dsRNA was normalized to the percent of response compared to control. Cell count from dose response curve was measured by counting DAPI-stained cells. Right side, representative images of the specified kinase inhibitor with low and high concentrations. Nuclei are stained with DAPI in blue and dsRNA is stained in green. Scale bar = 50µM. Error bars represent SEM.

**Table S1: Antibody dsRNA Area Under the Curve (AUC)**

|  | **Decreasing Dosage 1:20** | | | |  |
| --- | --- | --- | --- | --- | --- |
| **Antibody ID** | **Dose 1** | **Dose 2** | **Dose 3** | **Dose 4** | **Sum** |
| **2355** | **0** | **0** | **0** | **0** | **0** |
| **2381** | **0** | **0** | **0** | **0** | **0** |
| **2504** | **0** | **0** | **0** | **0** | **0** |
| **2691** | **0** | **0** | **0** | **0** | **0** |
| **2819** | **0** | **0** | **0** | **0** | **0** |
| **2832** | **0** | **0** | **0** | **0** | **0** |
| **2498** | **0** | **0** | **0** | **0.034282** | **0.034282** |
| **2391** | **0** | **0** | **0** | **0.042931** | **0.042931** |
| **2499** | **0.004381** | **0** | **0** | **0.086103** | **0.090484** |
| **2941** | **0.001553** | **0.001655** | **0.001602** | **0.105511** | **0.110322** |
| **2955** | **0** | **0** | **0.092276** | **0.02758** | **0.119856** |
| **2308** | **0** | **0.006481** | **0.003528** | **0.111864** | **0.121874** |
| **2539** | **0.009391** | **0** | **0.032509** | **0.091817** | **0.133716** |
| **2514** | **0** | **0.001193** | **0.001901** | **0.146693** | **0.149787** |
| **2531** | **0.002623** | **0.002023** | **0.036593** | **0.13553** | **0.176769** |
| **2353** | **0** | **0** | **0.094756** | **0.104947** | **0.199703** |
| **2733** | **0** | **0** | **0** | **0.25545** | **0.25545** |
| **2290** | **0** | **0** | **0.076816** | **0.219281** | **0.296097** |
| **2838** | **0.01561** | **0.125523** | **0.093655** | **0.067743** | **0.30253** |
| **2413** | **0** | **0.003581** | **0.176016** | **0.126839** | **0.306435** |
| **2859** | **0.064683** | **0.080574** | **0.074623** | **0.119486** | **0.339366** |
| **2485** | **0** | **0.024703** | **0.011633** | **0.312655** | **0.34899** |
| **2389** | **0.028526** | **0.03034** | **0.249269** | **0.141058** | **0.449193** |
| **2813** | **0.006993** | **0.004478** | **0.149629** | **0.301735** | **0.462835** |
| **2841** | **0** | **0** | **0.122719** | **0.345178** | **0.467897** |
| **2807** | **0.006037** | **0.306831** | **0** | **0.175823** | **0.488692** |
| **2341** | **0.233236** | **0.086005** | **0.124472** | **0.117795** | **0.561509** |
| **2919** | **0** | **0.100397** | **0.250101** | **0.228387** | **0.578884** |
| **2415** | **0.055398** | **0.176151** | **0.344762** | **0.010173** | **0.586484** |
| **2952** | **0** | **0.049558** | **0.002553** | **0.582294** | **0.634405** |
| **2489** | **0.001251** | **0** | **0.042514** | **0.634815** | **0.67858** |
| **2448** | **0.121064** | **0.131853** | **0.231625** | **0.202726** | **0.687268** |
| **2417** | **0.017218** | **0.218993** | **0.361723** | **0.096315** | **0.694249** |
| **2601** | **0.145502** | **0.174933** | **0.129349** | **0.261008** | **0.710794** |
| **2393** | **0.171955** | **0.151889** | **0.285624** | **0.1041** | **0.713567** |
| **2406** | **0.404072** | **0.098853** | **0.212046** | **0** | **0.71497** |
| **2835** | **0** | **0.01373** | **0.05808** | **0.70013** | **0.77194** |
| **2822** | **0.418079** | **0** | **0.010095** | **0.371416** | **0.79959** |
| **2589** | **0.007277** | **0.060538** | **0.451686** | **0.307908** | **0.827408** |
| **2717** | **0** | **0.087156** | **0.502349** | **0.261643** | **0.851147** |
| **2452** | **0.194675** | **0.292466** | **0.047501** | **0.323623** | **0.858266** |
| **2625** | **0.530779** | **0.109737** | **0.118047** | **0.107608** | **0.866171** |
| **2618** | **0.014705** | **0.153277** | **0.635626** | **0.07575** | **0.879358** |
| **2861** | **0.446715** | **0.21932** | **0.239473** | **0** | **0.905509** |
| **2399** | **0.03733** | **0.543622** | **0.282536** | **0.054105** | **0.917593** |
| **2358** | **0.097848** | **0.356754** | **0.455184** | **0.017341** | **0.927128** |
| **2364** | **0.002038** | **0.336557** | **0.355746** | **0.264111** | **0.958451** |
| **2398** | **0** | **0.072788** | **0.430367** | **0.45979** | **0.962946** |
| **2791** | **0.114229** | **0.00392** | **0.453327** | **0.396017** | **0.967492** |
| **2557** | **0.082252** | **0.358082** | **0.297132** | **0.232357** | **0.969823** |
| **2344** | **0.295984** | **0.404711** | **0.157044** | **0.112904** | **0.970643** |
| **2678** | **0** | **0** | **0.990999** | **0** | **0.990999** |
| **2394** | **0.212002** | **0.329673** | **0.051135** | **0.460715** | **1.053525** |
| **2369** | **0.087283** | **0.374468** | **0.46364** | **0.138245** | **1.063636** |
| **2354** | **0** | **0** | **0.807096** | **0.25728** | **1.064376** |
| **2894** | **0.007649** | **0.238954** | **0.516522** | **0.307708** | **1.070834** |
| **2295** | **0.492476** | **0.213316** | **0.257153** | **0.142491** | **1.105436** |
| **2526** | **0** | **0.926408** | **0.107009** | **0.074027** | **1.107444** |
| **2352** | **0.029007** | **0.367883** | **0.394524** | **0.323376** | **1.114791** |
| **2946** | **0.237466** | **0.154596** | **0.462878** | **0.264033** | **1.118973** |
| **2343** | **0.287072** | **0.386099** | **0.288903** | **0.173125** | **1.1352** |
| **2338** | **0** | **0.548486** | **0.149338** | **0.449417** | **1.14724** |
| **2572** | **0.101825** | **0.082369** | **0.271511** | **0.701709** | **1.157415** |
| **2837** | **0.247833** | **0.167945** | **0.365666** | **0.378159** | **1.159603** |
| **2711** | **0.958049** | **0.017952** | **0** | **0.204777** | **1.180777** |
| **2374** | **0.899838** | **0.01141** | **0.027563** | **0.246275** | **1.185086** |
| **2403** | **0.119641** | **0.425173** | **0.460479** | **0.184419** | **1.189713** |
| **2382** | **0.018432** | **0.111559** | **0.660831** | **0.415412** | **1.206234** |
| **2573** | **0.581382** | **0.055242** | **0.249787** | **0.320385** | **1.206796** |
| **2357** | **0** | **0.380225** | **0.50404** | **0.332963** | **1.217228** |
| **2412** | **0.146403** | **0.299915** | **0.714624** | **0.058196** | **1.219138** |
| **2386** | **0.329172** | **0.299022** | **0.175305** | **0.41617** | **1.219669** |
| **2694** | **0.026897** | **0.001634** | **0.360749** | **0.860283** | **1.249565** |
| **2337** | **0.251625** | **0.380675** | **0.076447** | **0.559461** | **1.268209** |
| **2350** | **0.068943** | **0.922148** | **0.281516** | **0** | **1.272607** |
| **2385** | **0.261145** | **0.373418** | **0.525057** | **0.113881** | **1.273501** |
| **2562** | **0** | **0** | **0.672187** | **0.604805** | **1.276992** |
| **2397** | **0.240649** | **0.147788** | **0.758177** | **0.132513** | **1.279128** |
| **2848** | **0.124972** | **0.110146** | **0.371466** | **0.702326** | **1.308911** |
| **2376** | **0.234343** | **0.221284** | **0.346777** | **0.511051** | **1.313454** |
| **2414** | **0.672089** | **0.430733** | **0.199986** | **0.01592** | **1.318727** |
| **2410** | **0.37644** | **0.379801** | **0.141918** | **0.426466** | **1.324625** |
| **2554** | **0** | **0.51631** | **0.61478** | **0.22351** | **1.3546** |
| **2447** | **0.450343** | **0.229324** | **0.275503** | **0.405192** | **1.360362** |
| **2395** | **0.150956** | **0.145448** | **0.422535** | **0.651117** | **1.370056** |
| **2342** | **0.009955** | **0.332174** | **0.343221** | **0.68721** | **1.372559** |
| **2379** | **0.091224** | **0.39386** | **0.219579** | **0.669286** | **1.373949** |
| **2834** | **0.052781** | **0.142245** | **1.158619** | **0.037858** | **1.391502** |
| **2468** | **0.31914** | **0.103657** | **0.324739** | **0.648859** | **1.396395** |
| **2535** | **0.356388** | **0.345201** | **0.158675** | **0.541531** | **1.401795** |
| **2416** | **0.218385** | **0.421309** | **0.329555** | **0.446087** | **1.415336** |
| **2882** | **0.017052** | **0.33713** | **0.421557** | **0.647834** | **1.423573** |
| **2846** | **0.51768** | **0.277875** | **0.272542** | **0.358884** | **1.42698** |
| **2828** | **0** | **0.026425** | **0.193597** | **1.22975** | **1.449773** |
| **2563** | **0.01307** | **0.813588** | **0.352049** | **0.272319** | **1.451026** |
| **2543** | **0.125623** | **0.570078** | **0.305744** | **0.480567** | **1.482012** |
| **2684** | **0** | **0.027748** | **0.160733** | **1.294534** | **1.483015** |
| **2335** | **0.58079** | **0.814525** | **0.029903** | **0.062586** | **1.487804** |
| **2937** | **0.536761** | **0.555386** | **0.286201** | **0.10965** | **1.487997** |
| **2346** | **0.669053** | **0.312915** | **0.48033** | **0.036328** | **1.498626** |
| **2862** | **0.006401** | **0.629049** | **0.507815** | **0.355599** | **1.498865** |
| **2375** | **0.186451** | **0.074613** | **0.684057** | **0.558039** | **1.50316** |
| **2704** | **0** | **1.272649** | **0** | **0.244495** | **1.517144** |
| **2493** | **0.530514** | **0** | **0.335125** | **0.653378** | **1.519016** |
| **2392** | **0.276619** | **0.21324** | **0.217442** | **0.819458** | **1.526758** |
| **2256** | **0** | **0.058808** | **1.34572** | **0.139548** | **1.544076** |
| **2863** | **0** | **0.460135** | **0.219564** | **0.86586** | **1.54556** |
| **2367** | **0.04517** | **0.438841** | **0.372166** | **0.699181** | **1.555358** |
| **2878** | **0** | **0.099192** | **0.419552** | **1.043154** | **1.561897** |
| **2506** | **0.166971** | **0.618363** | **0.19734** | **0.589126** | **1.5718** |
| **2935** | **0.214115** | **0.805124** | **0.340223** | **0.214049** | **1.57351** |
| **2360** | **0.052582** | **0.113324** | **0.850829** | **0.560743** | **1.577478** |
| **2926** | **0.252811** | **0.468572** | **0.401464** | **0.45535** | **1.578197** |
| **2461** | **0** | **0.635785** | **0.317831** | **0.652133** | **1.605749** |
| **2511** | **0.516783** | **0.413334** | **0.582638** | **0.093859** | **1.606615** |
| **2939** | **0.114109** | **0.000752** | **0.762617** | **0.733193** | **1.61067** |
| **2564** | **0.101904** | **0.626477** | **0.817961** | **0.065657** | **1.611998** |
| **2815** | **0.116341** | **0.51552** | **0.860618** | **0.127089** | **1.619568** |
| **2361** | **0.184217** | **0.964469** | **0.027533** | **0.448931** | **1.625149** |
| **2333** | **0.35487** | **0.060482** | **0.777561** | **0.43648** | **1.629393** |
| **2459** | **0.154364** | **0.844207** | **0.146114** | **0.491426** | **1.636112** |
| **2318** | **0** | **0.052203** | **0.724615** | **0.879586** | **1.656403** |
| **2460** | **0.70769** | **0.131317** | **0.156018** | **0.661543** | **1.656567** |
| **2821** | **0.289918** | **0.024661** | **0.010205** | **1.33734** | **1.662123** |
| **2592** | **0.465699** | **0.223065** | **0.4457** | **0.527997** | **1.662461** |
| **2432** | **0.109403** | **0.523232** | **0.522051** | **0.527254** | **1.681939** |
| **2574** | **0.799457** | **0.218088** | **0.49177** | **0.176032** | **1.685348** |
| **2916** | **0.907832** | **0.141205** | **0.072431** | **0.576383** | **1.69785** |
| **2703** | **0.24832** | **0.706552** | **0.002597** | **0.756944** | **1.714412** |
| **2540** | **0.814356** | **0.554917** | **0.101487** | **0.243749** | **1.714509** |
| **2693** | **0** | **0** | **0.327079** | **1.388966** | **1.716045** |
| **2400** | **0.376445** | **0.332316** | **0.873278** | **0.13673** | **1.718769** |
| **2345** | **0.196692** | **0.430931** | **0.080495** | **1.015436** | **1.723554** |
| **2430** | **0.305132** | **0.886057** | **0.307998** | **0.230029** | **1.729216** |
| **2561** | **0.855037** | **0.191069** | **0.030162** | **0.654911** | **1.731179** |
| **2252** | **0.615725** | **0.452421** | **0.0942** | **0.577859** | **1.740206** |
| **2559** | **0.709994** | **0.22225** | **0.563307** | **0.249944** | **1.745496** |
| **2409** | **0.204743** | **0.140997** | **1.07789** | **0.33007** | **1.753699** |
| **2953** | **0.015388** | **0.682215** | **0.468093** | **0.588481** | **1.754177** |
| **2753** | **0** | **0.239413** | **0.957162** | **0.559319** | **1.755894** |
| **2437** | **0** | **1.27041** | **0.08044** | **0.422877** | **1.773727** |
| **2829** | **0.480649** | **0.351415** | **0.179437** | **0.762956** | **1.774458** |
| **2443** | **0.30934** | **0.778197** | **0.550969** | **0.137304** | **1.77581** |
| **2914** | **0.43818** | **0.19857** | **0.8892** | **0.251355** | **1.777305** |
| **2843** | **0.655609** | **0.273933** | **0.23407** | **0.622801** | **1.786412** |
| **2396** | **0.857661** | **0.245742** | **0.171685** | **0.511681** | **1.786769** |
| **2585** | **0.713776** | **0.511482** | **0.154945** | **0.443632** | **1.823835** |
| **2790** | **0** | **1.033334** | **0.498794** | **0.296622** | **1.82875** |
| **2555** | **0.341677** | **0.039223** | **1.103932** | **0.347288** | **1.83212** |
| **2875** | **0.583443** | **0.522881** | **0.202138** | **0.529446** | **1.837908** |
| **2510** | **0.007015** | **0.13658** | **0.552229** | **1.147378** | **1.843202** |
| **2934** | **0.163693** | **0.501926** | **0.909343** | **0.268961** | **1.843923** |
| **2626** | **0.72358** | **0.150528** | **0.297478** | **0.679138** | **1.850724** |
| **2545** | **0.179466** | **0.500915** | **0.406062** | **0.767213** | **1.853657** |
| **2594** | **0.608514** | **0.552958** | **0.590923** | **0.101326** | **1.853722** |
| **2404** | **0.23117** | **0.254458** | **0.984758** | **0.400043** | **1.870429** |
| **2581** | **0.545522** | **0.19027** | **0.595865** | **0.551178** | **1.882835** |
| **2347** | **0.724948** | **0.682072** | **0.228308** | **0.252987** | **1.888315** |
| **2958** | **0.64568** | **0.354174** | **0.552877** | **0.347838** | **1.90057** |
| **2571** | **0.461495** | **0.724186** | **0.427267** | **0.293574** | **1.906522** |
| **2368** | **0.253972** | **0.442238** | **0.613342** | **0.600369** | **1.909922** |
| **2362** | **0.141751** | **0.191202** | **1.314281** | **0.267706** | **1.91494** |
| **2446** | **0.089231** | **0.406585** | **1.201906** | **0.223678** | **1.9214** |
| **2947** | **0.275542** | **0.246366** | **0.855931** | **0.566514** | **1.944353** |
| **2455** | **1.002766** | **0.199828** | **0.434014** | **0.308346** | **1.944953** |
| **2349** | **0.060691** | **0.141604** | **0.630429** | **1.115654** | **1.948378** |
| **2930** | **0.641589** | **0.858584** | **0.32235** | **0.138765** | **1.961288** |
| **2556** | **0.793295** | **0.229305** | **0.29786** | **0.648731** | **1.969191** |
| **2378** | **0.093424** | **0.166318** | **0.337344** | **1.377973** | **1.975058** |
| **2542** | **0.404767** | **0.314378** | **0.79768** | **0.459818** | **1.976643** |
| **2548** | **0.513859** | **0.977934** | **0.294219** | **0.201572** | **1.987584** |
| **2366** | **0.531135** | **0.400598** | **0.302866** | **0.755505** | **1.990104** |
| **2811** | **0.423536** | **0.396974** | **0.895085** | **0.277388** | **1.992982** |
| **2532** | **1.014589** | **0.31282** | **0.41259** | **0.256114** | **1.996114** |
| **2552** | **0** | **0** | **0.810497** | **1.190243** | **2.00074** |
| **2336** | **1.650896** | **0.266734** | **0.047906** | **0.036095** | **2.00163** |
| **2454** | **1.129453** | **0.090546** | **0.184607** | **0.60594** | **2.010547** |
| **2940** | **0.426445** | **0.305464** | **0.927882** | **0.350883** | **2.010675** |
| **2533** | **0.37706** | **0.835602** | **0.289547** | **0.512159** | **2.014369** |
| **2428** | **0.632624** | **0.627921** | **0.206235** | **0.557747** | **2.024527** |
| **2613** | **0.695691** | **0.169189** | **0.799704** | **0.36979** | **2.034374** |
| **2852** | **0.657774** | **0.297805** | **0.503639** | **0.584239** | **2.043458** |
| **2466** | **0.378687** | **0.580441** | **0.440047** | **0.645259** | **2.044435** |
| **2438** | **0.154442** | **1.179243** | **0.54343** | **0.170128** | **2.047243** |
| **2401** | **0.156811** | **0.521486** | **0.402371** | **0.968294** | **2.048962** |
| **2929** | **0.414178** | **0.306775** | **0.194403** | **1.149874** | **2.06523** |
| **2657** | **1.191936** | **0.275867** | **0.4758** | **0.126874** | **2.070477** |
| **2473** | **0.146529** | **0.604353** | **1.178792** | **0.150669** | **2.080342** |
| **2418** | **0.056198** | **0.378199** | **1.429597** | **0.238496** | **2.10249** |
| **2569** | **0.989304** | **0.679131** | **0.074949** | **0.359415** | **2.102798** |
| **2809** | **0.318687** | **0.637997** | **0.037413** | **1.114533** | **2.108629** |
| **2348** | **0.113842** | **0** | **0.888421** | **1.115462** | **2.117726** |
| **2407** | **1.134749** | **0.171109** | **0.327006** | **0.493035** | **2.125899** |
| **2575** | **0.445633** | **1.134548** | **0.231959** | **0.324384** | **2.136525** |
| **2673** | **1.294697** | **0.374048** | **0.442164** | **0.037547** | **2.148457** |
| **2373** | **0.213239** | **0.560811** | **0.4459** | **0.938604** | **2.158554** |
| **2760** | **0** | **0.007486** | **0.28581** | **1.874098** | **2.167394** |
| **2356** | **0.442085** | **1.104363** | **0.484089** | **0.142496** | **2.173033** |
| **2839** | **0.938884** | **0.516431** | **0.367301** | **0.355801** | **2.178418** |
| **2271** | **0.21173** | **0.16063** | **0.647616** | **1.159272** | **2.179247** |
| **2634** | **0.374961** | **0.21206** | **0.863673** | **0.732825** | **2.183518** |
| **2659** | **1.589007** | **0.112003** | **0.28465** | **0.200268** | **2.185927** |
| **2465** | **0** | **0.508297** | **0.502608** | **1.177177** | **2.188082** |
| **2844** | **0.1014** | **1.157821** | **0.280763** | **0.659736** | **2.19972** |
| **2567** | **1.115589** | **0.354406** | **0.49228** | **0.253124** | **2.215398** |
| **2636** | **0.155052** | **0.370799** | **1.44114** | **0.248705** | **2.215697** |
| **2688** | **0** | **1.583723** | **0.130143** | **0.504912** | **2.218779** |
| **2365** | **0.085222** | **1.097785** | **0.234275** | **0.810347** | **2.227629** |
| **2363** | **0.114985** | **0.095152** | **0.278719** | **1.742738** | **2.231594** |
| **2604** | **0.350221** | **0.472738** | **0.780117** | **0.629754** | **2.23283** |
| **2537** | **0.27045** | **0.806736** | **0.695873** | **0.462369** | **2.235429** |
| **2462** | **0** | **0.423711** | **1.149061** | **0.684294** | **2.257066** |
| **2440** | **0.254189** | **0.050523** | **0.900349** | **1.070552** | **2.275613** |
| **2560** | **1.018998** | **0.332996** | **0.492469** | **0.437034** | **2.281497** |
| **2339** | **0.106376** | **1.233398** | **0.672646** | **0.273165** | **2.285586** |
| **2384** | **1.035207** | **0.121201** | **1.143034** | **0.001458** | **2.300899** |
| **2501** | **0.451894** | **1.008979** | **0.359819** | **0.480725** | **2.301417** |
| **2482** | **0.298341** | **0.766363** | **0.013247** | **1.23976** | **2.317711** |
| **2696** | **0.211167** | **1.068443** | **0.703583** | **0.335649** | **2.318843** |
| **2483** | **0.428655** | **1.560459** | **0.20151** | **0.145682** | **2.336306** |
| **2896** | **0.492354** | **0.530429** | **1.310709** | **0.006293** | **2.339785** |
| **2771** | **0.867158** | **0.529031** | **0.685605** | **0.261224** | **2.343019** |
| **2370** | **0.098117** | **0.145154** | **0.096594** | **2.02104** | **2.360906** |
| **2377** | **0.474877** | **0.242109** | **0.58786** | **1.059532** | **2.364379** |
| **2411** | **1.327431** | **0.070415** | **0.132722** | **0.841457** | **2.372025** |
| **2758** | **0** | **0.103382** | **2.102193** | **0.166927** | **2.372502** |
| **2922** | **1.614951** | **0.336082** | **0.041503** | **0.381194** | **2.373728** |
| **2402** | **0.951545** | **1.031152** | **0.283216** | **0.107936** | **2.373849** |
| **2305** | **0.75736** | **1.077195** | **0.46943** | **0.077795** | **2.38178** |
| **2497** | **0.34964** | **1.129116** | **0.477866** | **0.428751** | **2.385373** |
| **2913** | **0.815376** | **0.606447** | **0.287847** | **0.678492** | **2.388162** |
| **2390** | **0.585182** | **0.264576** | **0.606091** | **0.938199** | **2.394048** |
| **2250** | **0.093337** | **0.287196** | **1.240972** | **0.773311** | **2.394815** |
| **2429** | **0.132733** | **0.520074** | **0.999661** | **0.752309** | **2.404778** |
| **2475** | **0.179415** | **0.615974** | **1.253444** | **0.369167** | **2.418** |
| **2388** | **0.174657** | **0.343442** | **0.568968** | **1.337097** | **2.424163** |
| **2549** | **0.619095** | **0.099582** | **0.538514** | **1.183137** | **2.440328** |
| **2541** | **0.049464** | **0.467308** | **0.240273** | **1.687707** | **2.444752** |
| **2578** | **0.176606** | **1.564527** | **0.434505** | **0.276941** | **2.452579** |
| **2949** | **0.444744** | **0.373231** | **0.483852** | **1.1524** | **2.454227** |
| **2380** | **0.221627** | **0.754538** | **1.350433** | **0.140705** | **2.467303** |
| **2960** | **1.534363** | **0.114576** | **0.192588** | **0.643465** | **2.484992** |
| **2690** | **0.622979** | **0.046269** | **1.339104** | **0.495211** | **2.503563** |
| **2327** | **0.1328** | **0.590456** | **0.41555** | **1.365634** | **2.50444** |
| **2576** | **0.682481** | **0.742829** | **0.25657** | **0.823402** | **2.505282** |
| **2491** | **0.174392** | **0.460697** | **0.738834** | **1.133394** | **2.507316** |
| **2570** | **0.140467** | **1.07467** | **0.687502** | **0.608878** | **2.511517** |
| **2775** | **1.299123** | **0.687567** | **0.140192** | **0.385084** | **2.511967** |
| **2579** | **0.265231** | **0.965814** | **0.833333** | **0.473383** | **2.537761** |
| **2709** | **0** | **0.109951** | **1.38866** | **1.04177** | **2.540381** |
| **2903** | **1.096105** | **0.493658** | **0.75144** | **0.204226** | **2.545429** |
| **2450** | **0.214813** | **0.787595** | **1.384206** | **0.170969** | **2.557584** |
| **2851** | **0.208484** | **0.167481** | **0.430503** | **1.753556** | **2.560025** |
| **2383** | **0.048924** | **0.288955** | **2.007346** | **0.237829** | **2.583055** |
| **2371** | **0.534277** | **0.762651** | **0.892764** | **0.404278** | **2.59397** |
| **2759** | **0** | **0** | **1.965544** | **0.63033** | **2.595874** |
| **2817** | **0.005356** | **0.357722** | **0.140907** | **2.093877** | **2.597862** |
| **2442** | **1.083055** | **0.13546** | **1.174898** | **0.205286** | **2.598699** |
| **2823** | **0.003272** | **0.516486** | **1.736826** | **0.34433** | **2.600914** |
| **2876** | **1.720107** | **0.109929** | **0.477403** | **0.305643** | **2.613081** |
| **2631** | **1.325918** | **0.127146** | **0.742973** | **0.419693** | **2.615731** |
| **2853** | **0.421391** | **1.157036** | **0.958727** | **0.079266** | **2.61642** |
| **2484** | **0.507149** | **0.825035** | **0.5413** | **0.751667** | **2.62515** |
| **2472** | **0.345571** | **1.862283** | **0.121787** | **0.301294** | **2.630935** |
| **2936** | **0.221946** | **0.265064** | **0.638636** | **1.508461** | **2.634108** |
| **2606** | **0.87735** | **0.174413** | **0.676067** | **0.90816** | **2.635991** |
| **2519** | **0.708668** | **0.154404** | **1.488684** | **0.289525** | **2.641281** |
| **2632** | **0.424483** | **2.121128** | **0.004986** | **0.117913** | **2.668509** |
| **2826** | **0.671259** | **0.249867** | **1.445428** | **0.323406** | **2.68996** |
| **2959** | **0.32313** | **0.826346** | **1.338576** | **0.204437** | **2.692488** |
| **2708** | **0.434806** | **0.067032** | **1.264647** | **0.940318** | **2.706803** |
| **2477** | **0.69976** | **1.021806** | **0.579727** | **0.416099** | **2.717392** |
| **2854** | **0.908759** | **0.462629** | **1.309145** | **0.039986** | **2.720519** |
| **2524** | **0.697453** | **1.176794** | **0.440475** | **0.412709** | **2.72743** |
| **2612** | **0** | **0.554534** | **0.224019** | **1.949856** | **2.72841** |
| **2910** | **0.724572** | **0.42936** | **0.096231** | **1.4818** | **2.731963** |
| **2340** | **0.405881** | **0.608642** | **1.01904** | **0.699204** | **2.732767** |
| **2593** | **0.686376** | **1.081858** | **0.229865** | **0.739951** | **2.73805** |
| **2595** | **0.679911** | **0.937294** | **0.932758** | **0.21951** | **2.769474** |
| **2591** | **0.402885** | **1.205658** | **0.276198** | **0.889548** | **2.774289** |
| **2879** | **0.273402** | **1.085358** | **0.994306** | **0.424909** | **2.777975** |
| **2692** | **0.902044** | **0.25634** | **0.505862** | **1.123678** | **2.787924** |
| **2599** | **1.009643** | **1.324581** | **0.190151** | **0.276065** | **2.800441** |
| **2294** | **1.934737** | **0.429833** | **0.184398** | **0.255376** | **2.804343** |
| **2685** | **0.139117** | **0.511074** | **1.324108** | **0.84292** | **2.817219** |
| **2766** | **1.092352** | **0.919182** | **0.506329** | **0.316688** | **2.83455** |
| **2590** | **1.400176** | **0.049221** | **1.113328** | **0.287011** | **2.849735** |
| **2503** | **0.264596** | **0.42048** | **2.168865** | **0.012424** | **2.866365** |
| **2951** | **0.608249** | **1.179258** | **0.323681** | **0.776196** | **2.887384** |
| **2768** | **0** | **0.003208** | **2.445999** | **0.441941** | **2.891149** |
| **2433** | **1.241332** | **0.563021** | **0.572596** | **0.514215** | **2.891164** |
| **2885** | **0.894135** | **0.696733** | **0.993438** | **0.307704** | **2.89201** |
| **2300** | **0** | **1.164009** | **1.628118** | **0.10412** | **2.896247** |
| **2658** | **0.712881** | **0.738007** | **0.978406** | **0.474859** | **2.904153** |
| **2420** | **0.935688** | **0.858583** | **0.832184** | **0.279573** | **2.906028** |
| **2800** | **0.602346** | **0** | **1.185292** | **1.123653** | **2.91129** |
| **2359** | **0.982892** | **0.157723** | **0.734579** | **1.044334** | **2.919529** |
| **2812** | **0** | **0.652157** | **1.661421** | **0.619108** | **2.932686** |
| **2906** | **0.334544** | **0.188858** | **1.736906** | **0.673633** | **2.933941** |
| **2587** | **0.106294** | **0.090861** | **0.592553** | **2.144548** | **2.934256** |
| **2492** | **0.529449** | **1.360578** | **0.590641** | **0.453655** | **2.934322** |
| **2534** | **1.081769** | **0.795714** | **0.417526** | **0.645166** | **2.940176** |
| **2860** | **0.154511** | **0.427595** | **1.795815** | **0.570522** | **2.948443** |
| **2950** | **1.258118** | **0.282895** | **0.44058** | **0.980222** | **2.961815** |
| **2507** | **1.692323** | **0.48419** | **0.270763** | **0.538269** | **2.985546** |
| **2880** | **0.432286** | **2.088868** | **0.449987** | **0.027319** | **2.99846** |
| **2833** | **2.476407** | **0.200458** | **0.073346** | **0.259536** | **3.009747** |
| **2605** | **0.425602** | **0.576104** | **1.083128** | **0.935922** | **3.020756** |
| **2730** | **1.044371** | **0.480054** | **1.027419** | **0.478962** | **3.030806** |
| **2810** | **0.327821** | **0.046971** | **0.205921** | **2.452597** | **3.03331** |
| **2516** | **0.698551** | **0.474628** | **1.187512** | **0.684446** | **3.045138** |
| **2874** | **0.646383** | **0.06213** | **1.724719** | **0.612797** | **3.046029** |
| **2270** | **0.11752** | **1.190269** | **0.743102** | **0.999628** | **3.050519** |
| **2330** | **2.559777** | **0.006345** | **0.002877** | **0.488643** | **3.057642** |
| **2640** | **0.976442** | **1.268751** | **0.216357** | **0.609949** | **3.071499** |
| **2836** | **0.666574** | **0** | **2.143293** | **0.262242** | **3.072108** |
| **2897** | **0.818057** | **0.272647** | **1.4404** | **0.555496** | **3.086601** |
| **2773** | **0.616996** | **0.159014** | **0.710161** | **1.611808** | **3.097979** |
| **2508** | **0.483974** | **2.026065** | **0.219587** | **0.373237** | **3.102863** |
| **2820** | **0.577671** | **0.142407** | **0.993802** | **1.399273** | **3.113154** |
| **2847** | **1.246954** | **0.987752** | **0.302332** | **0.581503** | **3.118541** |
| **2405** | **0.850476** | **0.957375** | **0.401761** | **0.910384** | **3.119997** |
| **2495** | **0.195216** | **0.264658** | **1.098372** | **1.587077** | **3.145323** |
| **2439** | **0.611149** | **1.017335** | **0.577832** | **0.940167** | **3.146483** |
| **2319** | **0.384219** | **2.281761** | **0.327719** | **0.163335** | **3.157035** |
| **2739** | **0.320388** | **1.121503** | **0.312989** | **1.403963** | **3.158843** |
| **2866** | **0.113986** | **0.796302** | **1.301998** | **0.95649** | **3.168777** |
| **2727** | **1.874804** | **0** | **0.924169** | **0.372811** | **3.171784** |
| **2471** | **0.838134** | **0.315647** | **1.288737** | **0.742025** | **3.184543** |
| **2470** | **0.872472** | **0.941246** | **0.650704** | **0.723046** | **3.187468** |
| **2444** | **1.861143** | **0.191064** | **0.856908** | **0.292462** | **3.201577** |
| **2321** | **1.291482** | **0.265515** | **1.081455** | **0.568707** | **3.207159** |
| **2676** | **0.044427** | **0** | **2.576741** | **0.59741** | **3.218579** |
| **2915** | **0.566564** | **0.848076** | **0.130242** | **1.685248** | **3.230131** |
| **2943** | **0.65733** | **0.696995** | **1.251581** | **0.630605** | **3.23651** |
| **2248** | **0** | **0.052495** | **0.784543** | **2.415381** | **3.252419** |
| **2796** | **0.372652** | **0.83848** | **1.751137** | **0.292576** | **3.254846** |
| **2253** | **0.037016** | **0.398667** | **1.373734** | **1.447784** | **3.257201** |
| **2945** | **0.070404** | **0.007698** | **2.623292** | **0.556652** | **3.258046** |
| **2699** | **0.714206** | **0.390624** | **1.290591** | **0.872785** | **3.268206** |
| **2449** | **0.218043** | **0.500885** | **1.527406** | **1.03217** | **3.278505** |
| **2289** | **0.149203** | **1.795046** | **0.397933** | **0.947427** | **3.289609** |
| **2435** | **0.885247** | **0.056613** | **2.069323** | **0.289824** | **3.301007** |
| **2312** | **0.76752** | **0.338643** | **0.740425** | **1.458893** | **3.305481** |
| **2500** | **1.048573** | **0.534611** | **1.075225** | **0.647977** | **3.306386** |
| **2558** | **0.038586** | **1.066154** | **0.691196** | **1.516812** | **3.312748** |
| **2881** | **1.253246** | **0.014792** | **1.084875** | **0.963503** | **3.316417** |
| **2434** | **0** | **1.487775** | **1.658926** | **0.174679** | **3.32138** |
| **2467** | **0.272411** | **1.077583** | **0.813703** | **1.159783** | **3.32348** |
| **2858** | **0.21027** | **1.739736** | **0.054125** | **1.321671** | **3.325802** |
| **2596** | **1.174151** | **0.417654** | **0.580184** | **1.169655** | **3.341644** |
| **2487** | **0.29765** | **0.975431** | **1.602943** | **0.469963** | **3.345987** |
| **2608** | **0.785129** | **1.180948** | **0.943813** | **0.436742** | **3.346633** |
| **2870** | **1.057279** | **1.511165** | **0.467668** | **0.32461** | **3.360721** |
| **2827** | **0.943154** | **1.996329** | **0.289618** | **0.143307** | **3.372409** |
| **2600** | **0.942805** | **0.492727** | **0.534917** | **1.40489** | **3.37534** |
| **2686** | **0.187591** | **0.39484** | **1.768102** | **1.029765** | **3.380297** |
| **2873** | **0** | **0.14291** | **0.955834** | **2.285344** | **3.384088** |
| **2954** | **0.905963** | **0.109908** | **1.268037** | **1.110055** | **3.393962** |
| **2921** | **1.262414** | **0.590617** | **0.534419** | **1.018866** | **3.406316** |
| **2932** | **0.850754** | **1.216013** | **0.116125** | **1.22488** | **3.407773** |
| **2784** | **2.018725** | **0.274434** | **0.136008** | **0.999514** | **3.42868** |
| **2857** | **1.195941** | **1.781942** | **0.33593** | **0.131662** | **3.445475** |
| **2701** | **0.83222** | **0.72931** | **1.019758** | **0.872047** | **3.453336** |
| **2804** | **0.495346** | **1.609722** | **0.60951** | **0.76743** | **3.482008** |
| **2933** | **0.395051** | **1.234306** | **1.376653** | **0.485877** | **3.491888** |
| **2957** | **0.883401** | **1.396456** | **0.830193** | **0.385202** | **3.495252** |
| **2740** | **0.094724** | **1.340266** | **0.6718** | **1.398901** | **3.505691** |
| **2783** | **0.054238** | **1.608897** | **1.130616** | **0.722014** | **3.515765** |
| **2583** | **0.724292** | **1.448492** | **1.046095** | **0.300162** | **3.519042** |
| **2240** | **0.091999** | **1.002321** | **1.293335** | **1.136083** | **3.523738** |
| **2544** | **0.673099** | **1.096828** | **1.511712** | **0.26762** | **3.549259** |
| **2610** | **0.574648** | **0.674287** | **1.967785** | **0.353725** | **3.570445** |
| **2527** | **1.157865** | **0.662134** | **0.808842** | **0.956485** | **3.585327** |
| **2927** | **1.263178** | **0.702857** | **0.95453** | **0.669286** | **3.58985** |
| **2656** | **1.056606** | **1.344103** | **1.025041** | **0.174937** | **3.600687** |
| **2547** | **0.220132** | **0.389692** | **1.961369** | **1.03023** | **3.601423** |
| **2898** | **0.844347** | **0.053826** | **1.775087** | **0.928985** | **3.602246** |
| **2872** | **1.871882** | **0.136868** | **0.218835** | **1.376003** | **3.603589** |
| **2311** | **0.603066** | **0.990731** | **1.454258** | **0.564423** | **3.612478** |
| **2607** | **2.161859** | **1.3231** | **0** | **0.140323** | **3.625282** |
| **2726** | **2.690052** | **0.451049** | **0.439778** | **0.050539** | **3.631418** |
| **2478** | **0.157754** | **0.872027** | **1.619303** | **0.982693** | **3.631777** |
| **2831** | **0.08072** | **0.975378** | **1.172352** | **1.414764** | **3.643214** |
| **2284** | **1.261587** | **0** | **2.225393** | **0.160921** | **3.647901** |
| **2891** | **0.786903** | **0.040726** | **0.802059** | **2.025588** | **3.655276** |
| **2494** | **2.611778** | **0.313818** | **0.592191** | **0.137518** | **3.655305** |
| **2627** | **1.768379** | **0.963499** | **0.160998** | **0.771734** | **3.66461** |
| **2868** | **1.82658** | **0.789256** | **0.646195** | **0.416249** | **3.67828** |
| **2752** | **0** | **0** | **0.044367** | **3.636947** | **3.681314** |
| **2481** | **0.618931** | **1.251558** | **1.371233** | **0.44288** | **3.684603** |
| **2924** | **2.256003** | **0.166114** | **0.492025** | **0.778859** | **3.693001** |
| **2707** | **1.542038** | **0.266912** | **0.355818** | **1.547242** | **3.71201** |
| **2565** | **0.673954** | **0.888825** | **1.033368** | **1.117322** | **3.713469** |
| **2887** | **1.641034** | **1.420841** | **0.471717** | **0.192201** | **3.725793** |
| **2806** | **1.143784** | **0.695195** | **1.595006** | **0.3124** | **3.746386** |
| **2710** | **0** | **0.842823** | **0.663308** | **2.271368** | **3.7775** |
| **2902** | **0.050625** | **0.961534** | **0.168742** | **2.602675** | **3.783576** |
| **2419** | **1.445223** | **0.320896** | **1.910177** | **0.138585** | **3.814881** |
| **2525** | **0.440102** | **1.365071** | **1.300733** | **0.723577** | **3.829483** |
| **2808** | **0.48159** | **1.421818** | **0.876285** | **1.057775** | **3.837468** |
| **2314** | **0.616153** | **0.507328** | **2.046629** | **0.673812** | **3.843922** |
| **2842** | **0.763533** | **1.429609** | **0.841182** | **0.819686** | **3.854011** |
| **2705** | **0.489019** | **2.461339** | **0.394579** | **0.510697** | **3.855633** |
| **2445** | **0.009569** | **0.779632** | **2.71394** | **0.352501** | **3.855641** |
| **2272** | **1.497868** | **1.610787** | **0.015756** | **0.735152** | **3.859563** |
| **2580** | **2.034218** | **0.577762** | **0.309242** | **0.938746** | **3.859967** |
| **2588** | **1.742046** | **1.118902** | **0.76726** | **0.234344** | **3.862552** |
| **2288** | **0.8046** | **0.770073** | **1.9931** | **0.304209** | **3.871982** |
| **2886** | **0.704895** | **0.599633** | **0.636703** | **1.938564** | **3.879794** |
| **2956** | **0.38216** | **2.080602** | **0.579427** | **0.856951** | **3.899139** |
| **2474** | **0.759953** | **1.625318** | **1.044963** | **0.471408** | **3.901642** |
| **2463** | **0.265101** | **1.397866** | **0.272448** | **1.9685** | **3.903916** |
| **2719** | **1.650058** | **0.314923** | **1.939047** | **0** | **3.904027** |
| **2923** | **0.774479** | **1.071361** | **0.179595** | **1.916124** | **3.94156** |
| **2258** | **0** | **0** | **0.102389** | **3.843831** | **3.94622** |
| **2637** | **0.638444** | **1.252739** | **1.83853** | **0.23345** | **3.963163** |
| **2905** | **1.01936** | **1.50077** | **0.133557** | **1.31256** | **3.966247** |
| **2431** | **0.46111** | **0.270341** | **0.592914** | **2.654908** | **3.979274** |
| **2456** | **0.19879** | **1.612619** | **1.843137** | **0.355586** | **4.010132** |
| **2818** | **0.722297** | **1.295621** | **0.498211** | **1.50787** | **4.024** |
| **2278** | **0.280655** | **0.635806** | **0.269609** | **2.84357** | **4.02964** |
| **2849** | **0.787507** | **0.59506** | **0.700966** | **1.958888** | **4.042421** |
| **2551** | **0.532783** | **1.546962** | **0.521424** | **1.442023** | **4.043192** |
| **2716** | **0.884627** | **1.680342** | **0.583474** | **0.90732** | **4.055762** |
| **2476** | **1.11301** | **1.26864** | **0.874079** | **0.804481** | **4.060209** |
| **2855** | **0.432946** | **1.118378** | **1.893954** | **0.615561** | **4.060839** |
| **2287** | **0** | **0** | **0.383088** | **3.708513** | **4.091601** |
| **2469** | **1.474106** | **0.42491** | **1.729608** | **0.467958** | **4.096583** |
| **2695** | **1.962266** | **0.001663** | **0.809537** | **1.324999** | **4.098465** |
| **2372** | **0.838697** | **0.144096** | **1.985034** | **1.170071** | **4.137898** |
| **2244** | **1.745029** | **1.340812** | **1.033636** | **0.022405** | **4.141882** |
| **2765** | **0.124239** | **0.154196** | **1.828453** | **2.054363** | **4.161252** |
| **2254** | **0.811647** | **0.478806** | **2.83069** | **0.041641** | **4.162784** |
| **2615** | **2.263812** | **0.659327** | **0.78372** | **0.458331** | **4.16519** |
| **2255** | **0.762422** | **2.173102** | **1.027659** | **0.20567** | **4.168853** |
| **2918** | **1.067748** | **1.871618** | **0.045696** | **1.208353** | **4.193416** |
| **2265** | **2.057772** | **1.978356** | **0.029586** | **0.185094** | **4.250808** |
| **2715** | **2.118506** | **0.493666** | **0.773498** | **0.870763** | **4.256434** |
| **2655** | **0.975216** | **0.153584** | **2.039176** | **1.100827** | **4.268802** |
| **2303** | **0.381665** | **2.191874** | **1.047918** | **0.649546** | **4.271004** |
| **2263** | **0.169547** | **2.71614** | **0.840633** | **0.544744** | **4.271065** |
| **2624** | **0.02572** | **0.567881** | **1.642488** | **2.044789** | **4.280878** |
| **2529** | **0.930996** | **1.723346** | **0.838504** | **0.80292** | **4.295766** |
| **2512** | **1.742461** | **1.356153** | **0.670285** | **0.549348** | **4.318248** |
| **2689** | **1.093448** | **0.193762** | **1.945819** | **1.086871** | **4.3199** |
| **2351** | **2.336269** | **0.450563** | **1.059853** | **0.500507** | **4.347192** |
| **2865** | **0.773338** | **0.270855** | **1.623719** | **1.722469** | **4.390381** |
| **2509** | **0.335843** | **0.332245** | **1.273746** | **2.482522** | **4.424356** |
| **2742** | **0.457107** | **0.596538** | **0.190336** | **3.181761** | **4.425742** |
| **2877** | **1.756053** | **0.917913** | **0.644969** | **1.119217** | **4.438153** |
| **2480** | **2.904961** | **0.654033** | **0.610034** | **0.269399** | **4.438428** |
| **2647** | **1.462575** | **0.433643** | **1.45835** | **1.094359** | **4.448927** |
| **2893** | **0.819472** | **2.34139** | **0.956589** | **0.342372** | **4.459823** |
| **2931** | **2.217377** | **1.310751** | **0.317199** | **0.6273** | **4.472627** |
| **2917** | **2.471612** | **0.331661** | **1.603714** | **0.081183** | **4.488171** |
| **2611** | **0.539473** | **1.713788** | **1.633555** | **0.621638** | **4.508455** |
| **2513** | **0.24859** | **2.171325** | **1.347141** | **0.745331** | **4.512386** |
| **2387** | **0.36027** | **0.705018** | **2.148238** | **1.299872** | **4.513397** |
| **2538** | **0.726746** | **1.037903** | **2.0023** | **0.762423** | **4.529372** |
| **2422** | **2.366065** | **0.849035** | **0.346139** | **0.987479** | **4.548717** |
| **2502** | **2.350629** | **0.249116** | **0.531921** | **1.43963** | **4.571296** |
| **2546** | **2.9117** | **0.158463** | **1.209431** | **0.304696** | **4.584291** |
| **2629** | **0.522781** | **1.048576** | **0.968155** | **2.046107** | **4.585619** |
| **2425** | **0.268247** | **2.351296** | **0.83631** | **1.144469** | **4.600322** |
| **2436** | **1.197715** | **0.57094** | **2.053194** | **0.809676** | **4.631525** |
| **2925** | **1.526677** | **0.568074** | **1.896942** | **0.654464** | **4.646157** |
| **2239** | **3.451201** | **0** | **0.802257** | **0.399391** | **4.652849** |
| **2275** | **1.692303** | **0.28207** | **1.716051** | **0.990294** | **4.680719** |
| **2667** | **1.731128** | **0.60735** | **2.317099** | **0.037809** | **4.693386** |
| **2458** | **0.261827** | **0.944515** | **0.615895** | **2.886701** | **4.708938** |
| **2900** | **1.626145** | **2.135005** | **0.457512** | **0.502182** | **4.720843** |
| **2486** | **1.656098** | **1.127544** | **0.436516** | **1.50801** | **4.728168** |
| **2633** | **2.386824** | **0.913598** | **1.384456** | **0.047152** | **4.73203** |
| **2845** | **0.521894** | **3.823186** | **0** | **0.393318** | **4.738398** |
| **2757** | **1.467957** | **1.373592** | **0.403574** | **1.518895** | **4.764018** |
| **2944** | **1.072854** | **0.568563** | **2.145282** | **0.999756** | **4.786455** |
| **2550** | **0.157032** | **1.961027** | **0.779399** | **1.899645** | **4.797104** |
| **2920** | **1.022593** | **0.476097** | **1.735831** | **1.586067** | **4.820588** |
| **2643** | **0.137223** | **1.399201** | **0.827213** | **2.46169** | **4.825326** |
| **2644** | **0.030564** | **0.795826** | **2.544496** | **1.475129** | **4.846014** |
| **2242** | **1.77706** | **0.889625** | **0.25865** | **1.924448** | **4.849783** |
| **2899** | **2.639707** | **0.638841** | **0.874062** | **0.716415** | **4.869025** |
| **2938** | **0.183271** | **2.817753** | **0.488971** | **1.381749** | **4.871743** |
| **2315** | **0.740436** | **1.311781** | **0.640166** | **2.18088** | **4.873263** |
| **2269** | **0.612592** | **1.048393** | **1.998974** | **1.233455** | **4.893414** |
| **2259** | **3.754996** | **0** | **0.780828** | **0.392897** | **4.928721** |
| **2770** | **1.715946** | **0.924277** | **1.696159** | **0.605572** | **4.941954** |
| **2840** | **0.264993** | **2.110674** | **0.523884** | **2.077206** | **4.976756** |
| **2749** | **0** | **2.028693** | **0.459038** | **2.493258** | **4.980989** |
| **2794** | **1.388363** | **0.697478** | **0.388611** | **2.507996** | **4.982449** |
| **2530** | **1.916634** | **1.461464** | **1.550606** | **0.059429** | **4.988132** |
| **2668** | **0.238326** | **0.117916** | **0.079217** | **4.561806** | **4.997264** |
| **2928** | **2.436036** | **0.961538** | **0.158808** | **1.464944** | **5.021326** |
| **2746** | **0.48958** | **1.516368** | **1.779193** | **1.252143** | **5.037285** |
| **2722** | **0** | **0** | **0.698992** | **4.338873** | **5.037865** |
| **2299** | **0.380178** | **3.353856** | **0.626664** | **0.706286** | **5.066983** |
| **2641** | **0.124514** | **2.301753** | **0.628608** | **2.024289** | **5.079164** |
| **2249** | **0.888651** | **2.002809** | **0.771103** | **1.427562** | **5.090125** |
| **2681** | **0.207108** | **3.346413** | **0.763858** | **0.780788** | **5.098165** |
| **2520** | **0.232665** | **1.014881** | **1.792002** | **2.061714** | **5.101262** |
| **2262** | **0** | **0.00539** | **4.209373** | **0.905414** | **5.120177** |
| **2424** | **2.0185** | **1.462764** | **0.768137** | **0.881945** | **5.131346** |
| **2856** | **0.548115** | **0.674569** | **1.319066** | **2.590084** | **5.131834** |
| **2298** | **2.079283** | **1.994794** | **0.552587** | **0.535604** | **5.162268** |
| **2528** | **0.346927** | **3.772174** | **0.515171** | **0.556095** | **5.190367** |
| **2697** | **0.452974** | **1.987003** | **0.504455** | **2.29526** | **5.239692** |
| **2683** | **0.427104** | **2.158931** | **1.009699** | **1.687746** | **5.28348** |
| **2747** | **0.598363** | **0.587451** | **0.596529** | **3.501375** | **5.283718** |
| **2304** | **1.312304** | **1.813893** | **1.977577** | **0.221196** | **5.32497** |
| **2427** | **0.481441** | **2.074797** | **1.17487** | **1.625385** | **5.356493** |
| **2948** | **2.229177** | **0.574703** | **1.738212** | **0.825806** | **5.367897** |
| **2586** | **0.733797** | **1.064951** | **0.419991** | **3.152221** | **5.370959** |
| **2408** | **0.915583** | **2.786736** | **1.538323** | **0.130402** | **5.371045** |
| **2320** | **2.090958** | **0.840637** | **1.211552** | **1.259935** | **5.403082** |
| **2441** | **1.20677** | **1.281155** | **0.668073** | **2.248548** | **5.404546** |
| **2316** | **0.682856** | **0.537013** | **1.357986** | **2.829462** | **5.407317** |
| **2297** | **1.018575** | **2.098834** | **0.836052** | **1.522868** | **5.476329** |
| **2942** | **0.484685** | **1.695262** | **2.102649** | **1.218932** | **5.501529** |
| **2761** | **1.138245** | **2.268637** | **0.924992** | **1.181461** | **5.513334** |
| **2700** | **0.878394** | **3.364868** | **0.520052** | **0.757242** | **5.520556** |
| **2630** | **0.846015** | **2.466707** | **1.633873** | **0.614291** | **5.560887** |
| **2687** | **0.092166** | **3.33726** | **0.004467** | **2.128646** | **5.562539** |
| **2598** | **3.865192** | **0.144248** | **1.059888** | **0.512382** | **5.581711** |
| **2518** | **0.042155** | **1.742327** | **0.956869** | **2.858207** | **5.599558** |
| **2521** | **0** | **0.015785** | **1.198975** | **4.410766** | **5.625526** |
| **2680** | **0.818027** | **3.177143** | **1.339538** | **0.361159** | **5.695867** |
| **2674** | **1.050458** | **1.63736** | **0.917391** | **2.183942** | **5.789151** |
| **2268** | **0.007952** | **0.475866** | **3.8697** | **1.490803** | **5.844321** |
| **2767** | **1.618446** | **0.225889** | **0.606283** | **3.416158** | **5.866776** |
| **2660** | **0.265079** | **2.382597** | **1.163026** | **2.099268** | **5.90997** |
| **2675** | **0.743744** | **3.24673** | **1.422245** | **0.56877** | **5.98149** |
| **2577** | **2.356539** | **2.351835** | **1.016934** | **0.280745** | **6.006053** |
| **2883** | **1.266302** | **1.405263** | **1.235587** | **2.108566** | **6.015718** |
| **2257** | **1.022393** | **1.445134** | **3.296532** | **0.269414** | **6.033473** |
| **2282** | **0.562751** | **1.58388** | **2.747633** | **1.188821** | **6.083086** |
| **2663** | **3.020849** | **1.607538** | **0.605627** | **0.860131** | **6.094144** |
| **2553** | **2.606177** | **1.563295** | **0.956079** | **0.983606** | **6.109157** |
| **2421** | **3.87785** | **0.324365** | **1.338627** | **0.570095** | **6.110938** |
| **2496** | **0.283336** | **0.521325** | **2.858787** | **2.4498** | **6.113248** |
| **2280** | **0.927588** | **1.746651** | **1.8458** | **1.597295** | **6.117335** |
| **2720** | **0.293991** | **3.801045** | **0.740448** | **1.317462** | **6.152947** |
| **2619** | **0.173953** | **3.139212** | **2.171493** | **0.717953** | **6.20261** |
| **2802** | **0.160471** | **3.580033** | **1.777425** | **0.698147** | **6.216076** |
| **2755** | **0.971074** | **2.067905** | **2.774783** | **0.41158** | **6.225342** |
| **2247** | **3.711992** | **1.828487** | **0.362863** | **0.357193** | **6.260537** |
| **2731** | **0.281138** | **3.128943** | **1.357563** | **1.50031** | **6.267954** |
| **2789** | **0.257385** | **2.697508** | **0.321122** | **3.106149** | **6.382164** |
| **2723** | **0.970348** | **0.606496** | **1.234118** | **3.57454** | **6.385501** |
| **2661** | **0.890476** | **2.681299** | **1.353883** | **1.472273** | **6.397931** |
| **2505** | **0.535698** | **2.560604** | **0.540517** | **2.767624** | **6.404444** |
| **2628** | **0.002478** | **2.495173** | **0.106355** | **3.818809** | **6.422815** |
| **2329** | **0.233257** | **3.964642** | **0.991486** | **1.279545** | **6.468929** |
| **2774** | **3.611388** | **1.553196** | **0.810793** | **0.497611** | **6.472988** |
| **2266** | **0.504572** | **1.405141** | **2.388489** | **2.196952** | **6.495154** |
| **2664** | **1.362638** | **1.346564** | **0.331328** | **3.456685** | **6.497215** |
| **2623** | **1.385885** | **0.195175** | **1.395198** | **3.573984** | **6.550243** |
| **2464** | **0.0466** | **4.631908** | **1.34616** | **0.530571** | **6.55524** |
| **2522** | **2.84452** | **2.204184** | **0.640346** | **0.881658** | **6.570708** |
| **2799** | **0.418343** | **2.527779** | **0.545468** | **3.169877** | **6.661466** |
| **2786** | **0.583413** | **2.865467** | **0.332974** | **2.89946** | **6.681314** |
| **2620** | **2.299175** | **2.107343** | **1.289763** | **1.012359** | **6.70864** |
| **2725** | **1.411242** | **0.262982** | **3.948434** | **1.096098** | **6.718756** |
| **2653** | **2.568896** | **1.962098** | **1.967802** | **0.23894** | **6.737736** |
| **2734** | **0** | **4.205452** | **1.707796** | **0.824653** | **6.737901** |
| **2597** | **0.663317** | **4.97502** | **0.359564** | **0.778872** | **6.776772** |
| **2781** | **3.256189** | **2.029139** | **1.18915** | **0.32298** | **6.797458** |
| **2451** | **0.240714** | **1.311897** | **2.145793** | **3.109637** | **6.808042** |
| **2651** | **3.069529** | **0.903097** | **2.407581** | **0.437687** | **6.817894** |
| **2279** | **1.771889** | **2.663787** | **0.452433** | **1.941068** | **6.829178** |
| **2622** | **0.848198** | **0.17072** | **4.547634** | **1.330314** | **6.896866** |
| **2241** | **0.03239** | **2.339776** | **3.763765** | **0.761661** | **6.897591** |
| **2792** | **0.653998** | **1.306054** | **1.961777** | **3.009888** | **6.931717** |
| **2614** | **0.256356** | **2.074812** | **2.570895** | **2.045226** | **6.947289** |
| **2621** | **1.224601** | **0.530474** | **1.619239** | **3.575631** | **6.949945** |
| **2331** | **0.022986** | **2.104255** | **2.726514** | **2.097743** | **6.951497** |
| **2286** | **0.515423** | **0.543983** | **5.133594** | **0.857297** | **7.050297** |
| **2738** | **0.760353** | **0.673647** | **4.455318** | **1.2039** | **7.093217** |
| **2816** | **3.715713** | **0.548977** | **2.330875** | **0.527887** | **7.123452** |
| **2801** | **1.404405** | **0.520817** | **2.204376** | **3.083954** | **7.213552** |
| **2302** | **6.666753** | **0.06403** | **0.292334** | **0.216842** | **7.239959** |
| **2326** | **3.577067** | **0.817746** | **0.744181** | **2.129881** | **7.268875** |
| **2798** | **0.718823** | **4.254161** | **2.300924** | **0.011304** | **7.285212** |
| **2457** | **0.827378** | **0.908466** | **3.36978** | **2.191696** | **7.297321** |
| **2568** | **4.5916** | **1.524003** | **0.302129** | **0.940016** | **7.357749** |
| **2679** | **2.736157** | **2.577316** | **0.870391** | **1.178221** | **7.362085** |
| **2277** | **1.204263** | **0.999815** | **0** | **5.205417** | **7.409495** |
| **2426** | **0.843298** | **1.2308** | **1.748224** | **3.622909** | **7.445231** |
| **2698** | **3.179344** | **1.069359** | **0.166531** | **3.041507** | **7.456741** |
| **2488** | **2.038751** | **3.378169** | **1.526117** | **0.560014** | **7.503051** |
| **2824** | **1.072917** | **4.701563** | **1.347934** | **0.423209** | **7.545623** |
| **2515** | **0.698846** | **3.689043** | **1.223621** | **1.982759** | **7.594269** |
| **2788** | **3.075928** | **1.709109** | **1.374249** | **1.489281** | **7.648567** |
| **2830** | **0.007692** | **0.129935** | **7.281402** | **0.240213** | **7.659242** |
| **2825** | **0.518494** | **2.341812** | **3.903186** | **0.988883** | **7.752375** |
| **2756** | **1.643784** | **0.582845** | **0.005483** | **5.522297** | **7.75441** |
| **2301** | **0.112513** | **1.243844** | **0.081955** | **6.339895** | **7.778206** |
| **2737** | **1.92082** | **1.23826** | **3.89568** | **0.800776** | **7.855536** |
| **2584** | **0.834254** | **4.000905** | **1.9794** | **1.071324** | **7.885883** |
| **2260** | **1.193882** | **3.458901** | **2.273113** | **0.961102** | **7.886998** |
| **2267** | **0.02514** | **1.312566** | **1.676944** | **4.939306** | **7.953956** |
| **2713** | **0** | **0.803841** | **1.200443** | **5.952451** | **7.956736** |
| **2777** | **0.884296** | **0.456733** | **2.155999** | **4.507551** | **8.004578** |
| **2273** | **2.764608** | **3.14717** | **1.813427** | **0.303755** | **8.028959** |
| **2724** | **1.372557** | **4.478695** | **1.969127** | **0.295582** | **8.115961** |
| **2702** | **1.579163** | **1.688822** | **4.490703** | **0.481347** | **8.240036** |
| **2603** | **3.478313** | **4.216949** | **0.257065** | **0.315696** | **8.268023** |
| **2328** | **0.145857** | **6.309081** | **1.013706** | **0.809266** | **8.27791** |
| **2677** | **0** | **0** | **1.68414** | **6.595049** | **8.279189** |
| **2423** | **1.596819** | **3.756314** | **2.376673** | **0.614794** | **8.3446** |
| **2735** | **1.070837** | **0.657341** | **1.574309** | **5.060364** | **8.362852** |
| **2317** | **3.333783** | **2.533509** | **0** | **2.540936** | **8.408229** |
| **2732** | **3.499855** | **0.201377** | **1.808126** | **2.942067** | **8.451426** |
| **2638** | **3.593224** | **0.580947** | **1.902549** | **2.388613** | **8.465334** |
| **2261** | **3.570568** | **2.005172** | **0.194146** | **2.967858** | **8.737743** |
| **2721** | **1.179135** | **0.509264** | **6.835259** | **0.258237** | **8.781896** |
| **2743** | **2.039485** | **0.763519** | **5.440141** | **0.560081** | **8.803226** |
| **2296** | **0.455985** | **2.659943** | **2.034053** | **3.664625** | **8.814605** |
| **2850** | **1.140381** | **6.446992** | **0.840501** | **0.438067** | **8.86594** |
| **2283** | **2.938858** | **1.214691** | **2.312287** | **2.408491** | **8.874327** |
| **2814** | **0.110413** | **0.142368** | **8.31385** | **0.357136** | **8.923766** |
| **2714** | **5.167299** | **1.691586** | **1.17452** | **1.133729** | **9.167134** |
| **2274** | **4.238172** | **1.40995** | **1.977982** | **1.657063** | **9.283167** |
| **2602** | **3.544736** | **0.90509** | **2.030318** | **2.834378** | **9.314521** |
| **2453** | **0.443999** | **0.304188** | **4.345566** | **4.291268** | **9.38502** |
| **2706** | **1.944133** | **3.72675** | **3.304427** | **0.533527** | **9.508837** |
| **2332** | **0.300811** | **7.160398** | **1.435792** | **0.725207** | **9.622209** |
| **2744** | **6.293804** | **1.643438** | **0.391353** | **1.367935** | **9.69653** |
| **2490** | **0.1235** | **6.738118** | **0.841717** | **2.021377** | **9.724712** |
| **2310** | **0.325897** | **4.158082** | **3.280974** | **2.064434** | **9.829386** |
| **2323** | **5.67428** | **0.289326** | **3.827468** | **0.065618** | **9.856692** |
| **2616** | **0.355791** | **2.303506** | **2.329101** | **4.924116** | **9.912514** |
| **2754** | **1.153772** | **2.507235** | **0.755658** | **5.653636** | **10.0703** |
| **2728** | **0.523353** | **0.774776** | **7.753078** | **1.030164** | **10.08137** |
| **2639** | **5.777024** | **1.015276** | **2.265184** | **1.045051** | **10.10254** |
| **2718** | **0.346026** | **6.726148** | **0** | **3.040884** | **10.11306** |
| **2322** | **1.251465** | **0.846545** | **0.818972** | **7.266298** | **10.18328** |
| **2246** | **1.789292** | **6.933244** | **0** | **1.550209** | **10.27275** |
| **2782** | **0.835896** | **2.072845** | **6.983272** | **0.416894** | **10.30891** |
| **2805** | **5.378** | **1.04917** | **2.722836** | **1.245206** | **10.39521** |
| **2243** | **2.416393** | **2.175277** | **2.5773** | **3.479902** | **10.64887** |
| **2536** | **0.095884** | **3.561788** | **5.82059** | **1.339093** | **10.81736** |
| **2238** | **0.369387** | **0.742187** | **1.510427** | **8.211061** | **10.83306** |
| **2750** | **1.707919** | **4.251871** | **4.76343** | **0.150689** | **10.87391** |
| **2582** | **0.015488** | **7.063076** | **2.857923** | **1.001419** | **10.93791** |
| **2763** | **0.837641** | **1.910903** | **2.306422** | **6.438629** | **11.4936** |
| **2712** | **1.220704** | **3.76591** | **0.904302** | **5.723938** | **11.61485** |
| **2776** | **0.829593** | **6.22405** | **2.933064** | **1.661297** | **11.648** |
| **2745** | **0.483571** | **7.320622** | **3.363013** | **0.494163** | **11.66137** |
| **2292** | **6.336444** | **0.755493** | **0.917318** | **3.729713** | **11.73897** |
| **2313** | **1.667143** | **5.449233** | **1.05964** | **3.572653** | **11.74867** |
| **2797** | **1.177325** | **2.867035** | **5.626742** | **2.078636** | **11.74974** |
| **2517** | **1.221803** | **1.909419** | **6.763958** | **1.994974** | **11.89015** |
| **2772** | **2.646193** | **2.331562** | **4.805164** | **2.226701** | **12.00962** |
| **2736** | **0.484217** | **1.103899** | **4.32752** | **6.178309** | **12.09394** |
| **2764** | **0.631487** | **1.695593** | **2.203302** | **7.563906** | **12.09429** |
| **2793** | **1.489648** | **1.161479** | **1.848343** | **7.710947** | **12.21042** |
| **2795** | **4.984398** | **2.095374** | **3.134766** | **2.759567** | **12.9741** |
| **2325** | **1.238588** | **4.883085** | **4.696405** | **2.29939** | **13.11747** |
| **2729** | **1.588777** | **0.723542** | **10.01355** | **0.941833** | **13.2677** |
| **2309** | **3.67601** | **7.875024** | **1.130204** | **0.646231** | **13.32747** |
| **2779** | **4.80094** | **1.564679** | **2.307191** | **4.689823** | **13.36263** |
| **2769** | **7.138941** | **0.780596** | **3.375685** | **2.178357** | **13.47358** |
| **2780** | **0** | **0** | **0.080836** | **13.41036** | **13.4912** |
| **2293** | **2.924648** | **8.642375** | **1.481113** | **0.740947** | **13.78908** |
| **2251** | **3.64777** | **6.164063** | **3.173318** | **0.833415** | **13.81857** |
| **2741** | **0.400138** | **4.647692** | **7.94182** | **0.992412** | **13.98206** |
| **2617** | **4.338716** | **0.590508** | **7.669857** | **1.826844** | **14.42593** |
| **2785** | **0.430656** | **3.851002** | **6.809705** | **3.641328** | **14.73269** |
| **2669** | **2.33311** | **0.755658** | **3.993671** | **7.883608** | **14.96605** |
| **2609** | **14.18666** | **0.19146** | **0.534177** | **0.334837** | **15.24714** |
| **2264** | **5.988512** | **1.948305** | **1.068748** | **6.533843** | **15.53941** |
| **2324** | **4.450963** | **0.542893** | **10.80094** | **0.051012** | **15.84581** |
| **2291** | **11.32019** | **1.110001** | **1.255965** | **2.227192** | **15.91335** |
| **2762** | **2.502233** | **8.734668** | **0.901248** | **3.885327** | **16.02348** |
| **2751** | **4.399732** | **8.68468** | **0.298078** | **3.663625** | **17.04612** |
| **2307** | **7.975978** | **6.311459** | **2.093122** | **0.954098** | **17.33466** |
| **2281** | **8.308053** | **1.125215** | **6.634586** | **1.30942** | **17.37727** |
| **2276** | **2.622151** | **2.280251** | **5.388041** | **7.502551** | **17.79299** |
| **2245** | **0.30772** | **15.33213** | **1.894182** | **1.003903** | **18.53793** |
| **2306** | **3.757092** | **6.979365** | **5.416031** | **2.409613** | **18.5621** |
| **2787** | **3.793091** | **8.192233** | **2.602365** | **4.318974** | **18.90666** |
| **2778** | **7.244116** | **2.638545** | **5.773732** | **4.604549** | **20.26094** |

**Table S2: Kinase Inhibitor dsRNA Area Under the Curve (AUC)**

|  |  |  |  |  | **dsRNA signal, AU** |  |  |
| --- | --- | --- | --- | --- | --- | --- | --- |
|  | **Cat#** | **Item Name** | **Target** | **Family Name** | **High** | **Low** | **Total** |
| **1** | **A8310** | **PF-562271** | **Pyk2** | **TK** | **0.0** | **690.0** | **690.0** |
| **2** | **N1338** | **Sanguinarine** | **Metabolic Disease** | **Other** | **0.0** | **895.9** | **895.9** |
| **3** | **A8619** | **TIC10** | **Akt** | **AGC** | **0.0** | **1481.6** | **1481.6** |
| **4** | **C6184** | **3-Methylindole** | **NF-?B** | **Other** | **0.0** | **1602.7** | **1602.7** |
| **5** | **B1135** | **GDC-0623** | **MEK1/2** | **STE** | **0.0** | **1908.1** | **1908.1** |
| **6** | **B1539** | **Tideglusib** | **GSK-3** | **CMGC** | **352.7** | **1603.7** | **1956.4** |
| **7** | **B1130** | **GLPG0634** | **JAK** | **TK** | **36.1** | **2690.2** | **2726.3** |
| **8** | **N1715** | **Diosgenin** | **JAK** | **TK** | **0.0** | **2764.9** | **2764.9** |
| **9** | **A3628** | **MLN120B** | **I?B/IKK** | **Other** | **3021.2** | **0.0** | **3021.2** |
| **10** | **A3417** | **Flavopiridol** | **Cyclin-Dependent Kinases** | **CMGC** | **411.3** | **2633.8** | **3045.1** |
| **11** | **A3479** | **Hydroxyfasudil hydrochloride** | **ROCK** | **AGC** | **2331.1** | **751.7** | **3082.8** |
| **12** | **A3524** | **kb NB 142-70** | **PKD** | **CAMK** | **958.0** | **2847.9** | **3805.9** |
| **13** | **A4148** | **NVP-BSK805 2HCl** | **JAK** | **TK** | **1810.2** | **2038.5** | **3848.7** |
| **14** | **A4115** | **JNJ-7706621** | **Aurora Kinase** | **Other** | **0.0** | **4378.3** | **4378.3** |
| **15** | **A3535** | **KX2-391 dihydrochloride** | **Src** | **TK** | **1685.1** | **2878.5** | **4563.6** |
| **16** | **A4150** | **WHI-P154** | **JAK** | **TK** | **5163.1** | **0.0** | **5163.1** |
| **17** | **A8628** | **Chloroquine diphosphate** | **Autophagy** | **Other** | **4180.7** | **1072.6** | **5253.3** |
| **18** | **N2206** | **Mollugin** | **I?B/IKK** | **Other** | **394.8** | **5157.1** | **5551.9** |
| **19** | **A3248** | **BMS345541 hydrochloride** | **I?B/IKK** | **Other** | **0.0** | **5872.9** | **5872.9** |
| **20** | **A5566** | **LY2228820** | **p38** | **CMGC** | **88.7** | **5927.3** | **6016.0** |
| **21** | **A8616** | **A-674563** | **Akt** | **AGC** | **0.0** | **6156.4** | **6156.4** |
| **22** | **A5653** | **AT7867** | **Akt** | **AGC** | **150.0** | **6554.9** | **6704.9** |
| **23** | **B1544** | **Tyrphostin AG 879** | **HER2** | **TK** | **54.0** | **6760.3** | **6814.3** |
| **24** | **N2060** | **12-O-tetradecanoyl phorbol-13-acetate (PMA)** | **Others** | **Other** | **6922.4** | **0.0** | **6922.4** |
| **25** | **A8319** | **Dacomitinib (PF299804, PF299)** | **EGFR** | **TK** | **0.0** | **7030.9** | **7030.9** |
| **26** | **N1677** | **Sennoside B** | **STAT** | **TK** | **3953.3** | **3310.7** | **7264.0** |
| **27** | **B5946** | **EW-7197** | **TGF-? Receptor** | **TKL** | **168.7** | **7205.0** | **7373.6** |
| **28** | **B2171** | **Imatinib (STI571)** | **c-Kit** | **TK** | **4336.5** | **3075.6** | **7412.1** |
| **29** | **B4969** | **KN-93 Phosphate** | **CaM kinase II** | **CAMK** | **4943.4** | **2840.4** | **7783.8** |
| **30** | **C4009** | **CP21R7** | **GSK-3** | **CMGC** | **5189.9** | **2975.5** | **8165.4** |
| **31** | **B3252** | **Dorsomorphin (Compound C)** | **AMPK** | **CAMK** | **6395.6** | **1963.8** | **8359.4** |
| **32** | **A8237** | **SKLB610** | **VEGFR** | **TK** | **126.4** | **8430.0** | **8556.4** |
| **33** | **A8301** | **GW788388** | **TGF-?R1(ALK5)** | **TKL** | **8820.5** | **0.0** | **8820.5** |
| **34** | **A3771** | **RKI-1447** | **ROCK** | **AGC** | **1722.0** | **7250.9** | **8972.9** |
| **35** | **A1805** | **Imatinib Mesylate (STI571)** | **Bcr-Abl** | **TK** | **2759.7** | **6865.2** | **9624.9** |
| **36** | **A8253** | **PD 173074** | **FGFR** | **TK** | **6568.6** | **3359.6** | **9928.2** |
| **37** | **A3337** | **CX-6258** | **Pim** | **CAMK** | **0.0** | **9957.5** | **9957.5** |
| **38** | **N2083** | **Ligustroflavone** | **I?B/IKK** | **Other** | **4049.2** | **6040.1** | **10089.3** |
| **39** | **A8618** | **Palomid 529** | **Akt** | **AGC** | **5778.6** | **4358.2** | **10136.8** |
| **40** | **A8412** | **Dinaciclib (SCH727965)** | **Cyclin-Dependent Kinases** | **CMGC** | **10476.7** | **0.0** | **10476.7** |
| **41** | **A8617** | **CCT128930** | **Akt** | **AGC** | **1507.6** | **9152.4** | **10660.0** |
| **42** | **A2080** | **PIK-75** | **PI3K** | **TK** | **6934.4** | **3754.5** | **10688.9** |
| **43** | **A8207** | **AZD6244 (Selumetinib)** | **MEK1/2** | **STE** | **7299.3** | **3732.5** | **11031.8** |
| **44** | **N2179** | **Cyasterone** | **DYRK** | **CMGC** | **81.6** | **11248.2** | **11329.8** |
| **45** | **A4138** | **Tofacitinib (CP-690550,Tasocitinib)** | **JAK** | **TK** | **6571.2** | **4801.5** | **11372.7** |
| **46** | **B2286** | **K02288** | **TGF-?R1(ALK5)** | **TKL** | **4456.5** | **7061.1** | **11517.7** |
| **47** | **N1878** | **Fumalic acid** | **Others** | **Other** | **4691.7** | **6874.3** | **11566.0** |
| **48** | **A5096** | **PF-04217903** | **c-MET** | **TK** | **9610.7** | **2049.9** | **11660.6** |
| **49** | **A8565** | **Purvalanol B** | **Cyclin-Dependent Kinases** | **CMGC** | **4137.6** | **7805.6** | **11943.2** |
| **50** | **A8394** | **CHIR-124** | **Chk** | **CAMK** | **9794.1** | **2244.6** | **12038.6** |
| **51** | **A4193** | **SMI-4a** | **Pim** | **CAMK** | **0.0** | **12244.4** | **12244.4** |
| **52** | **B5817** | **GDC-0994** | **MEK1/2** | **STE** | **4912.8** | **7646.0** | **12558.8** |
| **53** | **A8889** | **G-749** | **FLT3** | **TK** | **0.0** | **12828.3** | **12828.3** |
| **54** | **A8322** | **Neratinib (HKI-272)** | **EGFR** | **TK** | **1101.0** | **11881.7** | **12982.6** |
| **55** | **A3527** | **Ki20227** | **c-FMS** | **TK** | **0.0** | **13064.0** | **13064.0** |
| **56** | **A4139** | **AG-490** | **EGFR** | **TK** | **8748.3** | **4415.8** | **13164.1** |
| **57** | **A3194** | **AST 487** | **FLT3** | **TK** | **7505.2** | **5709.8** | **13214.9** |
| **58** | **B4786** | **Zotarolimus(ABT-578)** | **mTOR** | **Atypical** | **11435.6** | **1864.8** | **13300.4** |
| **59** | **A1986** | **Nu 6027** | **Cyclin-Dependent Kinases** | **CMGC** | **2596.9** | **10731.2** | **13328.1** |
| **60** | **A8249** | **SB 431542** | **TGF-?R1(ALK5)** | **TKL** | **540.4** | **12958.4** | **13498.8** |
| **61** | **A8548** | **Fingolimod (FTY720)** | **S1P receptor** | **Other** | **0.0** | **13731.4** | **13731.4** |
| **62** | **B7947** | **Aurora Kinase Inhibitor III** | **Aurora Kinase** | **Other** | **2378.0** | **11511.4** | **13889.4** |
| **63** | **B4764** | **TCS-PIM-1-4a** | **Pim** | **CAMK** | **1917.5** | **12250.9** | **14168.4** |
| **64** | **A8247** | **Afatinib (BIBW2992)** | **EGFR** | **TK** | **1924.8** | **12314.4** | **14239.2** |
| **65** | **B6054** | **EAI045** | **EGFR** | **TK** | **0.0** | **14269.6** | **14269.6** |
| **66** | **A3530** | **KN-92 hydrochloride** | **P2X purinergic receptor** | **Other** | **0.0** | **14359.5** | **14359.5** |
| **67** | **A4114** | **MLN8054** | **Aurora Kinase** | **Other** | **4632.4** | **9929.7** | **14562.2** |
| **68** | **B4923** | **TA 01** | **HSC** | **Other** | **672.4** | **13985.5** | **14657.9** |
| **69** | **A3576** | **LY2874455** | **FGFR** | **TK** | **11432.5** | **3320.6** | **14753.1** |
| **70** | **A5573** | **Pimasertib (AS-703026)** | **MEK1/2** | **STE** | **11387.3** | **3667.2** | **15054.4** |
| **71** | **A8464** | **LY2109761** | **TGF-?R1(ALK5)** | **TKL** | **9694.6** | **5823.5** | **15518.1** |
| **72** | **B5970** | **Sanguinarine chloride** | **Protein Ser/Thr Phosphatases** | **CMGC** | **0.0** | **15548.2** | **15548.2** |
| **73** | **A4146** | **CEP-33779** | **JAK** | **TK** | **12244.2** | **3458.4** | **15702.6** |
| **74** | **A8223** | **CID 2011756** | **Protein Ser/Thr Phosphatases** | **CMGC** | **3490.5** | **12424.4** | **15914.8** |
| **75** | **A2168** | **Dovitinib (TKI-258, CHIR-258)** | **FGFR** | **TK** | **15765.0** | **183.2** | **15948.2** |
| **76** | **A8312** | **Torin 1** | **mTOR** | **Atypical** | **1083.7** | **14896.3** | **15980.1** |
| **77** | **B1235** | **NVP 231** | **CERK** | **Other** | **1973.6** | **14037.2** | **16010.8** |
| **78** | **B6025** | **DASA-58** | **PKM2** | **Other** | **7492.3** | **8577.3** | **16069.6** |
| **79** | **B2287** | **LY364947** | **SMAD** | **CMGC** | **12638.5** | **3456.2** | **16094.7** |
| **80** | **B7174** | **DCA** | **Others** | **Other** | **2582.6** | **13605.1** | **16187.7** |
| **81** | **A8640** | **Flavopiridol hydrochloride** | **Cyclin-Dependent Kinases** | **CMGC** | **187.1** | **16139.6** | **16326.6** |
| **82** | **A3342** | **D4476** | **CK1** | **CK1** | **6178.6** | **10405.5** | **16584.1** |
| **83** | **A8678** | **CID 755673** | **PKD** | **CAMK** | **9033.3** | **7582.1** | **16615.4** |
| **84** | **B1537** | **AZD2858** | **GSK-3** | **CMGC** | **10014.5** | **6684.6** | **16699.1** |
| **85** | **A8373** | **AZD2014** | **mTOR** | **Atypical** | **4130.5** | **12584.4** | **16714.9** |
| **86** | **B3288** | **APY29** | **IRE1** | **Other** | **1195.6** | **15781.0** | **16976.7** |
| **87** | **A1882** | **Cediranib (AZD217)** | **VEGFR** | **TK** | **1359.3** | **15765.2** | **17124.5** |
| **88** | **B6007** | **AZD6738** | **ATM/ATR** | **Atypical** | **290.6** | **16844.7** | **17135.3** |
| **89** | **A3660** | **Nilotinib monohydrochloride monohydrate** | **Bcr-Abl** | **TK** | **2759.7** | **14574.2** | **17333.9** |
| **90** | **A8689** | **PD 169316** | **p38** | **CMGC** | **4123.0** | **13218.6** | **17341.6** |
| **91** | **B1494** | **Tyrphostin 9** | **VEGFR** | **TK** | **58.0** | **17328.1** | **17386.1** |
| **92** | **B2176** | **GSK1059615** | **PI3K** | **TK** | **832.8** | **16639.6** | **17472.4** |
| **93** | **N1828** | **Apigenin** | **Immunology & Inflammation related** | **Other** | **1738.8** | **15810.5** | **17549.4** |
| **94** | **B3209** | **Triapine** | **DNA Synthesis** | **Other** | **0.0** | **17637.1** | **17637.1** |
| **95** | **A2754** | **TG100-115** | **PI3K** | **TK** | **401.6** | **17405.4** | **17807.0** |
| **96** | **A3931** | **VX-11e** | **ERK** | **CMGC** | **6059.4** | **11825.4** | **17884.8** |
| **97** | **A3197** | **AT7519 Hydrochloride** | **Cyclin-Dependent Kinases** | **CMGC** | **0.0** | **18031.2** | **18031.2** |
| **98** | **A4136** | **TG101348 (SAR302503)** | **JAK** | **TK** | **3065.1** | **15118.2** | **18183.4** |
| **99** | **B3275** | **TCS JNK 5a** | **JNK** | **CMGC** | **1144.9** | **17159.4** | **18304.3** |
| **100** | **B1124** | **Pitavastatin** | **HMG-CoA Reductase** | **Other** | **8150.8** | **10215.5** | **18366.3** |
| **101** | **A4521** | **TCS PIM-1 1** | **Pim** | **CAMK** | **1935.2** | **16463.0** | **18398.2** |
| **102** | **A8350** | **AZD4547** | **FGFR** | **TK** | **14827.8** | **3634.0** | **18461.7** |
| **103** | **A1404** | **Rigosertib (ON-01910,Estybon)** | **PLK** | **Other** | **14175.0** | **4849.9** | **19024.8** |
| **104** | **B7808** | **NT157** | **IGF1R** | **TK** | **7372.8** | **11852.6** | **19225.3** |
| **105** | **B2173** | **CP-673451** | **VEGFR** | **TK** | **14720.6** | **4566.7** | **19287.4** |
| **106** | **N1908** | **Ginkgolide C** | **P450 (e.g. CYP17)** | **TK** | **4330.9** | **14963.6** | **19294.5** |
| **107** | **B1969** | **Mesalamine** | **PGE synthase** | **AGC** | **17187.6** | **2129.6** | **19317.3** |
| **108** | **A3861** | **TBB** | **CK2** | **CMGC** | **3323.3** | **16520.0** | **19843.3** |
| **109** | **A3719** | **PF-670462** | **CK1** | **CK1** | **1350.7** | **18716.0** | **20066.6** |
| **110** | **A8638** | **LY2603618** | **Chk** | **CAMK** | **13995.2** | **6197.3** | **20192.5** |
| **111** | **A4182** | **Resveratrol** | **Sirtuin** | **Other** | **13503.5** | **6902.6** | **20406.1** |
| **112** | **A3389** | **EMD638683** | **SGK** | **AGC** | **288.1** | **20264.2** | **20552.3** |
| **113** | **B1372** | **Dorsomorphin 2HCl** | **AMPK** | **CAMK** | **1809.4** | **18817.3** | **20626.7** |
| **114** | **A5707** | **PD318088** | **MEK1/2** | **STE** | **9157.7** | **11553.5** | **20711.2** |
| **115** | **A3532** | **KN-93** | **CaM kinase II** | **CAMK** | **14156.8** | **6603.8** | **20760.7** |
| **116** | **A2278** | **NVP-AEW541** | **IGF1R** | **TK** | **1274.0** | **19640.6** | **20914.5** |
| **117** | **A8407** | **Dibucaine (Cinchocaine) HCl** | **Sodium Channel** | **Other** | **9319.1** | **11778.5** | **21097.5** |
| **118** | **B5815** | **LY2584702** | **S6 Kinase** | **AGC** | **5638.3** | **15579.4** | **21217.6** |
| **119** | **B1970** | **Metformin HCl** | **Others** | **Other** | **10385.6** | **10840.1** | **21225.7** |
| **120** | **A3001** | **PCI-32765 (Ibrutinib)** | **BTK** | **TK** | **3646.9** | **17695.2** | **21342.1** |
| **121** | **A3751** | **Regorafenib monohydrate** | **c-RET** | **TK** | **7085.4** | **14285.8** | **21371.2** |
| **122** | **A3531** | **KN-92 phosphate** | **P2X purinergic receptor** | **Other** | **4429.3** | **16946.8** | **21376.0** |
| **123** | **A5979** | **Tie2 kinase inhibitor** | **Tie-2** | **TK** | **4787.9** | **16594.7** | **21382.6** |
| **124** | **A8476** | **MK-2461** | **PDGFR** | **TK** | **19211.6** | **2226.3** | **21437.9** |
| **125** | **A5639** | **BIRB 796 (Doramapimod)** | **p38** | **CMGC** | **8315.3** | **13140.6** | **21455.9** |
| **126** | **A8226** | **TAK-632** | **Raf** | **TKL** | **12667.4** | **8796.8** | **21464.2** |
| **127** | **A2477** | **Tyrphostin AG 1296** | **PDGFR** | **TK** | **2647.7** | **18881.3** | **21528.9** |
| **128** | **A5793** | **Quizartinib (AC220)** | **FLT3** | **TK** | **4499.9** | **17036.5** | **21536.5** |
| **129** | **A3940** | **XL388** | **mTOR** | **Atypical** | **5203.5** | **16443.4** | **21646.9** |
| **130** | **A3148** | **AIM-100** | **Ack1** | **TK** | **17272.0** | **4414.3** | **21686.3** |
| **131** | **C6104** | **L-Leucine** | **Anti-infection** | **Other** | **3716.4** | **18120.1** | **21836.5** |
| **132** | **A3184** | **AR-A014418** | **GSK-3** | **CMGC** | **79.4** | **22152.5** | **22232.0** |
| **133** | **A3397** | **Erlotinib** | **EGFR** | **TK** | **7272.0** | **15317.7** | **22589.7** |
| **134** | **B6674** | **Anisomycin** | **JNK** | **CMGC** | **20082.7** | **2629.3** | **22712.1** |
| **135** | **A3750** | **Regorafenib hydrochloride** | **c-RET** | **TK** | **724.9** | **22021.0** | **22745.9** |
| **136** | **B5663** | **SC 79** | **Akt** | **AGC** | **17618.8** | **5138.7** | **22757.5** |
| **137** | **A3125** | **5-Iodotubercidin** | **Adenosine Kinase** | **Other** | **2639.0** | **20167.4** | **22806.4** |
| **138** | **A4213** | **Necrostatin-1** | **TNF-?** | **CMGC** | **12590.5** | **10276.4** | **22866.9** |
| **139** | **A4510** | **1,2,3,4,5,6-Hexabromocyclohexane** | **JAK** | **TK** | **58.9** | **23091.7** | **23150.6** |
| **140** | **A3321** | **Cobimetinib** | **MEK1/2** | **STE** | **1412.8** | **21814.8** | **23227.7** |
| **141** | **A8304** | **FR 180204** | **ERK** | **CMGC** | **10806.0** | **12597.7** | **23403.7** |
| **142** | **A8347** | **Pazopanib Hydrochloride** | **VEGFR** | **TK** | **14157.8** | **9479.9** | **23637.7** |
| **143** | **A3570** | **LY2090314** | **GSK-3** | **CMGC** | **16169.3** | **7523.7** | **23693.1** |
| **144** | **B1587** | **IMD 0354** | **I?B/IKK** | **Other** | **3341.9** | **20862.1** | **24204.0** |
| **145** | **A4151** | **ZM 39923 HCl** | **JAK** | **TK** | **21395.1** | **2858.2** | **24253.3** |
| **146** | **B1401** | **Cabozantinib malate (XL184)** | **c-MET** | **TK** | **148.7** | **24276.1** | **24424.8** |
| **147** | **B5942** | **AS601245** | **Others** | **Other** | **18398.2** | **6237.2** | **24635.4** |
| **148** | **A2198** | **Genistein** | **Topoisomerase** | **Other** | **5526.1** | **19461.3** | **24987.4** |
| **149** | **A8883** | **SAR405** | **Autophagy** | **Other** | **1587.9** | **23454.9** | **25042.7** |
| **150** | **N2131** | **Wedelolactone** | **EGFR** | **TK** | **9001.4** | **16105.5** | **25106.9** |
| **151** | **B2276** | **Vinpocetine** | **PDE** | **Other** | **5212.2** | **19993.1** | **25205.3** |
| **152** | **A4333** | **CPI-613** | **Dehydrogenase** | **Other** | **36.8** | **25399.4** | **25436.2** |
| **153** | **B5712** | **ANA 12** | **Trk** | **TK** | **5083.5** | **20498.0** | **25581.6** |
| **154** | **A1952** | **NSC 23766** | **Rho** | **AGC** | **10192.1** | **15408.3** | **25600.4** |
| **155** | **A1723** | **Roscovitine (Seliciclib,CYC202)** | **Cyclin-Dependent Kinases** | **CMGC** | **8144.1** | **17609.2** | **25753.3** |
| **156** | **A8890** | **HTH-01-015** | **AMPK** | **CAMK** | **4731.3** | **21129.9** | **25861.2** |
| **157** | **A8303** | **GNF-5837** | **Trk** | **TK** | **2750.0** | **23319.5** | **26069.5** |
| **158** | **A5331** | **TSU-68 (SU6668,Orantinib)** | **VEGFR** | **TK** | **4841.4** | **21696.8** | **26538.2** |
| **159** | **B1642** | **WYE-125132 (WYE-132)** | **mTOR** | **Atypical** | **4894.7** | **21738.1** | **26632.8** |
| **160** | **A3504** | **IRAK inhibitor 6** | **IRAK** | **TKL** | **19776.5** | **7334.0** | **27110.5** |
| **161** | **A1450** | **Lidocaine** | **Histamine Receptor** | **Other** | **26152.6** | **999.3** | **27151.9** |
| **162** | **A3347** | **Dabrafenib Mesylate (GSK-2118436)** | **Raf** | **TKL** | **27414.0** | **0.0** | **27414.0** |
| **163** | **B8365** | **LY3214996** | **ERK** | **CMGC** | **4594.5** | **22927.9** | **27522.4** |
| **164** | **A2949** | **Linifanib (ABT-869)** | **VEGFR** | **TK** | **6252.9** | **21480.2** | **27733.1** |
| **165** | **A8885** | **Ro 3306** | **Cyclin-Dependent Kinases** | **CMGC** | **2807.1** | **25048.4** | **27855.5** |
| **166** | **A3794** | **SB1317** | **JAK** | **TK** | **1210.9** | **26680.2** | **27891.1** |
| **167** | **A3433** | **Gefitinib hydrochloride** | **EGFR** | **TK** | **0.0** | **28000.5** | **28000.5** |
| **168** | **A2306** | **Ellagic acid** | **Topoisomerase** | **Other** | **14274.7** | **14007.9** | **28282.5** |
| **169** | **A8636** | **RN486** | **BTK** | **TK** | **3432.0** | **25024.9** | **28456.9** |
| **170** | **B6116** | **GLPG0634-A** | **JAK** | **TK** | **20519.6** | **8028.8** | **28548.5** |
| **171** | **A3006** | **GDC-0068 (RG7440)** | **Akt** | **AGC** | **10110.1** | **18551.4** | **28661.5** |
| **172** | **B4877** | **URMC-099** | **Others** | **Other** | **20784.2** | **7905.7** | **28690.0** |
| **173** | **B3697** | **4E1RCat** | **Others** | **Other** | **11534.6** | **17206.1** | **28740.7** |
| **174** | **A8248** | **ZSTK474** | **PI3K** | **TK** | **6069.9** | **23073.5** | **29143.5** |
| **175** | **A5506** | **Thiazovivin** | **ROCK** | **AGC** | **2364.1** | **26827.3** | **29191.4** |
| **176** | **B2186** | **GSK2636771** | **PI3K** | **TK** | **841.3** | **28354.9** | **29196.1** |
| **177** | **C4445** | **2-D08** | **SUMOylation** | **Other** | **21678.5** | **7689.9** | **29368.4** |
| **178** | **A3206** | **AVL-292** | **BTK** | **TK** | **20034.7** | **9433.9** | **29468.6** |
| **179** | **B1621** | **TAK-733** | **MEK1/2** | **STE** | **15107.2** | **14755.9** | **29863.1** |
| **180** | **B1585** | **SC-514** | **I?B/IKK** | **Other** | **620.3** | **29713.1** | **30333.4** |
| **181** | **A8199** | **PD153035 hydrochloride** | **EGFR** | **TK** | **1669.4** | **28866.3** | **30535.6** |
| **182** | **A3943** | **XMD8-92** | **ERK** | **CMGC** | **359.5** | **30524.1** | **30883.6** |
| **183** | **A2412** | **CP-724714** | **EGFR** | **TK** | **1118.3** | **29983.8** | **31102.1** |
| **184** | **B1409** | **Benidipine HCl** | **Calcium Channel** | **Other** | **130.4** | **31078.1** | **31208.6** |
| **185** | **N1802** | **Oridonin** | **Anti-infection** | **Other** | **6061.2** | **25375.1** | **31436.3** |
| **186** | **A1655** | **GW2580** | **CSF-1R** | **TK** | **17974.9** | **13531.9** | **31506.8** |
| **187** | **A4192** | **SGI-1776 free base** | **Pim** | **CAMK** | **1583.1** | **29976.5** | **31559.5** |
| **188** | **A2552** | **Quercetin dihydrate** | **PI3K** | **TK** | **25411.6** | **6435.1** | **31846.8** |
| **189** | **A8551** | **INK 128 (MLN0128)** | **mTOR** | **Atypical** | **0.0** | **31875.7** | **31875.7** |
| **190** | **B3702** | **MSDC-0160** | **Others** | **Other** | **16528.8** | **15455.5** | **31984.3** |
| **191** | **A3392** | **Emodin** | **NF-?B** | **Other** | **20341.5** | **11688.8** | **32030.2** |
| **192** | **B2182** | **CAY10505** | **PI3K** | **TK** | **15375.8** | **16784.8** | **32160.6** |
| **193** | **A3792** | **SB 239063** | **p38** | **CMGC** | **16458.2** | **15855.6** | **32313.9** |
| **194** | **B4492** | **HS-173** | **PI3K** | **TK** | **345.7** | **31987.4** | **32333.1** |
| **195** | **B1525** | **SSR128129E** | **FGFR** | **TK** | **4736.0** | **27624.5** | **32360.5** |
| **196** | **A3306** | **Chelerythrine Chloride** | **PKC** | **AGC** | **10306.1** | **22120.5** | **32426.7** |
| **197** | **B5937** | **KPT-9274** | **PAK4** | **STE** | **6803.0** | **25813.6** | **32616.6** |
| **198** | **N1849** | **Dihydromyricetin** | **MEK** | **STE** | **20863.8** | **12380.4** | **33244.2** |
| **199** | **B1011** | **Bafetinib (INNO-406)** | **Bcr-Abl** | **TK** | **1113.6** | **32473.2** | **33586.8** |
| **200** | **B4842** | **LDN-214117** | **ALK** | **TK** | **4693.4** | **28899.6** | **33593.0** |
| **201** | **B1249** | **TDZD-8** | **GSK-3** | **CMGC** | **7580.3** | **26092.4** | **33672.7** |
| **202** | **B8316** | **Harmine** | **PPAR,MAO** | **Other** | **9958.7** | **24294.0** | **34252.7** |
| **203** | **A8902** | **6H05** | **Rho** | **AGC** | **9210.5** | **25435.6** | **34646.0** |
| **204** | **B1405** | **GW5074** | **Raf** | **TKL** | **14683.4** | **20240.6** | **34924.0** |
| **205** | **A3222** | **Baricitinib phosphate** | **JAK** | **TK** | **2518.9** | **32442.1** | **34961.0** |
| **206** | **B3570** | **CGP 57380** | **Others** | **Other** | **1947.6** | **33084.2** | **35031.8** |
| **207** | **A4152** | **BMS-911543** | **JAK** | **TK** | **18362.9** | **16684.7** | **35047.6** |
| **208** | **A1894** | **SL-327** | **MEK1/2** | **STE** | **10911.9** | **24198.7** | **35110.7** |
| **209** | **A3887** | **Trametinib DMSO solvate** | **MEK1/2** | **STE** | **10918.5** | **24279.8** | **35198.3** |
| **210** | **A3448** | **GSK2606414** | **PERK** | **Other** | **6871.8** | **28375.6** | **35247.4** |
| **211** | **A2846** | **OSU-03012 (AR-12)** | **PDK-1** | **AGC** | **17565.3** | **17813.3** | **35378.6** |
| **212** | **B2175** | **GSK2656157** | **PERK** | **Other** | **2371.9** | **33522.6** | **35894.5** |
| **213** | **A8546** | **R406** | **Spleen Tyrosine Kinase (Syk)** | **TK** | **5357.1** | **30563.4** | **35920.5** |
| **214** | **B1496** | **Icotinib** | **EGFR** | **TK** | **11365.1** | **24561.1** | **35926.2** |
| **215** | **B4809** | **K-115** | **Rho** | **AGC** | **25318.1** | **10922.3** | **36240.4** |
| **216** | **A3923** | **VO-Ohpic trihydrate** | **PTEN** | **TK** | **28838.2** | **7631.3** | **36469.5** |
| **217** | **A8320** | **PF-00562271** | **FAK** | **TK** | **0.0** | **36485.5** | **36485.5** |
| **218** | **N1308** | **Biochanin A** | **Antioxidant** | **Other** | **14982.3** | **21751.5** | **36733.9** |
| **219** | **B2303** | **Apatinib** | **VEGFR** | **TK** | **181.1** | **36614.7** | **36795.8** |
| **220** | **B4986** | **LY2409881** | **I?B/IKK** | **Other** | **22360.9** | **15064.3** | **37425.3** |
| **221** | **A8324** | **LDN-193189** | **SMAD** | **CMGC** | **30043.0** | **7399.9** | **37442.8** |
| **222** | **A4120** | **MK-5108 (VX-689)** | **Aurora Kinase** | **Other** | **3258.6** | **34308.8** | **37567.4** |
| **223** | **B4904** | **ACTB-1003** | **FGFR** | **TK** | **28009.9** | **9584.0** | **37593.9** |
| **224** | **A3626** | **ML-7 hydrochloride** | **ATPase** | **Other** | **19324.6** | **18346.9** | **37671.5** |
| **225** | **N1827** | **Artemisinine** | **P450 (e.g. CYP17)** | **TK** | **2826.9** | **35035.3** | **37862.2** |
| **226** | **A3529** | **KN-92** | **CaM kinase II** | **CAMK** | **8269.0** | **30018.5** | **38287.5** |
| **227** | **B6115** | **RG 13022** | **EGFR** | **TK** | **36154.5** | **2414.3** | **38568.8** |
| **228** | **A2600** | **(-)-Epigallocatechin gallate (EGCG)** | **PKC** | **AGC** | **14230.9** | **24448.0** | **38678.9** |
| **229** | **A1632** | **SB202190 (FHPI)** | **p38** | **CMGC** | **29217.1** | **9508.1** | **38725.2** |
| **230** | **B9000** | **8-Bromo-cAMP, sodium salt** | **cAMP** | **AGC** | **25859.7** | **12957.0** | **38816.6** |
| **231** | **A4237** | **Amuvatinib (MP-470, HPK 56)** | **c-RET** | **TK** | **6649.7** | **32179.3** | **38829.0** |
| **232** | **A4121** | **SNS-314 Mesylate** | **Aurora Kinase** | **Other** | **5716.5** | **33317.4** | **39033.9** |
| **233** | **A3805** | **SCH772984** | **MEK1/2** | **STE** | **21169.8** | **17905.7** | **39075.6** |
| **234** | **A8504** | **Pitavastatin Calcium** | **HMG-CoA Reductase** | **Other** | **16024.1** | **23195.0** | **39219.2** |
| **235** | **A8881** | **WZ3146** | **EGFR** | **TK** | **23148.6** | **16337.5** | **39486.1** |
| **236** | **B5487** | **EHT 1864** | **Amyloid ?** | **Other** | **22471.9** | **17099.7** | **39571.7** |
| **237** | **A8214** | **AZD8055** | **mTOR** | **Atypical** | **0.0** | **39910.8** | **39910.8** |
| **238** | **A3193** | **ASP3026** | **ALK** | **TK** | **11916.3** | **27995.4** | **39911.7** |
| **239** | **A8318** | **PP242** | **mTOR** | **Atypical** | **7321.2** | **32614.7** | **39936.0** |
| **240** | **B5853** | **MHY1485** | **mTOR** | **Atypical** | **4962.4** | **35655.5** | **40617.9** |
| **241** | **B6193** | **AMG 337** | **c-MET** | **TK** | **6788.1** | **33845.4** | **40633.5** |
| **242** | **B5950** | **AZD8186** | **PI3K** | **TK** | **4769.3** | **35908.0** | **40677.3** |
| **243** | **A2067** | **PI-103** | **PI3K** | **TK** | **0.0** | **40731.1** | **40731.1** |
| **244** | **A1792** | **PD184352 (CI-1040)** | **MEK1/2** | **STE** | **0.0** | **41067.4** | **41067.4** |
| **245** | **B5846** | **Radotinib(IY-5511)** | **Bcl-2 Family** | **TK** | **11726.8** | **29413.0** | **41139.9** |
| **246** | **C4953** | **HA-100 (hydrochloride)** | **Broad Spectrum Protein Kinase Inhibitor** | **Other** | **27295.9** | **14084.7** | **41380.5** |
| **247** | **A3676** | **NVP-LCQ195** | **Cyclin-Dependent Kinases** | **CMGC** | **6040.0** | **35350.1** | **41390.1** |
| **248** | **N1743** | **Curcumol** | **Others** | **Other** | **0.0** | **41441.3** | **41441.3** |
| **249** | **B4907** | **Mps1-IN-1** | **Mps1** | **Other** | **7290.6** | **34349.9** | **41640.6** |
| **250** | **A5092** | **JNJ-38877605** | **c-MET** | **TK** | **9345.0** | **32468.4** | **41813.4** |
| **251** | **A3556** | **LKB1 (AAK1 dual inhibitor)** | **Pim** | **CAMK** | **42067.1** | **0.0** | **42067.1** |
| **252** | **A8336** | **KU-60019** | **ATM/ATR** | **Atypical** | **31471.9** | **10671.9** | **42143.8** |
| **253** | **B5832** | **Altiratinib** | **c-MET** | **TK** | **7645.1** | **34585.3** | **42230.4** |
| **254** | **A4110** | **MLN8237 (Alisertib)** | **Aurora Kinase** | **Other** | **9976.5** | **32491.8** | **42468.3** |
| **255** | **B1536** | **AZD1080** | **GSK-3** | **CMGC** | **30210.5** | **12322.5** | **42533.0** |
| **256** | **B4357** | **UNC2881** | **Axl** | **TK** | **446.4** | **42094.2** | **42540.5** |
| **257** | **A3320** | **CO-1686 (AVL-301)** | **EGFR** | **TK** | **16434.5** | **26200.1** | **42634.6** |
| **258** | **B4787** | **SF1670** | **PTEN** | **TK** | **30745.4** | **12301.2** | **43046.5** |
| **259** | **A8326** | **AZD-5438** | **Cyclin-Dependent Kinases** | **CMGC** | **10643.7** | **32871.5** | **43515.3** |
| **260** | **C5579** | **Brilliant Blue G** | **Others** | **Other** | **11949.6** | **31753.4** | **43703.1** |
| **261** | **A3505** | **IRAK-1-4 Inhibitor I** | **IRAK** | **TKL** | **21347.7** | **22363.0** | **43710.7** |
| **262** | **A1173** | **AG-18** | **EGFR** | **TK** | **29672.3** | **14434.9** | **44107.2** |
| **263** | **A8711** | **DDD107498** | **Antimalaria** | **Other** | **38942.1** | **5409.3** | **44351.3** |
| **264** | **B4094** | **NMS-1286937** | **PLK** | **Other** | **5848.8** | **38547.7** | **44396.5** |
| **265** | **A5611** | **GSK429286A** | **ROCK** | **AGC** | **10329.9** | **34075.4** | **44405.3** |
| **266** | **A1196** | **SGX-523** | **c-MET** | **TK** | **18702.9** | **25732.4** | **44435.3** |
| **267** | **B2219** | **EHop-016** | **Rho** | **AGC** | **0.0** | **44568.8** | **44568.8** |
| **268** | **B4660** | **PI-3065** | **PI3K** | **TK** | **625.6** | **44145.1** | **44770.7** |
| **269** | **B3696** | **4EGI-1** | **Others** | **Other** | **27081.3** | **17726.3** | **44807.6** |
| **270** | **B1760** | **Flufenamic acid** | **AMPK; Calcium Channel; Chloride Channel; COX; Potassium Channel** | **CAMK** | **29762.4** | **15182.6** | **44945.1** |
| **271** | **B2032** | **SRPIN340** | **SRPK** | **CMGC** | **38969.9** | **6071.8** | **45041.7** |
| **272** | **A8184** | **AICAR** | **AMPK** | **CAMK** | **10649.1** | **34470.9** | **45119.9** |
| **273** | **A3209** | **AXL1717** | **IGF1R** | **TK** | **13380.1** | **31807.4** | **45187.5** |
| **274** | **A5071** | **GDC-0879** | **Raf** | **TKL** | **7041.8** | **38492.2** | **45534.0** |
| **275** | **A5072** | **GSK690693** | **Akt** | **AGC** | **6007.3** | **39582.1** | **45589.3** |
| **276** | **A8251** | **TAE684 (NVP-TAE684)** | **ALK** | **TK** | **41419.4** | **4182.3** | **45601.7** |
| **277** | **A8679** | **CRT 0066101** | **PKD** | **CAMK** | **0.0** | **45629.7** | **45629.7** |
| **278** | **B8328** | **AZD1390** | **ATM/ATR** | **Atypical** | **7959.8** | **37767.2** | **45727.1** |
| **279** | **A1169** | **10058-F4** | **c-Myc** | **Other** | **21785.0** | **24031.2** | **45816.2** |
| **280** | **A2673** | **AG-1024** | **IGF1R** | **TK** | **1211.9** | **44737.5** | **45949.4** |
| **281** | **N1768** | **Rosmarinic acid** | **Others** | **Other** | **3864.4** | **42457.9** | **46322.3** |
| **282** | **A2307** | **PHA-665752** | **c-MET** | **TK** | **34815.2** | **11571.8** | **46387.0** |
| **283** | **B1236** | **BML-277** | **Chk** | **CAMK** | **33021.5** | **13488.9** | **46510.4** |
| **284** | **C5108** | **Amlexanox** | **Melatonin Receptors** | **Other** | **17956.6** | **28672.3** | **46628.9** |
| **285** | **A4132** | **CCT137690** | **Aurora Kinase** | **Other** | **0.0** | **46657.6** | **46657.6** |
| **286** | **A5057** | **MGCD-265** | **Tie-2** | **TK** | **4249.4** | **42710.5** | **46959.9** |
| **287** | **A3018** | **Trametinib (GSK1120212)** | **MEK1/2** | **STE** | **36456.5** | **10541.3** | **46997.8** |
| **288** | **A3014** | **BGJ398** | **FGFR** | **TK** | **24356.1** | **22673.4** | **47029.5** |
| **289** | **B1262** | **HG-10-102-01** | **LRRK2** | **TKL** | **14367.4** | **32713.5** | **47080.9** |
| **290** | **B7912** | **Pyridoxine** | **Vitamin** | **Other** | **17847.0** | **29400.2** | **47247.2** |
| **291** | **B6017** | **AZD3264** | **I?B/IKK** | **Other** | **223.2** | **47155.4** | **47378.5** |
| **292** | **A8625** | **CP-466722** | **ATM/ATR** | **Atypical** | **23326.1** | **24439.1** | **47765.2** |
| **293** | **A8525** | **Sotrastaurin (AEB071)** | **PKC** | **AGC** | **6965.2** | **40818.2** | **47783.4** |
| **294** | **A8348** | **LY2157299** | **TGF-?R1(ALK5)** | **TKL** | **33079.0** | **15107.1** | **48186.1** |
| **295** | **B1492** | **PYR-41** | **E1 Activating** | **Other** | **41809.3** | **6659.6** | **48468.9** |
| **296** | **A3353** | **DB07268** | **JNK** | **CMGC** | **10652.5** | **37894.5** | **48547.0** |
| **297** | **B6171** | **CC-223** | **mTOR** | **Atypical** | **6587.4** | **41988.9** | **48576.2** |
| **298** | **B5484** | **7,8-Dihydroxyflavone** | **Trk** | **TK** | **9230.6** | **39399.4** | **48630.0** |
| **299** | **A3354** | **DCC-2618** | **PDGFR** | **TK** | **31390.7** | **17341.0** | **48731.7** |
| **300** | **B6114** | **CC-115** | **DNA-PK** | **Other** | **5666.1** | **44138.8** | **49805.0** |
| **301** | **B1293** | **Y-27632** | **ROCK** | **AGC** | **3508.5** | **46665.7** | **50174.2** |
| **302** | **A8620** | **AZD-3463** | **ALK** | **TK** | **9390.1** | **40804.7** | **50194.8** |
| **303** | **A8541** | **Triciribine** | **Akt** | **AGC** | **3840.6** | **46439.7** | **50280.3** |
| **304** | **A3420** | **FMK** | **S6 Kinase** | **AGC** | **36246.0** | **14082.6** | **50328.7** |
| **305** | **B4688** | **3CAI** | **Akt** | **AGC** | **6736.4** | **43810.0** | **50546.4** |
| **306** | **B2189** | **YM201636** | **PIKfyve** | **TK** | **7268.3** | **43298.6** | **50566.9** |
| **307** | **A8300** | **ZCL278** | **Cdc42** | **CMGC** | **9541.3** | **41590.0** | **51131.3** |
| **308** | **B2297** | **GW441756** | **Trk** | **TK** | **33214.4** | **18067.5** | **51281.9** |
| **309** | **A1387** | **AZD5363** | **Akt** | **AGC** | **11890.0** | **39804.7** | **51694.7** |
| **310** | **A2597** | **Brivanib (BMS-540215)** | **VEGFR** | **TK** | **51042.4** | **684.7** | **51727.0** |
| **311** | **B4754** | **LDC000067** | **Cyclin-Dependent Kinases** | **CMGC** | **31035.0** | **20712.1** | **51747.0** |
| **312** | **A4141** | **Baricitinib (LY3009104, INCB028050)** | **JAK** | **TK** | **27614.1** | **24279.2** | **51893.3** |
| **313** | **A2977** | **Cabozantinib (XL184, BMS-907351)** | **c-MET** | **TK** | **13075.8** | **39205.8** | **52281.6** |
| **314** | **B4990** | **Purvalanol A** | **Cyclin-Dependent Kinases** | **CMGC** | **51443.1** | **865.0** | **52308.0** |
| **315** | **A2689** | **Butein** | **EGFR** | **TK** | **38311.3** | **14009.4** | **52320.7** |
| **316** | **A4143** | **CYT387** | **JAK** | **TK** | **36485.3** | **15837.3** | **52322.6** |
| **317** | **B1162** | **FRAX597** | **PAK1** | **STE** | **1538.8** | **50796.9** | **52335.7** |
| **318** | **A3302** | **CGI-1746** | **BTK** | **TK** | **34116.6** | **18368.1** | **52484.7** |
| **319** | **B2226** | **SKI II** | **S1P receptor** | **Other** | **31334.0** | **21341.0** | **52674.9** |
| **320** | **B4924** | **TA 02** | **HSC** | **Other** | **8658.0** | **44038.4** | **52696.4** |
| **321** | **A3013** | **PD0325901** | **MEK1/2** | **STE** | **5022.8** | **48059.9** | **53082.7** |
| **322** | **C5386** | **N-Acetylserotonin** | **MKK** | **CMGC** | **219.3** | **53222.8** | **53442.1** |
| **323** | **B6108** | **IC261** | **CK1** | **CK1** | **6564.9** | **46952.8** | **53517.7** |
| **324** | **A8354** | **A66** | **PI3K** | **TK** | **12407.5** | **41352.4** | **53759.9** |
| **325** | **A8661** | **MNS** | **Integrin** | **Other** | **1958.5** | **51870.8** | **53829.2** |
| **326** | **N2182** | **Harmine hydrochloride** | **STAT3** | **TK** | **0.0** | **53937.1** | **53937.1** |
| **327** | **A8240** | **SB 216763** | **GSK-3** | **CMGC** | **596.7** | **53506.5** | **54103.1** |
| **328** | **C5638** | **13(Z)-Docosenoic Acid** | **Others** | **Other** | **4336.2** | **49926.0** | **54262.2** |
| **329** | **A5112** | **XL147** | **PI3K** | **TK** | **6847.8** | **47539.1** | **54386.9** |
| **330** | **B1639** | **Ridaforolimus (Deforolimus, MK-8669)** | **mTOR** | **Atypical** | **23072.2** | **31512.5** | **54584.8** |
| **331** | **N1753** | **Scutellarin** | **I?B/IKK** | **Other** | **1769.9** | **52840.4** | **54610.3** |
| **332** | **A8250** | **LY 294002** | **PI3K** | **TK** | **11164.3** | **43579.3** | **54743.6** |
| **333** | **A3145** | **Afatinib dimaleate** | **HER2** | **TK** | **15371.8** | **40279.2** | **55651.0** |
| **334** | **B7853** | **Tandutinib (MLN518) HCl** | **FLT3** | **TK** | **42444.4** | **13615.8** | **56060.2** |
| **335** | **A8234** | **Erlotinib Hydrochloride** | **EGFR** | **TK** | **3624.7** | **52457.7** | **56082.4** |
| **336** | **A3432** | **GDC-0941 dimethanesulfonate** | **PI3K** | **TK** | **20950.7** | **35251.8** | **56202.5** |
| **337** | **A3674** | **NVP-BKM120 Hydrochloride** | **PI3K** | **TK** | **0.0** | **56514.5** | **56514.5** |
| **338** | **A8330** | **CX-4945 (Silmitasertib)** | **CK2** | **CMGC** | **35326.5** | **21811.4** | **57137.9** |
| **339** | **A4488** | **Anacardic acid** | **Aurora Kinase** | **Other** | **12779.5** | **44703.4** | **57482.9** |
| **340** | **A3939** | **XL228** | **Aurora Kinase** | **Other** | **17836.6** | **39819.2** | **57655.8** |
| **341** | **A8210** | **GDC-0941** | **PI3K** | **TK** | **590.8** | **57346.3** | **57937.2** |
| **342** | **B3688** | **ML347** | **TGF-?R1(ALK5)** | **TKL** | **0.0** | **58224.3** | **58224.3** |
| **343** | **A5703** | **BMS-777607** | **c-MET** | **TK** | **39539.6** | **18689.6** | **58229.2** |
| **344** | **N2643** | **Shikonin** |  | **Other** | **54262.1** | **4009.3** | **58271.4** |
| **345** | **A8308** | **PH-797804** | **p38** | **CMGC** | **733.9** | **57938.0** | **58671.9** |
| **346** | **B3686** | **DMH-1** | **TGF-?R1(ALK5)** | **TKL** | **13545.0** | **45359.2** | **58904.2** |
| **347** | **A8353** | **3-Methyladenine** | **Autophagy** | **Other** | **32357.3** | **26977.2** | **59334.5** |
| **348** | **N1620** | **Ginsenoside Rb1** | **Others** | **Other** | **7414.4** | **52585.8** | **60000.1** |
| **349** | **A3847** | **SU5416** | **c-RET** | **TK** | **39551.1** | **20455.1** | **60006.1** |
| **350** | **B1439** | **Golvatinib (E7050)** | **c-MET** | **TK** | **36777.1** | **23267.5** | **60044.6** |
| **351** | **N2031** | **Piceatannol** | **PKC; SPHK** | **AGC** | **12025.1** | **48262.2** | **60287.2** |
| **352** | **A8167** | **Rapamycin (Sirolimus)** | **mTOR** | **Atypical** | **1745.1** | **58744.6** | **60489.7** |
| **353** | **A2065** | **IC-87114** | **PI3K** | **TK** | **0.0** | **60540.6** | **60540.6** |
| **354** | **C6276** | **Ethyl gallate** | **Others** | **Other** | **180.5** | **60976.1** | **61156.7** |
| **355** | **A5176** | **AS-605240** | **PI3K** | **TK** | **11785.5** | **49405.3** | **61190.8** |
| **356** | **A1663** | **PD98059** | **MEK1/2** | **STE** | **33353.3** | **27955.5** | **61308.8** |
| **357** | **A3005** | **CAL-101 (Idelalisib, GS-1101)** | **PI3K** | **TK** | **57616.6** | **3708.3** | **61324.9** |
| **358** | **A8489** | **NVP-BVU972** | **c-MET** | **TK** | **9035.4** | **52705.1** | **61740.5** |
| **359** | **A3388** | **EMD-1214063** | **c-MET** | **TK** | **14120.6** | **47689.1** | **61809.6** |
| **360** | **B3661** | **(+)-Usniacin** | **Others** | **Other** | **3049.6** | **59009.6** | **62059.2** |
| **361** | **N1958** | **Galangin;3,5,7-Trihydroxyflavone** | **I?B/IKK** | **Other** | **16189.5** | **46015.9** | **62205.4** |
| **362** | **A3149** | **AKT inhibitor VIII** | **Akt** | **AGC** | **14379.0** | **47838.1** | **62217.1** |
| **363** | **A8232** | **Nilotinib(AMN-107)** | **Bcr-Abl** | **TK** | **28653.4** | **33622.0** | **62275.3** |
| **364** | **A3732** | **Poloxin** | **PLK** | **Other** | **3268.7** | **59035.4** | **62304.1** |
| **365** | **B7716** | **Senexin A** | **CDK** | **CMGC** | **30520.6** | **31796.6** | **62317.2** |
| **366** | **B1431** | **TG003** | **CDK** | **CMGC** | **1126.7** | **61445.9** | **62572.6** |
| **367** | **A5602** | **SB525334** | **TGF-?R1(ALK5)** | **TKL** | **34680.3** | **27941.4** | **62621.6** |
| **368** | **A1821** | **Ki8751** | **VEGFR** | **TK** | **16810.1** | **45822.4** | **62632.5** |
| **369** | **A2822** | **AC480 (BMS-599626)** | **EGFR** | **TK** | **468.1** | **62401.2** | **62869.3** |
| **370** | **A1186** | **AMG-208** | **c-MET** | **TK** | **40405.8** | **22501.3** | **62907.1** |
| **371** | **A8603** | **GNF 2** | **Bcr-Abl** | **TK** | **2578.1** | **60765.8** | **63344.0** |
| **372** | **A8236** | **Regorafenib** | **c-RET** | **TK** | **93.0** | **63322.3** | **63415.2** |
| **373** | **A5803** | **BIX 02188** | **MEK1/2** | **STE** | **40230.6** | **23376.7** | **63607.2** |
| **374** | **B1371** | **Miltefosine** | **Akt** | **AGC** | **27276.6** | **36330.6** | **63607.3** |
| **375** | **A3022** | **Pazopanib (GW-786034)** | **PDGFR** | **TK** | **41143.8** | **22671.3** | **63815.1** |
| **376** | **B2190** | **H 89 2HCl** | **PKA** | **AGC** | **7807.6** | **56279.7** | **64087.4** |
| **377** | **A8418** | **Dovitinib Dilactic acid** | **VEGFR** | **TK** | **47107.5** | **17331.2** | **64438.8** |
| **378** | **A3260** | **BMX-IN-1** | **BMX Kinase** | **TK** | **37737.5** | **26747.2** | **64484.7** |
| **379** | **A8329** | **R428** | **Axl** | **TK** | **1050.4** | **64667.0** | **65717.4** |
| **380** | **A8325** | **Tivantinib (ARQ 197)** | **c-MET** | **TK** | **46631.5** | **19485.6** | **66117.2** |
| **381** | **A8604** | **GNF 5** | **Bcr-Abl** | **TK** | **498.5** | **65630.6** | **66129.1** |
| **382** | **A8550** | **Telatinib (BAY 57-9352)** | **VEGFR** | **TK** | **46565.4** | **19646.6** | **66212.0** |
| **383** | **B5860** | **TAK960** | **PLK** | **Other** | **9929.6** | **56355.2** | **66284.8** |
| **384** | **B9001** | **Dibutyryl-cAMP, sodium salt** | **cAMP** | **AGC** | **10702.1** | **55803.3** | **66505.4** |
| **385** | **A3545** | **LDN193189 Hydrochloride** | **SMAD** | **CMGC** | **42525.8** | **24650.0** | **67175.8** |
| **386** | **B1998** | **Oxfendazole** | **Anti-infection** | **Other** | **20012.9** | **47370.5** | **67383.3** |
| **387** | **B3699** | **ISRIB (trans-isomer)** | **PERK** | **Other** | **6980.3** | **60689.2** | **67669.5** |
| **388** | **B1543** | **Mubritinib (TAK 165)** | **HER2** | **TK** | **19342.6** | **48533.2** | **67875.8** |
| **389** | **B1027** | **2-Deoxy-D-glucose** | **Hexokinase** | **Other** | **5572.7** | **62378.7** | **67951.4** |
| **390** | **B1641** | **WAY-600** | **mTOR** | **Atypical** | **1003.4** | **67106.4** | **68109.8** |
| **391** | **A8357** | **AG-1478** | **EGFR** | **TK** | **1676.2** | **67129.9** | **68806.1** |
| **392** | **B1374** | **WZ4003** | **AMPK** | **CAMK** | **30589.4** | **38446.0** | **69035.5** |
| **393** | **A2323** | **TCS 359** | **FLT3** | **TK** | **5896.0** | **63380.8** | **69276.8** |
| **394** | **A5719** | **AT7519** | **Cyclin-Dependent Kinases** | **CMGC** | **23488.0** | **46607.1** | **70095.0** |
| **395** | **B1437** | **PF-477736** | **Chk** | **CAMK** | **45938.3** | **24364.8** | **70303.1** |
| **396** | **B7850** | **BMS-582949 hydrochloride** | **p38** | **CMGC** | **7441.1** | **64364.9** | **71806.0** |
| **397** | **A8688** | **TAK-715** | **p38** | **CMGC** | **42569.8** | **29513.1** | **72082.8** |
| **398** | **B2114** | **Mitoxantrone HCl** | **Topoisomerase** | **Other** | **48235.0** | **23848.1** | **72083.1** |
| **399** | **N1609** | **Ginsenoside Rh2** | **Others** | **Other** | **41827.1** | **30569.1** | **72396.2** |
| **400** | **A3011** | **CHIR-99021 (CT99021)** | **GSK-3** | **CMGC** | **62333.9** | **10204.7** | **72538.6** |
| **401** | **B7815** | **NSC228155** | **EGFR** | **TK** | **18547.9** | **54998.4** | **73546.3** |
| **402** | **N1789** | **Salidroside** | **Others** | **Other** | **19295.0** | **56220.3** | **75515.3** |
| **403** | **N1615** | **Ginsenoside Re** | **Others** | **Other** | **10054.1** | **65920.2** | **75974.2** |
| **404** | **A2251** | **Tivozanib (AV-951)** | **VEGFR** | **TK** | **40142.9** | **36133.0** | **76275.9** |
| **405** | **A3760** | **Reversine** | **Aurora Kinase** | **Other** | **27234.6** | **49415.3** | **76649.9** |
| **406** | **B5816** | **PLX647** | **c-FMS** | **TK** | **24285.0** | **52941.2** | **77226.3** |
| **407** | **B1540** | **TWS119** | **GSK-3** | **CMGC** | **48959.2** | **29417.1** | **78376.3** |
| **408** | **B2301** | **SAR131675** | **VEGFR** | **TK** | **44218.5** | **34484.3** | **78702.8** |
| **409** | **A8343** | **Go 6983** | **PKC** | **AGC** | **52923.0** | **26190.1** | **79113.1** |
| **410** | **A4512** | **Cucurbitacin I** | **JAK** | **TK** | **39627.9** | **39750.1** | **79378.0** |
| **411** | **A2149** | **Bosutinib (SKI-606)** | **Bcr-Abl** | **TK** | **38090.7** | **42027.4** | **80118.1** |
| **412** | **A3825** | **SLx-2119** | **ROCK** | **AGC** | **20785.5** | **59494.3** | **80279.8** |
| **413** | **B8023** | **Cerdulatinib (PRT062070)** | **JAK** | **TK** | **60131.0** | **21245.1** | **81376.1** |
| **414** | **B3300** | **Cromolyn sodium** | **Calcium Channel** | **Other** | **64024.7** | **17957.1** | **81981.8** |
| **415** | **A2174** | **Lenvatinib (E7080)** | **VEGFR** | **TK** | **26964.0** | **55504.0** | **82468.0** |
| **416** | **B4761** | **GNE-9605** | **LRRK2** | **TKL** | **40803.1** | **41939.9** | **82743.0** |
| **417** | **B5827** | **Poziotinib** | **HER2** | **TK** | **19439.7** | **63934.7** | **83374.5** |
| **418** | **A8215** | **PP 1** | **Src** | **TK** | **19934.9** | **63796.3** | **83731.2** |
| **419** | **A3132** | **A 77-01** | **TGF-?R1(ALK5)** | **TKL** | **63808.0** | **20371.9** | **84180.0** |
| **420** | **B3553** | **GS-9973** | **Spleen Tyrosine Kinase (Syk)** | **TK** | **32878.5** | **51541.3** | **84419.8** |
| **421** | **B4898** | **Bikinin** | **GSK-3** | **CMGC** | **39833.6** | **44813.6** | **84647.2** |
| **422** | **C4882** | **1,2-Dilauroyl-sn-glycerol** | **PKC** | **AGC** | **32229.3** | **52450.9** | **84680.2** |
| **423** | **B3033** | **Bay 11-7085** | **I?B/IKK** | **Other** | **63723.6** | **21705.2** | **85428.7** |
| **424** | **N1345** | **Obacunone** | **Others** | **Other** | **22534.9** | **63087.7** | **85622.6** |
| **425** | **A3575** | **LY2835219 free base** | **Cyclin-Dependent Kinases** | **CMGC** | **59180.8** | **26611.9** | **85792.7** |
| **426** | **A4605** | **KU 55933** | **ATM/ATR** | **Atypical** | **33867.0** | **52307.6** | **86174.6** |
| **427** | **A1947** | **MEK162 (ARRY-162, ARRY-438162)** | **MEK1/2** | **STE** | **32104.0** | **54835.5** | **86939.5** |
| **428** | **B4800** | **Defactinib** | **FAK** | **TK** | **31683.7** | **56452.3** | **88136.0** |
| **429** | **A4112** | **Barasertib (AZD1152-HQPA)** | **Aurora Kinase** | **Other** | **48647.4** | **41038.0** | **89685.3** |
| **430** | **A5760** | **KRN 633** | **VEGFR** | **TK** | **55097.1** | **35164.2** | **90261.3** |
| **431** | **A8374** | **AZD8330** | **MEK1/2** | **STE** | **29149.3** | **63631.5** | **92780.8** |
| **432** | **A3965** | **BI 2536** | **PLK** | **Other** | **34739.4** | **58236.8** | **92976.2** |
| **433** | **C5545** | **MK2 Inhibitor IV** | **P2X purinergic receptor** | **Other** | **42817.4** | **50349.5** | **93166.9** |
| **434** | **B4906** | **PF-06447475** | **LRRK2** | **TKL** | **53174.2** | **40604.3** | **93778.5** |
| **435** | **A8216** | **PP 2 (AG 1879)** | **Src** | **TK** | **53589.5** | **40414.9** | **94004.3** |
| **436** | **A3717** | **PF-543** | **S1P receptor** | **Other** | **28127.6** | **66805.4** | **94933.0** |
| **437** | **B1402** | **GZD824** | **Bcr-Abl** | **TK** | **64275.0** | **33439.8** | **97714.8** |
| **438** | **A5880** | **R406 (free base)** | **Spleen Tyrosine Kinase (Syk)** | **TK** | **42129.8** | **55638.2** | **97768.1** |
| **439** | **A8395** | **CHIR-98014** | **GSK-3** | **CMGC** | **63043.9** | **37241.6** | **100285.6** |
| **440** | **B5624** | **STF 083010** | **AChE** | **Other** | **54363.2** | **46405.9** | **100769.1** |
| **441** | **B5952** | **LFM-A13** | **BTK** | **TK** | **60025.3** | **42758.8** | **102784.1** |
| **442** | **A4124** | **TAK-901** | **Aurora Kinase** | **Other** | **42486.6** | **61332.6** | **103819.2** |
| **443** | **B5854** | **Pexidartinib (PLX3397)** | **CSF-1R** | **TK** | **64762.3** | **40586.0** | **105348.3** |
| **444** | **A8370** | **Axitinib (AG 013736)** | **VEGFR** | **TK** | **42600.6** | **65444.2** | **108044.8** |
| **445** | **A3519** | **JNK-IN-7** | **JNK** | **CMGC** | **66910.1** | **45982.8** | **112893.0** |
| **446** | **B1974** | **Methylthiouracil** | **Others** | **Other** | **61524.3** | **55636.1** | **117160.4** |
| **447** | **A4135** | **Tofacitinib (CP-690550) Citrate** | **JAK** | **TK** | **57107.2** | **61115.5** | **118222.7** |
| **448** | **A4541** | **Sal 003** | **Protein Ser/Thr Phosphatases** | **CMGC** | **58234.6** | **61292.8** | **119527.4** |
| **449** | **A8448** | **INCB28060** | **c-MET** | **TK** | **59276.3** | **63091.3** | **122367.6** |
| **450** | **B8016** | **UNC2025** | **FLT3** | **TK** | **55553.7** | **67297.4** | **122851.1** |
